# Structural and mechanistic basis of the EMC-dependent biogenesis of distinct transmembrane clients

**DOI:** 10.1101/2020.09.02.280008

**Authors:** Lakshmi E. Miller-Vedam, Bastian Bräuning, Katerina D. Popova, Nicole T. Schirle Oakdale, Jessica L. Bonnar, Jesuraj Rajan Prabu, Elizabeth A. Boydston, Natalia Sevillano, Matthew J. Shurtleff, Robert M. Stroud, Charles S. Craik, Brenda A. Schulman, Adam Frost, Jonathan S. Weissman

## Abstract

Membrane protein biogenesis in the endoplasmic reticulum (ER) is complex and failure-prone. The ER membrane protein complex (EMC), comprising eight conserved subunits, has emerged as a central player in this process. Yet, we have limited understanding of how EMC enables insertion and integrity of diverse clients, from tail-anchored to polytopic transmembrane proteins. Here, yeast and human EMC cryo-EM structures reveal conserved intricate assemblies and human-specific features associated with pathologies. Structure-based functional studies revealed at least two separable EMC activities, as an insertase regulating tail-anchored protein levels and as a polytopic membrane protein holdase chaperone. These depend on mechanistically coupled yet spatially distinct regions including two lipid-accessible membrane cavities which confer client-specific regulation, and a novel, non-insertase EMC function mediated by the EMC lumenal domain. Our studies illuminate the structural and mechanistic basis of EMC’s multifunctionality and point to its role in differentially regulating the biogenesis of distinct client protein classes.

## Introduction

Integral membrane proteins serve diverse and critical cellular roles, including signal transduction, lipid biosynthesis, adhesion, and transport of molecules across the bilayer. In eukaryotic cells, the endoplasmic reticulum (ER) serves as the primary site of integral membrane protein synthesis, targeting (co-or post-translationally), insertion, folding and quality control (Ellgaard 2016; Costa, 2018). However, the features of membrane-spanning regions (e.g. low hydrophobicity, charged residues, non-optimal lengths, lipid- and ion-binding sites and hairpins or kinked transmembrane helices) that mediate important functions pose particular challenges for transmembrane protein biosynthesis and folding. Consequently, membrane protein biogenesis is prone to failure, and this can lead to cellular stress and disease (Marinko, 2019). Thus, it is important to understand the cellular factors that facilitate proper membrane protein biogenesis for such challenging clients.

The ER membrane protein complex (EMC) has emerged as a conserved player in the process of membrane protein biogenesis. It was first identified in *Saccharomyces cerevisiae,* as an abundant and stable multi-protein membrane complex whose disruption results in stress mirroring that caused by misfolded membrane proteins (Jonikas, 2009). Loss of the EMC in mammalian cells is associated with failed biogenesis and degradation of a subset of membrane proteins (Christianson, 2012). Accordingly, the EMC has been implicated in several mechanistically distinct steps of membrane protein biogenesis, stabilization, and quality control (Bircham, 2011; Richard, 2013; Satoh, 2015; Savidis, 2016; Shurtleff and Ithzak, 2018; Volkmar, 2018; Tian, 2019).

One well-established EMC function is as an insertase for terminal transmembrane helices. EMC’s insertase function has been demonstrated for two classes of clients: low hydrophobicity tail-anchored proteins (i.e. those that contain C-terminal membrane anchors) and a subset of polytopic transmembrane proteins in which the first helix is inserted with the N-terminus in the lumen (Guna, 2018; Chitwood, 2018). However, many studies indicate EMC functions beyond initial insertion of N- or C-terminal helices. The EMC has been implicated in the biogenesis and stability of many membrane protein classes that do not require a terminal transmembrane insertase (Bircham, 2011; Louie, 2012; Richard, 2013; Shurtleff and Ithzak, 2018; Coelho, 2019; Luo, 2002; Volkmar, 2018; Talbot, 2019; Petkovic, 2020). Recent studies have shown that the EMC is required for stability of internal transmembrane helices of human and viral multi-pass membrane proteins (Hiramatsu, 2019; Lin 2019; Ngo, 2019; Coelho, 2019; Xiong, 2020). Additionally, the human EMC (hEMC) physically interacts with the NS4A-B region of the Dengue Virus polyprotein following Sec61-dependent translocation and signal peptidase cleavage, suggesting roles in post-translational stabilization of polytopic membrane proteins (Ngo, 2019; Lin 2019). Similarly, the *S. cerevisiae* EMC (yEMC) co-immunoprecipitated with full-length polytopic transmembrane clients, including Pma1p (Luo, 2002), Mrh1p, and Fks1p (Shurtleff and Ithzak, 2018). In addition to varying types of transmembrane protein clients, the EMC also associates with a range of regulatory factors, including many general and substrate-specific chaperones in the cytoplasm and in the ER lumen (Bagchi, 2016; Coelho, 2019; Kudze, 2018; Richard, 2013; Shurtleff and Ithzak, 2018).

The complex architecture of the EMC provides additional support for multifunctionality in membrane protein biogenesis. The EMC is an eight (yeast) or nine (mammalian) component, 248-284 kDa complex with considerable mass in the ER lumen, membrane and cytosol. The cytoplasmic domain contains conserved tetratricopeptide repeats (TPR) repeats in EMC2, and the human complex accommodates an additional subunit, EMC8/9, whose function is not yet understood. The ER lumenal domain in yeast does not contain an N-terminal EMC1 expansion seen in hEMC. Notably, the ER lumenal domain has been linked to a number of disease-associated phenotypes (Junes-Gill, 2010; Probert, 2015; Harel, 2016; Abu-Safieh, 2012; Diamantopoulou, 2017; Marquez, 2020), and presents the possibility of additional functions for the human lumenal domain. One EMC subunit (EMC3) shares limited sequence homology with a family of insertases that are evolutionarily related to the bacterial insertase YidC (Samuelson, 2000; Kumazaki, 2014; Borowska, 2015; Anghel, 2017), perhaps explaining the insertase function of the complex. Nonetheless, the elaboration of the EMC compared to other known transmembrane domain (TMD) insertases and a diverse client range suggest additional functionality that has so far eluded mechanistic explanation.

Here, we determined high-resolution cryo-EM structures of yeast and human EMC and characterized the phenotypes of three distinct classes of EMC clients associated with a series of structure-based EMC mutants. Both yEMC and hEMC structures reveal a path for transmembrane helix insertion from the cytoplasm into the membrane via a conserved cavity. Our structures and mutants also revealed a second lipid-filled cavity with regions of importance for all three client types probed. Analysis of human disease mutations in EMC1 and our structure-informed mutations enabled us to decouple the EMC insertase function from non-insertase functions and reveal a potential role of the EMC in differentially controlling the biogenesis of distinct classes of client proteins. These structure-function studies collectively establish that the EMC adopts a modular architecture enabling its diverse functions in membrane protein biogenesis.

## Results

### Overview of strategy to comprehensively reveal EMC structure and function

To comprehensively dissect both conserved and species-specific functions of the EMC, we developed approaches to produce EMC for structure determination and broad mutational analysis (**Figure 1**). We developed systems to produce robust quantities of pure intact yEMC and hEMC to determine structures for the two organisms in which different facets of EMC function have been described in detail (Jonikas, 2009; Christianson, 2012; Guna, 2018; Shurtleff and Ithzak, 2018). Parallel efforts converged on an approach involving FLAG affinity-tagging of the EMC5 C-terminus, which was performed for endogenous yEMC and recombinant hEMC in human embryonic kidney (HEK) cells (**Figure 1 – figure supplement 1-2**).

**Figure 1:**
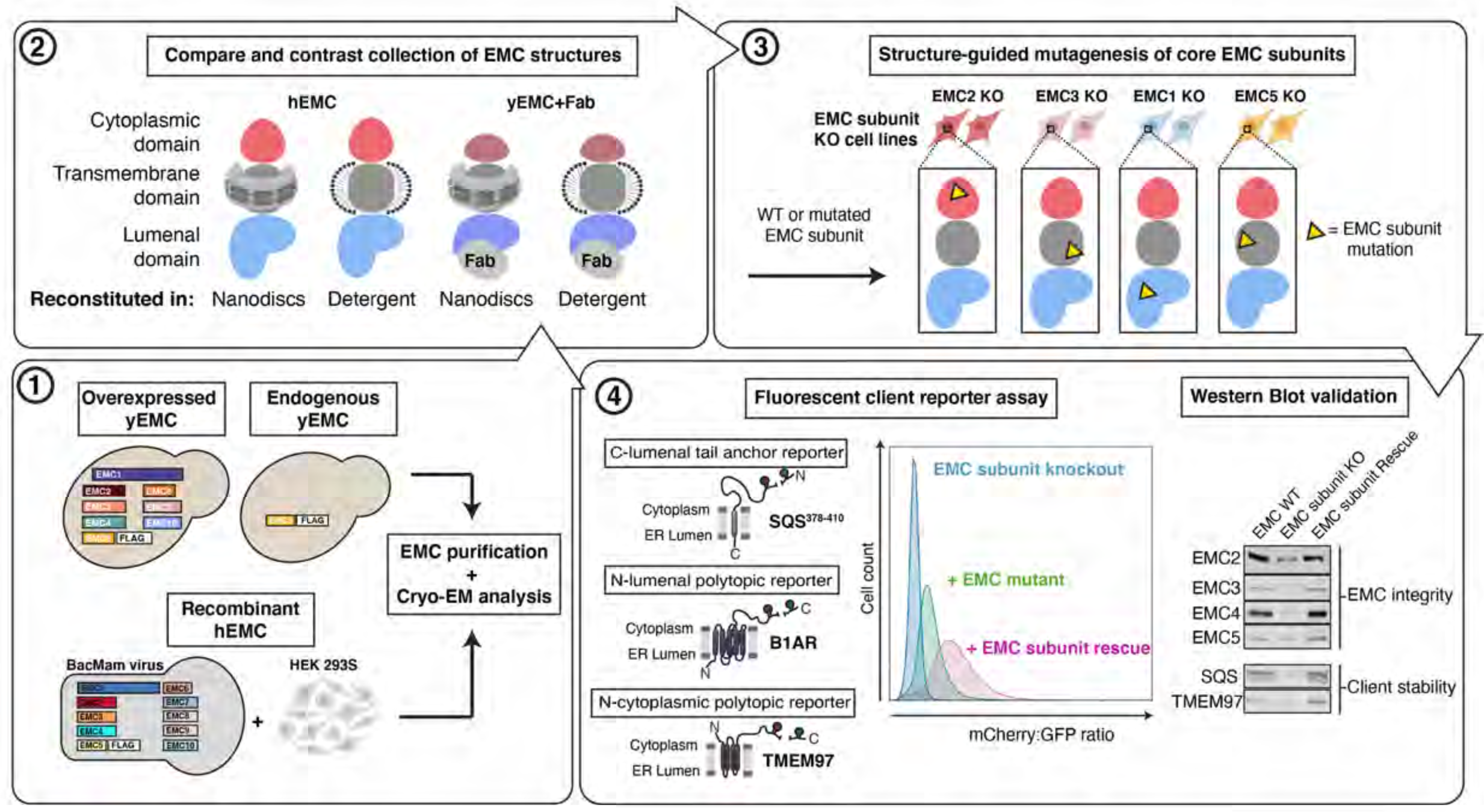
Experimental strategy for the dissection of EMC function. Schematic representation of the combined structural and mutational approach to dissect EMC function. (1) yEMC was expressed either by overexpression of all subunits together with Flag-tagged EMC5 or by endogenous expression from a genomic Flag-tagged EMC5 strain. For hEMC, all subunits were overexpressed together with Flag-tagged EMC5 via a single recombinant BacMam virus. EMC was purified by column chromatography and subjected to cryo-EM analysis. (2) The obtained collection of cryo-EM structures of yEMC and hEMC in nanodiscs or detergent were compared to identify similarities and differences. (3) Structure-guided mutagenesis was performed across four core EMC subunits: EMC1, EMC2, EMC3, and EMC5. (4) Each EMC subunit knock-out (KO) cell line was individually transduced with three different fluorescent client reporters: SQS^378-410^, full length B1AR, and full length TMEM97. Mutant EMC subunits were introduced into KO cell lines missing the corresponding EMC subunit in each of the three fluorescent EMC client reporter cell lines. EMC client stability in each mutant EMC subunit cell line was assessed by fluorescent cell sorting and quantifying the mCherry-to-GFP ratio. Western blotting was performed for each mutant-transduced cell line to assess EMC integrity (by immunoblotting for core EMC subunits) as well as client stability (by immunoblotting for EMC clients) and compared against both wild-type (WT) and KO cell lines.

**Video 1.**
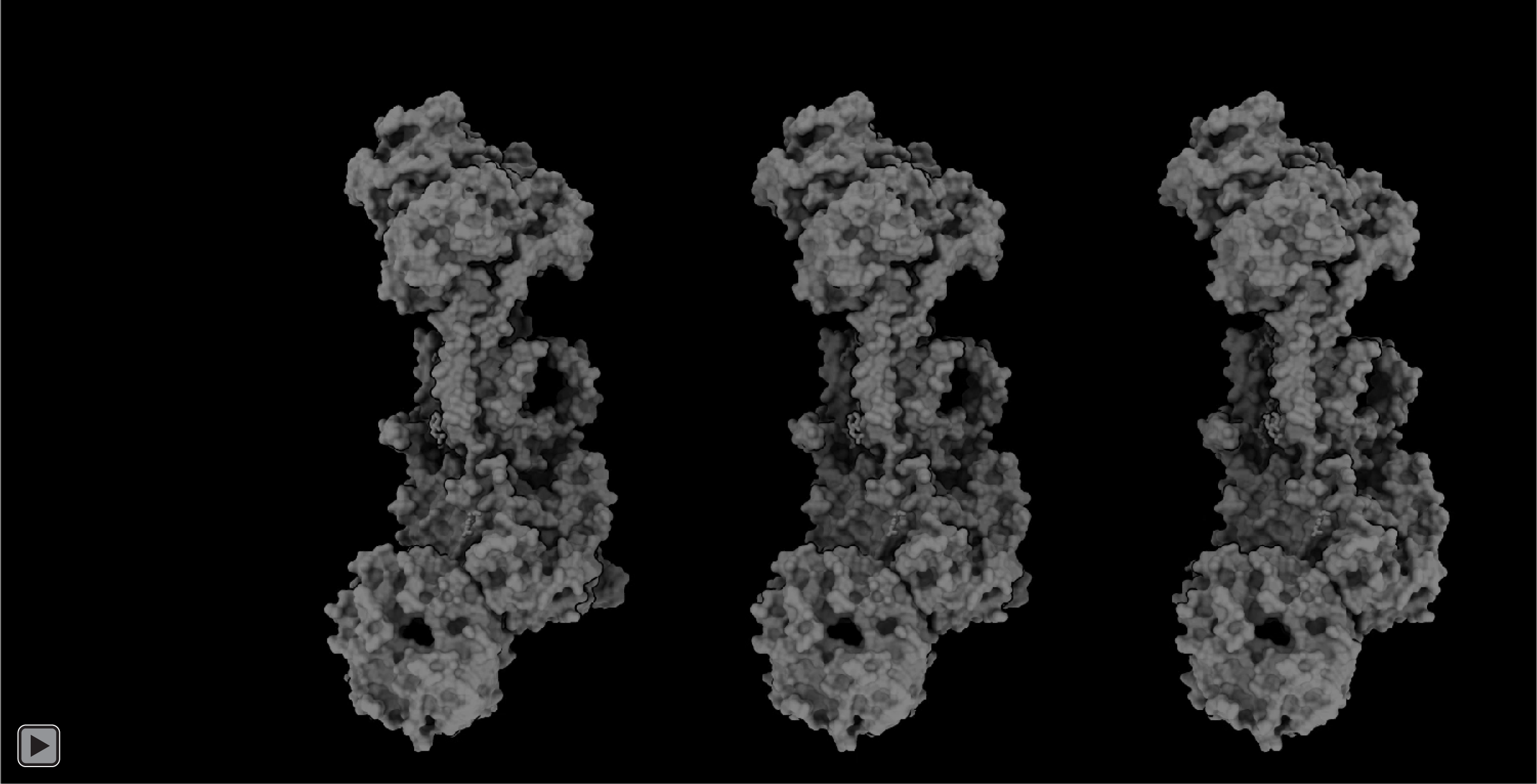
hEMC mutagenesis displayed on hEMC structures. Three identical copies of hEMC in nanodisc are displayed here. Subsequent labeling and animation color mutated residues by flow cytometric measure of reporter abundance, grouped into three categories: increased reporter levels (mCherry>GFP signal, colored magenta), wild type levels (mCherry signal is close to GFP signal, colored silver), and decreased reporter levels (mCherry<GFP signal, colored gold). Left hEMC structure displays phenotypes for the C-lumenal tail anchor reporter (GFP-P2A-mCherry-SQS^378-410^-opsin). Middle hEMC structure displays phenotypes for the N-lumenal polytopic reporter (B1AR-mCherry-P2A-GFP). Right hEMC structure displays phenotypes for the N-cytoplasmic polytopic reporter (TMEM97-mCherry-P2A-GFP). Mild phenotypes are displayed as wild type levels

In parallel, to enable testing of hypotheses based on structures, we created a suite of human (K562) knockout cell lines deleted for individual EMC subunits - EMC1 (lumen), EMC2 (cytoplasm), EMC3 and EMC5 (transmembrane) - and a series of reporters of EMC-dependent transmembrane protein biogenesis (**Figure 1 – figure supplement 3**).

Reintroduction of the wildtype EMC subunits in the respective knockout cells fully rescued the knockout phenotype (**Figure 1 – figure supplement 4-5**). This allowed for introduction of structure-based mutations in EMC subunits into the respective knockout cells to determine features supporting biogenesis of fluorescently-tagged versions of three different types of EMC clients: the transmembrane domain of a C-terminal tail-anchored transmembrane protein (squalene synthase, SQS^378-410^) (Guna, 2018), a polytopic transmembrane protein that depends on the EMC N-terminal insertase activity (Beta 1 adrenergic receptor, B1AR) (Chitwood, 2018), and a polytopic transmembrane protein (Sigma intracellular receptor 2, TMEM97) whose biogenesis requires the Sec61 translocon but does require a terminal helix insertase (**Figure 1 – figure supplement 3-7**). Monitoring the effect of an EMC mutation on fluorescent reporter levels provided a quantitative measure of its impact on EMC-dependent biogenesis of each class of client protein. We subsequently mapped these mutations onto the structure grouped by reporter phenotype (**Video 1**). Western blots against the endogenous EMC subunits allowed us to control for mutational effects on the production and stability of the EMC complex itself (**Figure 1 – figure supplement 4-5**). This strategy thus distinguishes effects resulting from a global disruption of the EMC complex from those caused be specific disruption of one or more EMC functions.

### The EMC is an intricate molecular machine spanning the ER membrane and exhibits a conserved core architecture

We determined structures of yEMC and hEMC — all showing overall compositional similarity, yet regional conformational differences between the yeast and human complexes (**Figure 2**). We obtained reconstructions of yEMC bound to an antigen binding fragment (Fab) and hEMC reconstituted both in detergent micelles and lipid nanodiscs, with the latter strategy yielding the most isotropic and highest resolution data. For yEMC+FabDH4 and hEMC, the global map resolutions reached 3.2 Å and 3.4 Å, respectively (**Figure 2 – figure supplement 1 - 4**). The cryo-EM maps allowed for *de novo* model building of both human and yeast complexes (**Figure 2 – figure supplement 5, 6**). We note that our maps and models are consistent with recent cryo-EM data from yeast EMC (Bai, 2020), human EMC (O’Donnell, Phillips, and Yagita, 2020; Pleiner, Tomaleri, and Januszyk, 2020), and a crystal structure of human EMC2-EMC9 (O’Donnell, Phillips, and Yagita, 2020) published while we were in the final stages of manuscript preparation. As described in the following sections, our multiple EMC structures enable a broad survey of its conserved architecture, with variations between the structures pointing to conformational and compositional differences (**Figure 2 – figure supplement 7**).

**Figure 2:**
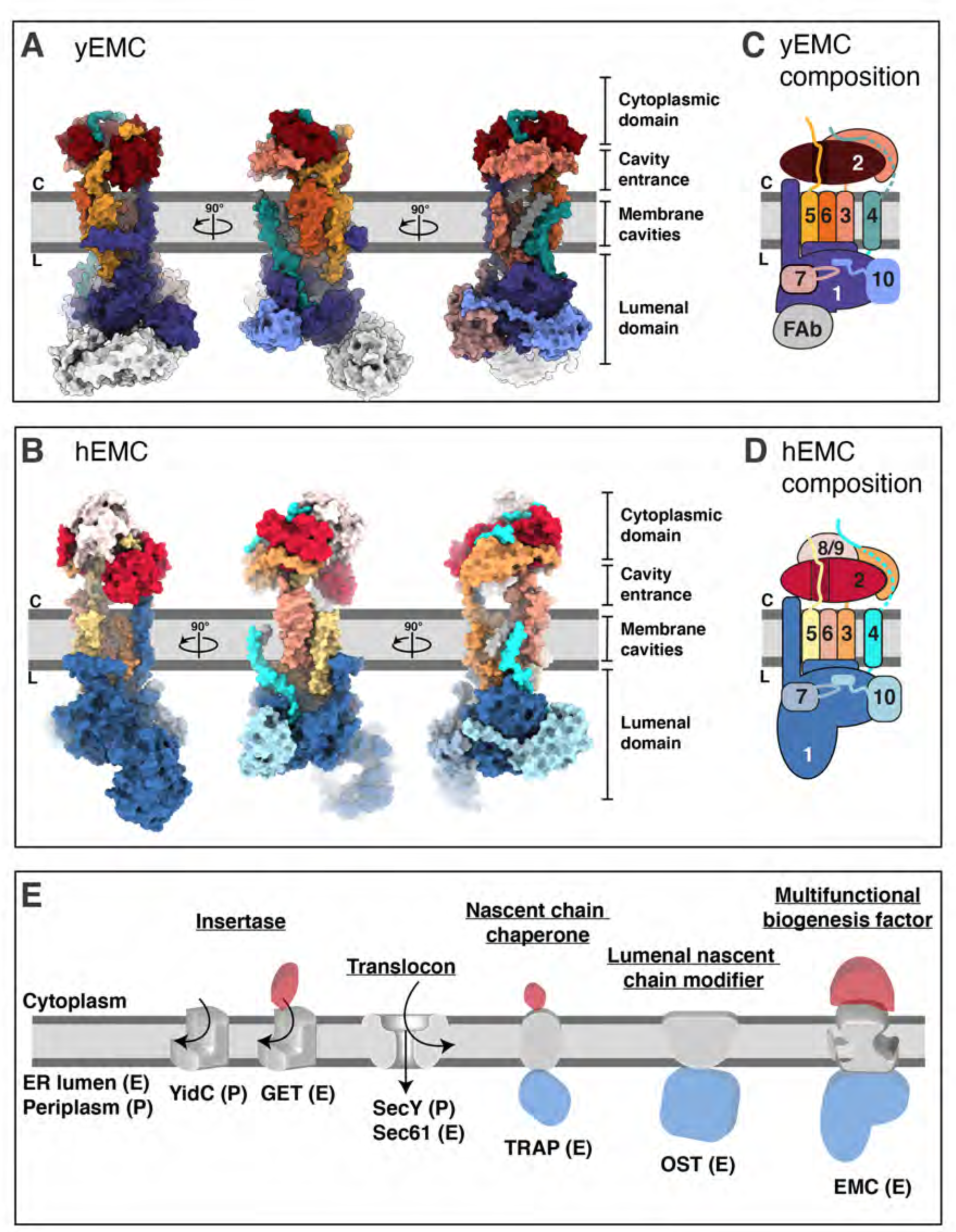
Overall structures of yeast and human EMC. A) Cryo-EM structure of yEMC in nanodiscs. Three orthogonal views of the yEMC cryo-EM structure shown as surface rendering. Grey bars delineate the approximate ER membrane boundaries with the cytoplasmic (C) and lumenal (L) sides indicated. The FAb molecule bound to the yEMC1 lumenal domain is colored in grey. B) Cryo-EM structure of hEMC in nanodiscs. Labeling as in A). C) Subunit composition and color scheme of yEMC used throughout the manuscript. Dotted line indicates a portion of EMC4 unresolved in the cryo-EM map and left unmodeled. D) Subunit composition and color scheme of hEMC used throughout the manuscript. E) Schematic depiction and comparison of the EMC architecture to known transmembrane protein biogenesis factors in the ER and the bacterial plasma membrane. Cytoplasmic, transmembrane and lumenal domains are depicted as cartoons colored red, grey and blue, respectively. E, eukaryotic; P, prokaryotic.

The EMC comprises cytoplasmic, transmembrane and lumenal domains arranged similarly for yeast and human, despite significant evolutionary separation (**Figure 2A-B**). For both species, subunits encompassing EMC2 to EMC7 form an interconnected core complex, while there is additional density capping both the cytoplasmic and lumenal domains of hEMC, occupied by an EMC8/9 and an EMC1 N-terminal expansion, respectively (**Figure 2C-D**).

EMC8 and EMC9 are paralogs of each other, which have not been identified in yeast (Wideman, 2015). We modeled and depict only EMC8 for clarity, but due to the 44% sequence identity with EMC9 and both being present in the recombinant system we refer to this as EMC8/9. The large EMC1 insertion in hEMC constitutes the majority of a membrane distal beta-propeller domain protruding into the lumen, a feature missing from *S. cerevisiae*. Compared to other ER-resident proteins (Suloway, 2009; Pfeffer, 2017; Ramírez, 2019), the arrangement of domains of the EMC is unusual with the transmembrane domain connecting prominent cytoplasmic and lumenal domains (**Figure 2E)**. On a global level, the structure suggests complexities beyond those of some other ER machineries fulfilling select functions in transmembrane protein biogenesis.

### The cytoplasmic domain provides a platform for protein-protein interactions

The exterior interface of the cytoplasmic domain is formed by EMC2, EMC3, EMC4, and parts of EMC8/9 (in human), while parts of EMC5, EMC2, and EMC8/9 are shielded from the cytoplasm (**Figure 3A-B; Figure 2 – figure supplement 7**). The helical fold of EMC2 constitutes the central organizer of this platform, established by five or six TPR motifs in human versus yeast, respectively (**Figure 3C**). TPR domains are commonly found mediating protein-protein interactions, and are present in numerous well-characterized chaperone-protein and other interaction networks (Blatch, 1999; Scheufler, 2000; Schlegel, 2007; Assimon, 2015; Krysztofinska, 2017; Graham, 2019). Yeast EMC2 features a more curved helical arrangement with N-and C-terminal domains in closer proximity to each other than seen in human EMC2. Notably, the canonical peptide-binding TPR groove is occupied by the partially helical C-terminus of EMC5, which forms a large interaction surface with EMC2. To test the functional roles of this interaction, we mutated three residues within the EMC2 TPR motif (hEMC2^K125E + R126D + K127E^) or a single EMC5 residue buried in the TPR binding groove (hEMC5^F90A^). The mutations on both sides of the interface decreased EMC integrity, with a modest (for the single point mutation) and significant (for the multi-residue mutation) reduction in the levels of several EMC subunits (**Figure 3C, Figure 3 – figure supplement 1-2, Figure 1 – figure supplement 4-5**). This suggests that this interface might be critical for EMC complex assembly rather than activity (e.g. client binding).

**Figure 3:**
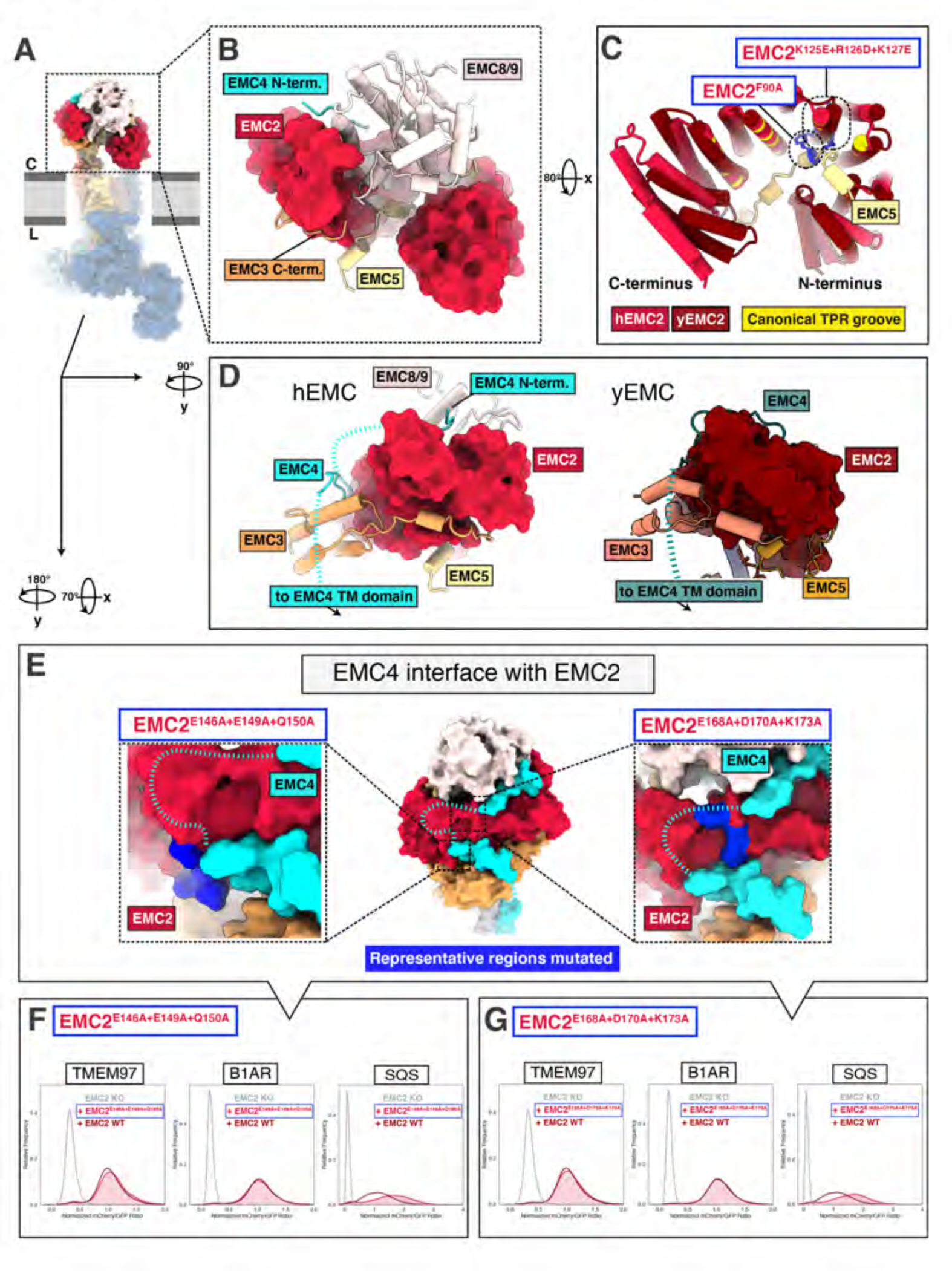
The EMC cytoplasmic domain contains conserved functional interfaces and may engage C-tail anchored clients directly. A) Position of the hEMC cytoplasmic domain relative to the membrane and the rest of the complex. Shown is the surface rendered hEMC structure reconstituted in nanodiscs. B) EMC2 nucleates a protein-protein interaction hub in the cytoplasm. Zoomed-in view of the cytoplasmic domain from A). EMC2 is shown as surface rendering while interacting EMC subunits are shown as cartoon cylinders. C) EMC2 forms a TPR domain which binds EMC5. Overlaid are hEMC2 (red) and yEMC2 (dark red), illustrating the more tightly wound yEMC2 TPR solenoid. Canonical residues lining the concave TPR interior are colored in yellow stripes on the dark red for hEMC2. Two mutants, one in EMC5 and three in EMC2, are colored in blue, and show destabilizing phenotypes for EMC integrity. D) A cytoplasmic cap structure involving EMC4 is conserved in yEMC and hEMC. Shown is a side-by-side comparison between the cytoplasmic domains of hEMC (left) and yEMC (right), highlighting the similar path EMC4 takes from the cytoplasmic domain towards the transmembrane domain. While an interaction surface between EMC8/9 and the EMC4 N-terminus is absent in yeast, yEMC4 binds at the top of the EMC2 TPR domain and assumes as similar position across the EMC3 cytoplasmic domain at the cytoplasm-membrane interface. E) Location of two EMC2-EMC4 interfaces targeted for mutagenesis. hEMC is shown as surface rendering with the mutated EMC2 regions highlighted in blue. F) Fluorescent client reporter stability assay for TMEM97 (N-cytoplasmic polytopic client), B1AR (N-lumenal polytopic client) and SQS^378-410^ (C-lumenal tail-anchored client) in EMC2 KO cells expressing mutant hEMC2^E146A+E149A+Q150A^ (shaded) or WT hEMC2 rescue (unshaded). G) Fluorescent client reporter stability assay, as in F, for the hEMC2^E168A+D170A+K173A^ mutant.

The multi-protein cytoplasmic cap has distinct elements between hEMC and yEMC. Capping the cytoplasmic domain in hEMC is EMC8/9 (**Figure 3 – figure supplement 3**), the functional roles of this cap-like structure are not yet clear. A groove on EMC8/9 cradles an N-terminal peptide of EMC4, which proceeds into the EMC4 segment that traverses over EMC2 and the three-helix bundle of EMC3 (**Figure 3D**). Though yEMC lacks EMC8/9, yeast EMC4 follows a similar binding trajectory along cytoplasmic EMC2 and EMC3 surfaces. We mutated two patches of EMC2 residues in proximity to the EMC4 cytoplasmic region (hEMC2^E146A+E149A+Q150A^, hEMC2^E168A+D170A+K173A^), which led to a modest accumulation of the tail-anchored client (SQS^378-410^) but did not affect polytopic client abundance or decrease abundance of EMC subunits (**Figure 3E-G, Figure 1 – figure supplement 4-5**). Several mutants across the cytoplasmic domain showed similar phenotypes, supporting a key role in tail anchor protein biogenesis (**Figure 3 – figure supplement 1-2**).

### Two distinct cavities are present in the transmembrane domain

The transmembrane core of EMC is predicted to include contributions from each subunit except for EMC2 and, in humans, EMC8/9 (**Figure 2C-D**). The EMC presents two distinct and structurally conserved cavities on opposite sides of the transmembrane core that differ in size, shape, subunit compositions and apparent function (**Figure 4A-B**). One cavity, which we refer to as the lipid-filled cavity, appears contiguous with the ER lipid environment (**Figure 4A**). The second cavity, which we refer to as the gated cavity, appears to open towards the cytoplasm in our structures and is more occluded by a transmembrane helix gate from the lipid environment (**Figure 4B**). Notable structural hallmarks present in both species include a superimposable core of nine transmembrane helices, a set of flexible gate helices, and an amphipathic EMC1 brace helix (**Figure 4C**).

**Figure 4:**
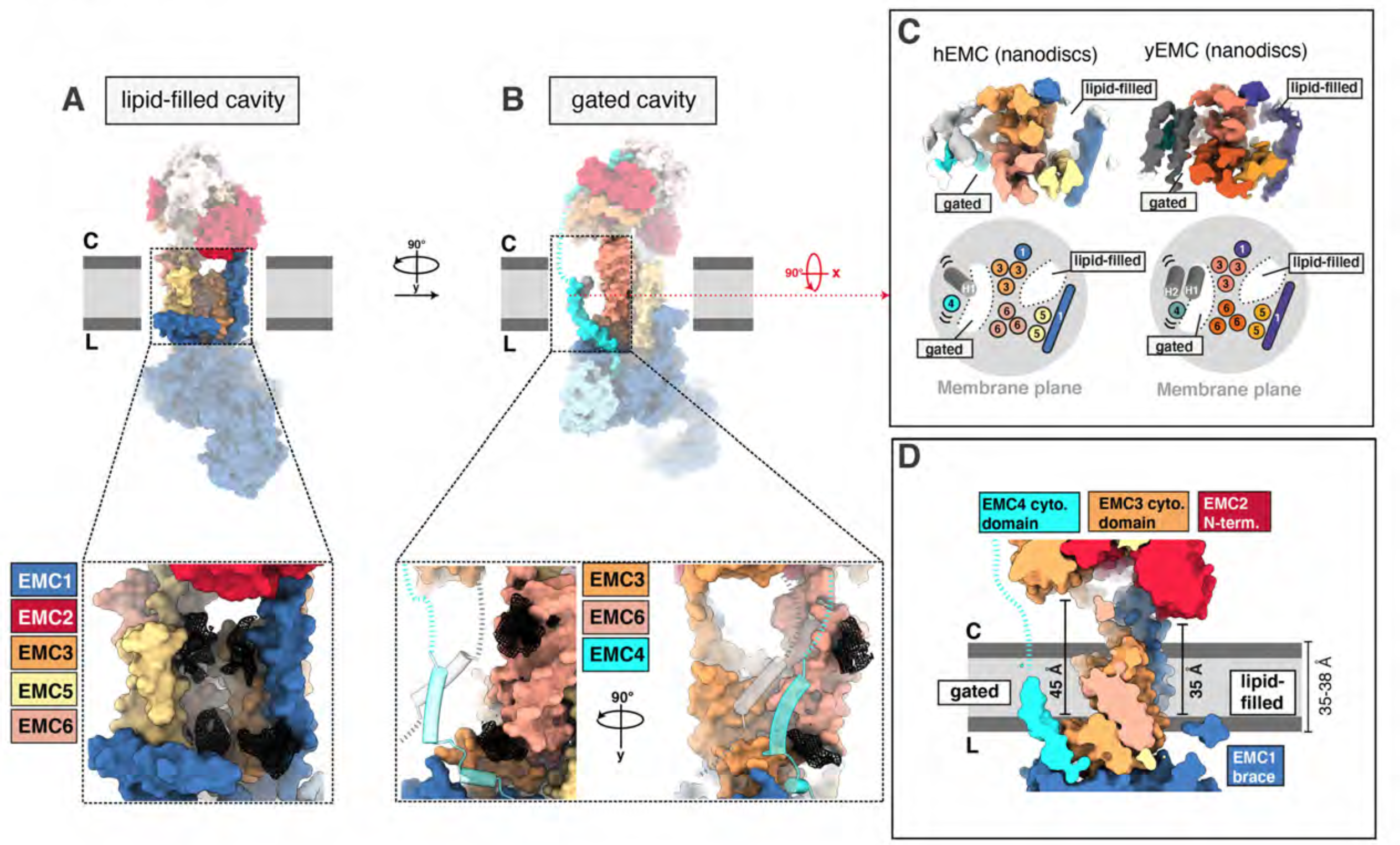
The EMC houses two transmembrane cavities with conserved core structures and distinct accessibilities. A) Location and composition of the lipid-filled cavity. A zoom-in view on the cavity is shown below, which is composed of EMC1, EMC3, EMC5 and EMC6. Resolved lipid densities from the cryo-EM map of hEMC in POPC nanodiscs are shown as black mesh zoned within 3 Å of modeled POPC molecules. B) Location and composition of the gated cavity. Two orthogonal zoom-in views of the cavity are shown below, which is composed of EMC3 and EMC6. A transmembrane gate opposite the cavity wall is depicted as transparent cartoon cylinders and has contributions from the C-terminal EMC4 transmembrane helix along with up to two additional, unassigned helices. Resolved lipid densities are shown as in A). C) The dual-cavity architecture of the EMC transmembrane domain is conserved between yEMC and hEMC. Unsharpened cryo-EM maps of hEMC and yEMC in nanodiscs (top) are shown along with corresponding schematic representations of the spatial organization of all transmembrane helices (bottom). The gate helices of the gated cavity represent the region of highest conformational heterogeneity across our collection of EMC structures. D) The two EMC transmembrane cavities feature distinct accessibilities. Shown is a central slice through the surface rendered hEMC nanodisc structure with the two membrane cavities on opposite sides. Measuring from the lumenal to the cytoplasmic side, gated and lipid-filled cavities measure 45 Å and 35 Å across, respectively. This suggests that the gated cavity has accessibility from the cytoplasm while the lipid-filled cavity does not.

### The gated cavity serves as a conduit for terminal helix insertion

Evaluating potential client paths from the cytoplasm into the transmembrane domain revealed a cavernous opening at the membrane-cytoplasmic interface of the gated cavity, wide enough to allow passage of a client helix, and tapering towards the lumen (**Figure 4D)**. Consistent with its potential role as a cytoplasmic conduit into the EMC, the EMC3 portion of the cytoplasmic domain, which delineates this opening, sits approximately 45 Å from the lumenal side of the gated cavity. This dimension exceeds the thickness of the ER membrane (Mitra, 2004; Heberle, 2020; Cornell, 2020) (**Figure 4D**). This cavity is lined primarily by EMC3, EMC4 and EMC6 (**Figure 5A**). Simulating the dimension of the first transmembrane helix of a known terminal insertase-client (B1AR, Chitwood, 2018) suggests that there is sufficient space for a client helix even in the client-free state of the EMC (**Figure 5B**). The gated cavity is hydrophilic on the cytoplasmic side and becomes increasingly hydrophobic towards the lumenal side (**Figure 5C**).

**Figure 5:**
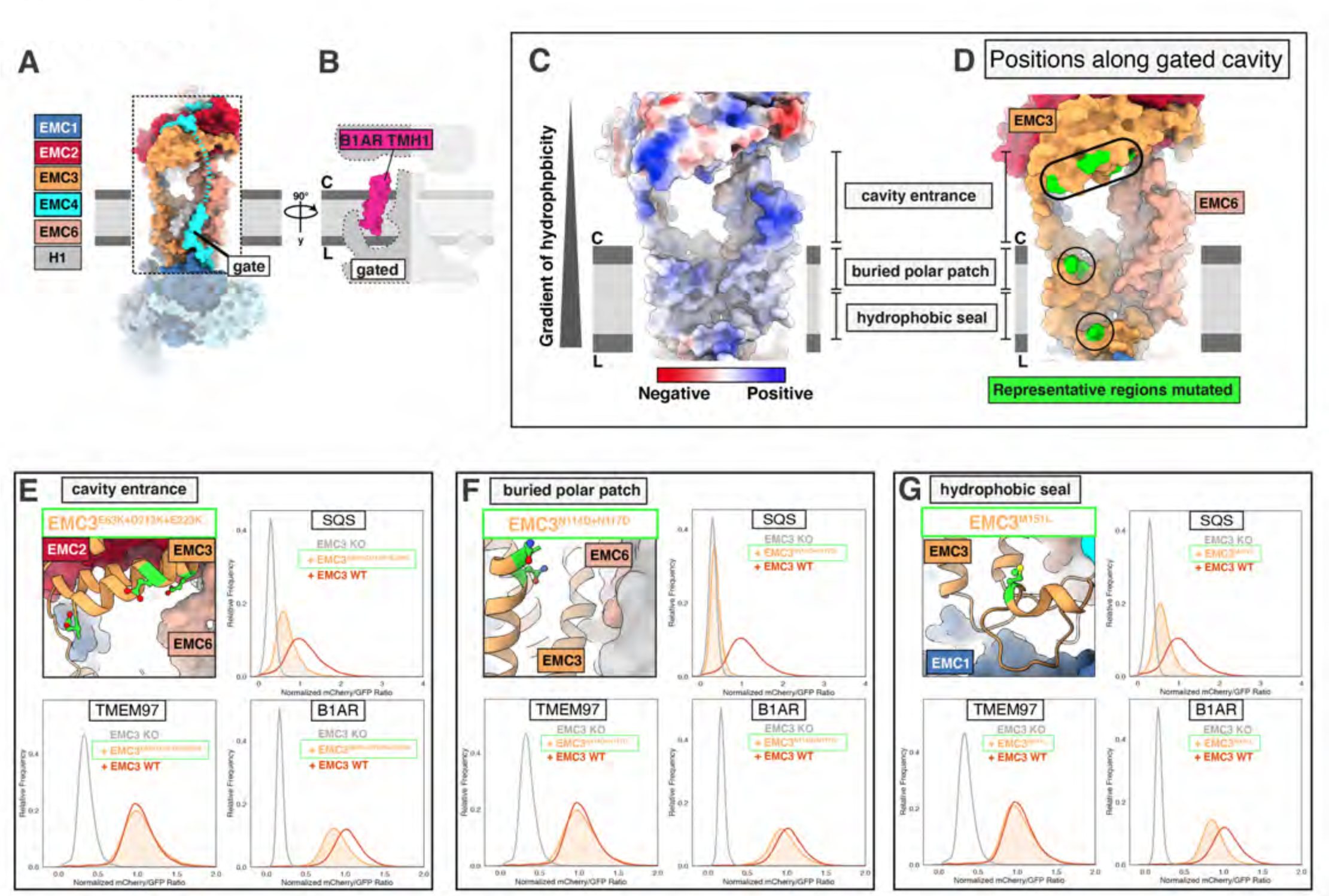
EMC houses an insertase module centered on EMC3 in the gated membrane cavity. A) A transmembrane gate anchored in the cytosol and the lumen is a structural hallmark of the EMC gated cavity. Shown is a surface rendering of the hEMC model in lipid nanodiscs with an unresolved EMC4 connection between the cytoplasm and the membrane depicted as a dashed line. An unassigned helix of the gate is shown in gray (H1). B) The gated cavity in the hEMC nanodisc structure has sufficient space to accommodate a client transmembrane helix. The space-filling model of the first transmembrane helix of B1AR (B1AR TMH1) is shown placed inside an outline of the gated EMC cavity. C) A hydrophobic gradient characterizes the surface of the EMC gated cavity from the cytoplasmic to the lumenal side. Gate helices have been omitted for clarity. The surface of the hEMC nanodisc structure is colored by electrostatic surface potential ranging from -15 (red) to +15 (blue) kcal/(mol·*e*). D) Distinct EMC3 regions along the gated cavity hydrophobic gradient targeted for mutagenesis. Mutated residues are colored in lime. E) Fluorescent client reporter stability assay for the EMC3 cavity entrance mutant, hEMC3^E63K+D213K+E223K^. A zoom-in on the mutated residues (colored lime) is shown in the top-right panel. F) As in E) for the EMC3 buried polar patch mutant, hEMC3^N114D+N117D^. G) As in E) for the EMC3 hydrophobic seal mutant, hEMC3^M151L^.

The entrance into the gated cavity interior (**Figure 5A**) is formed primarily by the EMC3 cytoplasmic domain. To test its function, charge swap mutations were introduced along the rim of this opening (hEMC3^E63K + D213K + E223K^, hEMC3^R59E + R62E + K216E^) (**Figure 5D, Figure 5 – figure supplement 2**). These mutants resulted in loss of the tail-anchored client (SQS^378-410^) and partial loss of the N-terminal insertase dependent polytopic client (B1AR), reflecting a failure to support insertase activity. These mutants had no appreciable effect on the abundance of the polytopic transmembrane client (TMEM97) reporter (**Figure 5E, Figure 5 – figure supplement 1-2**). A similar phenotype was observed with alanine substitutions for a pair of lysines at the periphery of this cytoplasmic rim (hEMC3^K42A + K43A^) (**Figure 5 – figure supplement 2**).

Having identified a functionally important entry route for terminal helix insertase clients, we next considered potential surfaces inside the cavity that might accommodate a client helix. A polar patch close to the membrane interior of this cavity was conspicuous, even though the specific amino acid residues are not strictly conserved (**Figure 5 - figure supplement 2, Figure 1 – figure supplement 7**). Mutating a pair of adjacent asparagine residues to equivalently sized but negatively charged aspartates (hEMC3^N114D+N117D^) resulted in a dramatic decrease in SQS^378-410^ and a mild decrease in the polytopic terminal insertase client (B1AR) (**Figure 5F**). Meanwhile, mutating a neighboring positively charged residue to an alanine (hEMC3^R180A^), a residue that is conserved in some of the YidC-superfamily insertase proteins (**Figure 5 – figure supplement 4**) (Anghel, 2017), resulted in partial loss of only the tail-anchored insertase client (SQS^378-410^) (**Figure 5 - figure supplement 1**).

Lastly, we surveyed residues closer to the hydrophobic lumenal side of the gated cavity. Lipid density was resolved at positions along the cavity in hEMC and yEMC cryo-EM maps (**Figure 4B**) and the properties of this hydrophobic seal to the lumen are conserved (**Figure 5 - figure supplement 2**). The importance of this hydrophobic seal is suggested by the strong effect of a structurally-mild mutation of a conserved methionine to a leucine (hEMC3^M151L^), which caused significant decrease in both SQS^378-410^ and B1AR abundance (**Figure 5G**).

Mutation of a neighboring aromatic residue (hEMC3^F148L^), contacting both a lipid and the EMC4 C-terminal transmembrane helix, caused a marked decrease in all three client types without altering the levels of EMC subunits (**Figure 5 – figure supplement 1-2, Figure 1 – figure supplement 7**). Together these results indicate that proper EMC insertase function depends on the exact composition of the cavity and not simply on its hydrophobic nature.

### Structural heterogeneity suggests a role for the gate in regulating access to the insertase transmembrane cavity

While the core transmembrane helices of the gated cavity are superimposable in all four of our EMC structures, the adjacent gate helices appear in different relative orientations. The structural variability likely reflects dynamics of the gate (**Figure 4C**). Comparing detergent and nanodisc maps for both species identified two major gate conformations (**Video 2**). One of the conformations, referred to as the closed gate conformation, results in a more occluded membrane cavity. The other conformation, referred to as the open gate conformation, would provide space for client accommodation.

The C-terminal transmembrane helix of EMC4 and ensuing lumenal segment are well resolved in all four structures; however, other regions of EMC4, including the segment connecting the cytoplasmic domain to the transmembrane gate helices, were poorly resolved perhaps owing to mobility. The yEMC detergent map, yEMC nanodisc map, and hEMC detergent map all show the unassigned helices in the closed conformation, preventing client residence in the gated cavity. By contrast, the hEMC nanodisc map reveals an open gate conformation with the unassigned helices shifted away from the transmembrane core to provide space for a client (**Figure 5B**). We note that the conformational heterogeneity and concomitant lower resolution of the gate likely accounts for the challenges in making unambiguous subunit assignments (**Figure 5 – figure supplement 3**), reflected by the three different interpretations reported in recent structures (Pleiner, Tomaleri, and Januszyk, 2020; O’Donnell, Phillips, and Yagita, 2020; Bai, 2020).

Considering the apparent flexibility of the gate, we sought to mutate the EMC4 interfaces resolved in the cytoplasm versus the membrane. As described above, mutating residues that together form a composite binding surface for the cytoplasmic domain of EMC4 (hEMC2^E146A+E149A+Q150A^, hEMC2^E168A + D170A + K173A^, **Figure 3F**), we observed a modest accumulation of the tail-anchored insertase client (SQS^378-410^) (**Figure 3G, Figure 3 – figure supplement 1-2**). Likewise, mutating residues in the center of the gated cavity, close to one of the unassigned helices in the closed gate conformation (hEMC3^V118A + I122A^) (**Video 2, Figure 5 – figure supplement 1**) led to an increase of SQS^378-410^. This SQS^378-410^ accumulation effect stands in contrast to mutating a residue that contacts the lumenal anchor of EMC4 (hEMC3^F148L^), which caused a reduction of SQS^378-410^ levels (**Figure 5 – figure supplement 1**).

**Video 2.**
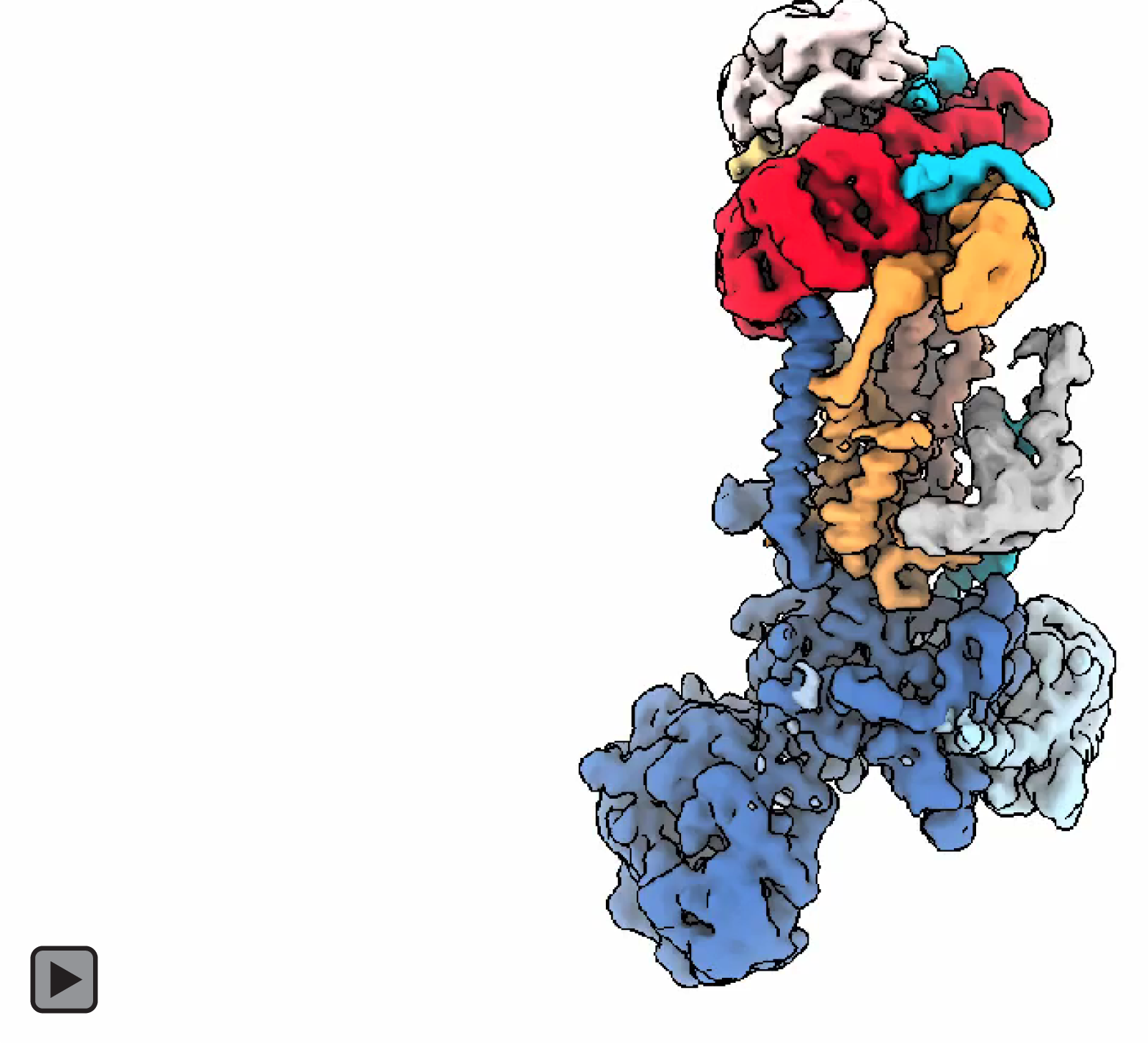
EMC transmembrane cavity gate conformations. Overview of hEMC colored and labeled by subunit. Volume fades away to hEMC nanodisc model. hEMC nanodisc model remains constant as segmented maps of the unassigned gate helices are shown of hEMC detergent, yEMC detergent, and yEMC nanodisc maps. hEMC is colored cyan, yEMC is colored dark cyan, and gate helices are colored in shades of grey and purple as indicated by the label on the left. Two residues (hEMC3 N114 and N117) are shown in stick representation colored magenta and one residue (hEMC3 R180) is colored in gold.

### The lipid-filled cavity is critical for both insertase-dependent and insertase-independent EMC functions

In addition to the gated cavity, the EMC harbors another membrane-accessible cavity. The surface of the lipid-filled cavity includes contributions from EMC1, EMC3, EMC5 and EMC6 (**Figure 6A**). In our structures the EMC2 N-terminus occludes cytoplasmic accessibility to this cavity (**Figure 4D**, **Figure 6A-B**). However, this cavity may be accessible from the membrane or the ER lumen. The respective distance from the cytoplasmic EMC2 N-terminus to the lumenal side of the lipid-filled cavity is approximately 35 Å across, which is close to the average ER membrane thickness (Mitra, 2004).

**Figure 6:**
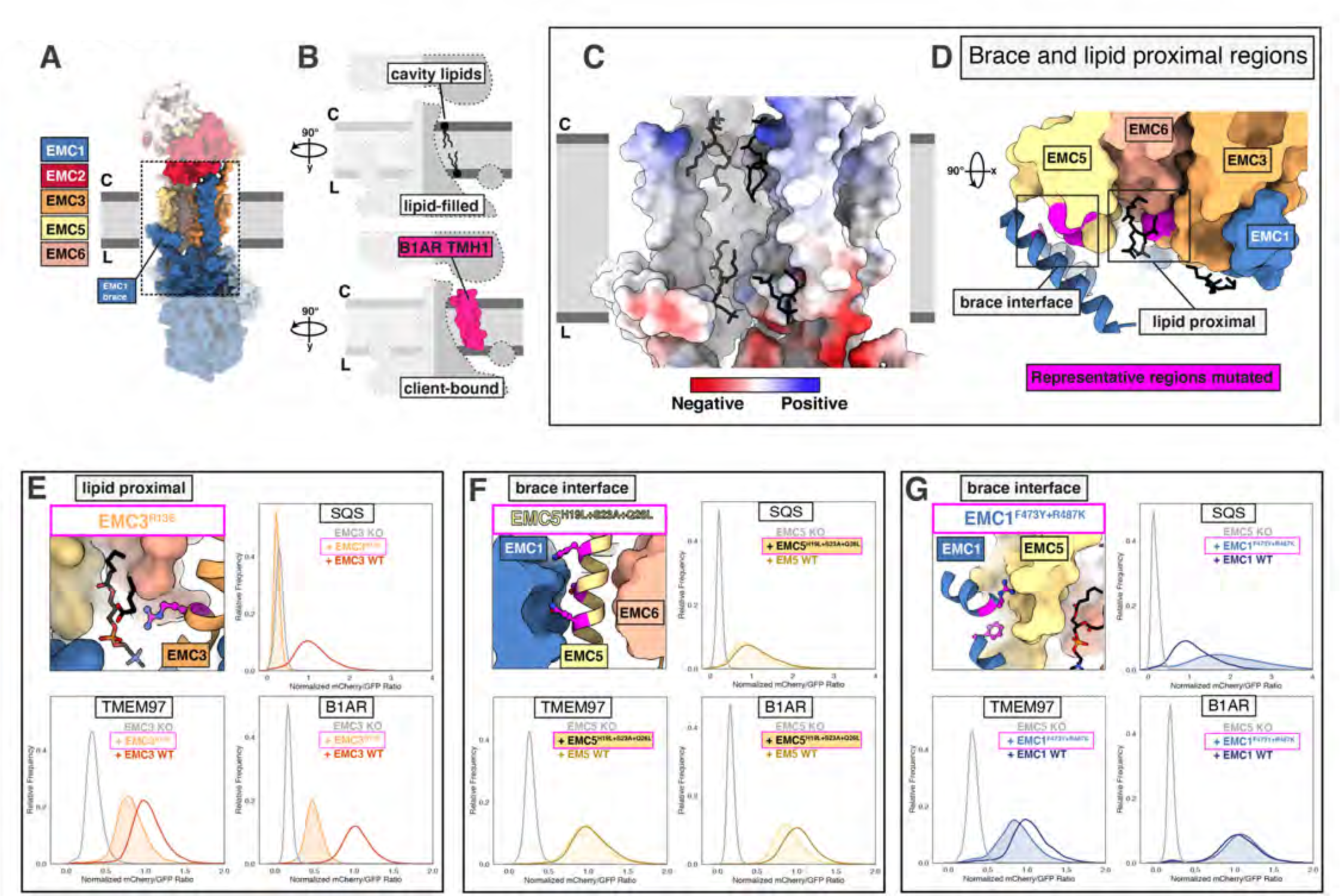
A lipid-filled cavity in the EMC transmembrane domain stabilizes disparate client proteins. A) An EMC1 amphipathic brace helix delineates the boundary of the lipid-filled transmembrane cavity and packs against EMC5. Shown is a surface rendering of the hEMC model in nanodiscs. EMC4, EMC5, EMC6, and EMC1 subunits all contribute to the cavity lining. B) The lipid-filled cavity in the hEMC nanodisc is occupied by several lipid molecules. Cartoon outlines of the gated cavity illustrate that the cavity could in principle allow for occupancy of a client helix (B1AR TMH1), possibly by lipid displacement or movement of the EMC1 brace helix. C) The lipid-filled cavity has a uniform hydrophobic lining. Shown is an electrostatic surface rendering of the hEMC nanodisc structure colored as in Figure 5C. The cytoplasm-membrane interface contains positively charged residues and the lumenal interface contains negatively charged residues. Modeled phospholipid molecules are displayed in black. D) Lipid-proximal and brace interface residues targeted for mutagenesis. Selected regions targeted for mutagenesis are colored in magenta and include brace interface mutations both in EMC1 and EMC5, as well as a lipid-proximal residue in EMC3. E) Fluorescent client reporter stability assay for the hEMC3^R13E^ mutant, which is in close proximity to a modeled POPC molecule. F) As in E) for the hEMC5^H19L+S23A+Q26L^ mutant, which sits at the interface to the EMC1 amphipathic brace helix. G) As in E) for the hEMC1^F473Y+R487K^ mutant, which sits at the interface to the EMC5 transmembrane helices.

The lipid-filled cavity features a uniformly hydrophobic surface (**Figure 6C**) and superimposes across our ensemble of EMC structures. As noted, we resolved several lipids in our cryo-EM maps lining the cavity wall and modeled four POPC (1-palmitoyl-2-oleoyl-sn-glycero-3 phosphatidylcholine) molecules in the hEMC nanodisc map (**Figure 6C**). The residues in close proximity to these lipids are moderately conserved (**Figure 6 – figure supplement 1**). To characterize the functional role of the lipid-filled cavity, we mutated cavity-lining and lipid-proximal residues (**Figure 6D, Figure 6 – figure supplement 2-3**).

Most of these mutations resulted in an increased abundance of the tail-anchored reporter (SQS^378-410^) and mild decrease in production of polytopic client reporters (B1AR, TMEM97). However, one lipid-proximal mutant showed decreased levels of all three client reporter types with varying severity (hEMC3^R13E^) without altering overall EMC levels. An analogous mutation in drosophila EMC3 was recently was reported to cause reduced levels of Rh1 in this mutant background (Xiong, 2020). The amphipathic EMC1 brace helix, which packs against the transmembrane helices of EMC5, is a structural hallmark of the lipid-filled cavity, (**Figure 6D**). Here, mutating interfacial residues from EMC5 (EMC5^H19L+S23A+Q26L^) caused a marked decrease in the N-lumenal polytopic reporter (B1AR) and a mild decrease in the tail-anchored client (SQS^378-410^) with no effect on the TMEM97 reporter (**Figure 6F**).

Unexpectedly, mutating interfacial residues from EMC1 (EMC1^F437Y+R487K^) showed a diametrically opposed phenotype in which B1AR was unaffected, SQS^378-410^ accumulated and TMEM97 levels markedly decreased (**Figure 6G**). Two other interfacial brace mutants which had yet different resulting client flow cytometry profiles (**Figure 6 – figure supplement 3B-C**). The pleiotropic client phenotypes across the panel of interfacial brace mutants suggest that this feature is critical for multiple EMC functions.

### The EMC lumenal domain is crucial for the biogenesis of multi-pass transmembrane proteins

Composed primarily of EMC1, EMC7 and EMC10, the extensive EMC lumenal domain (**Figure 7A**) is important for polytopic client biogenesis and interactions with lumenal chaperones (Luo, 2002; Shurtleff and Ithzak, 2018; Hiramatsu, 2019; Coehlo, 2019). EMC7 and EMC10 are scaffolded on two beta-propellers of EMC1, one distal and the other proximal to the membrane. The lumenal cap differs between hEMC and yEMC, with a four-bladed distal beta-propeller in yeast and eight-bladed distal propellor the human complex (**Figure 7B**). All three lumenal EMC subunits have structural folds known to participate in protein-protein interactions (Reinisch & De Camilli, 2017). Mutations in this lumenal domain have been linked to loss of the EMC complex (Bircham, 2011), a trafficking delay for membrane protein Pma1 (Luo, 2002), and male infertility (Zhou, 2018).

**Figure 7:**
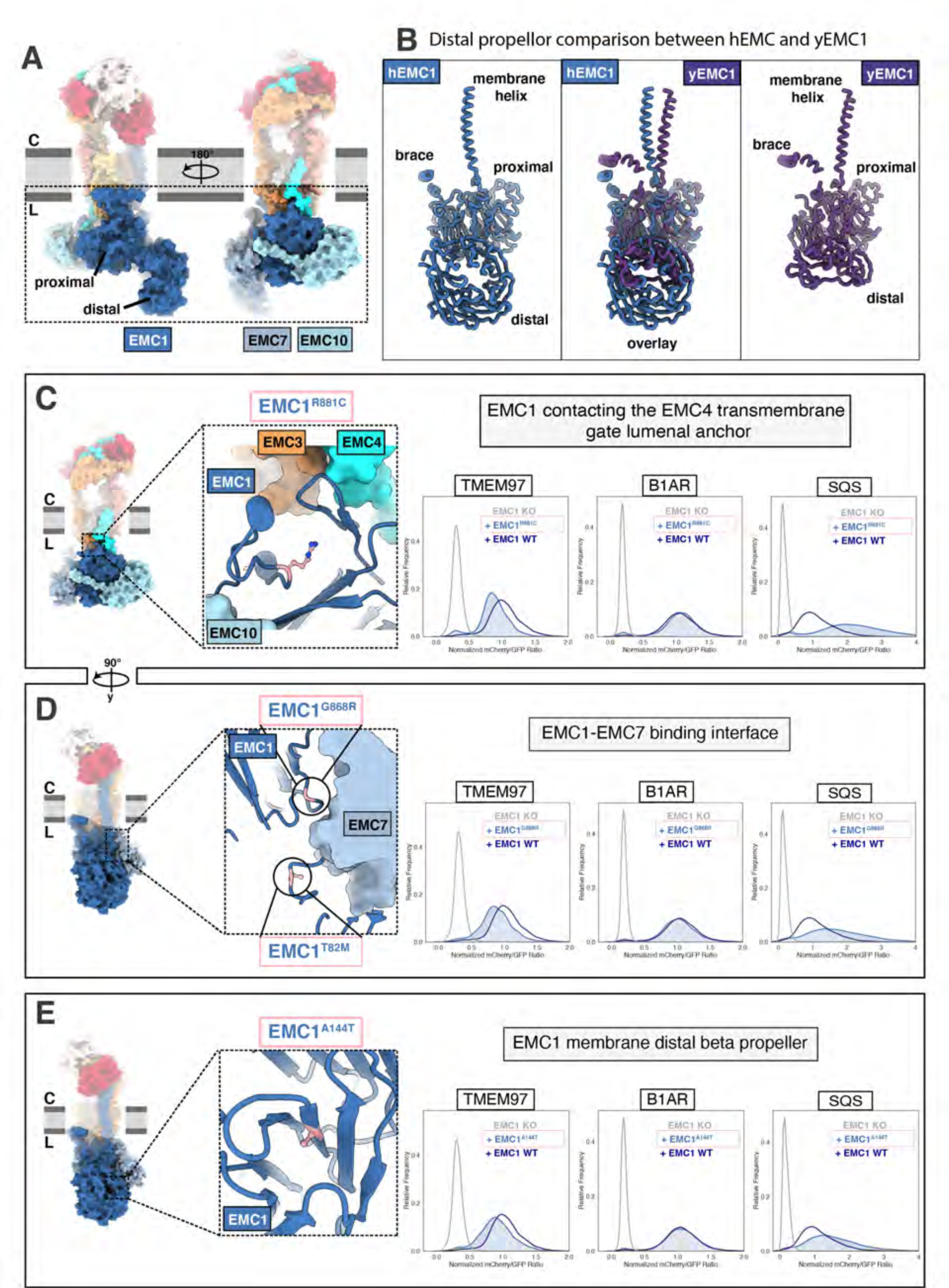
The large EMC lumenal domain is the site for several annotated disease mutations. A) Two views of the hEMC nanodisc structure. Two beta propellers are present in EMC1, one proximal to the membrane and one distal. B) EMC1 is the largest EMC subunit and differs in size between yeast and human. Shown are human EMC1 (nanodisc), an overlay of human and yeast EMC1 (both nanodisc), and yeast EMC1 (nanodisc). C) The hEMC1^R881C^ mutant sits near the EMC4 lumenal gate anchor. Left: Location of the mutation (colored pink). Right: Fluorescent client reporter stability assay for hEMC1^R881C^. D) As in C) for the hEMC^G868R^ mutant. E) As in C) for the hEMC^A144T^ mutant.

Several regions of the lumenal domain form stabilizing interactions with the membrane cavities. The gate helices of the gated cavity are anchored via the embedding of EMC4’s C-terminus within the membrane-proximal EMC1 propeller. The lipid-filled cavity is connected to the ER lumenal domain via the amphipathic EMC1 brace helix, which is tethered to the membrane-proximal EMC1 beta-propeller. The connections between the lumenal domain and the transmembrane cavities could allow for conformational coupling during client handling. Indeed, superimposing the two conformations presented above, the open and closed gate states, revealed not only differences in the transmembrane domain but also a rotation of the lumenal domain relative to the membrane cavities (**Video 3**). The lumenal positioning is consistent for all three of our closed gate conformation reconstructions (hEMC detergent, yEMC nanodisc, yEMC detergent). By contrast the one map with an open gated cavity displayed a lumenal rotation and concomitant shifts in position of the EMC1 brace helix (**Figure 7 – figure supplement 1**). Indeed, our set of interfacial EMC1 brace mutants described above (**Figure 6F-G, Figure 6 – figure supplement 3**), showed differing client phenotypes when mutated from either the EMC1 or the EMC5 side. This suggests a complex conformational interplay between lumenal and transmembrane domains during the engagement of diverse client types.

We investigated several known disease mutations in both conserved and human-specific regions of EMC1 (**Figure 7C-E**, **Figure 7 – figure supplement 2-3)** (Harel, 2016; Abu-Safieh, 2013; Amberger, 2019). One of these disease-associated residues sits near the anchor point for the lumenal EMC4 transmembrane gate helix (hEMC1^R881C^), while the majority are found farther from the membrane (hEMC1^G868R^, hEMC1^A144T^, hEMC1^T82M^) (**Figure 7C-E**). Incorporating each of these disease mutations into our EMC functional assay resulted in lower levels of the N-cytoplasmic polytopic client (TMEM97) and an increase in the level of the tail-anchored client (SQS^378-410^), discussed in more detail below.

Two different EMC1 mutants associated with cerebellar atrophy, visual impairment, and psychomotor retardation (hEMC1^T82M^, hEMC1^G868R^), map to the hinge region between the EMC1 beta propellers where EMC7 binds (**Figure 7D**). Both of these mutants, at this protein-protein interface, result in depletion of the N-cytoplasmic polytopic client (TMEM97). EMC7 and EMC10 form beta-sandwich domains on either side of the membrane-proximal beta-propeller of EMC1, and contact each other across the EMC1 surface. Consistent with our structures, coupling of these subunits is supported by the prior finding that in the absence of EMC7, EMC10 is also lost from the complex while the other EMC components appear unaffected (Shurtleff and Ithzak, 2018). EMC7 and EMC10 have been proposed to be auxiliary components with weaker phenotypes compared to core EMC subunits (Jonikas, 2009; Shurtleff and Ithzak, 2018; Dickinson, 2016). Upon deleting EMC7, multi-pass transmembrane clients are retained in the ER but tail-anchored clients decrease in abundance (Shurtleff and Ithzak, 2018).

Several features of our data suggest dynamic association of EMC7. Density for the EMC7 beta-sandwich at the hinge between the two EMC1 beta propellers was relatively weak in the consensus hEMC nanodisc map (**Figure 2 – figure supplement 4**). Additional rounds of 3D classification revealed two distinct classes, one with clear density for EMC7 and one with weak density in this region. Mass spectrometric analysis of purified hEMC, however, revealed that the abundance of EMC7 was similar to that of the other EMC components (**Figure 1 – figure supplement 2; Source Data 1 - Supplementary Table 3**). Both reconstructions, with and without density for the EMC7 lumenal domain, displayed well-resolved density for EMC10. Together, we conclude that EMC7 is associated with EMC1 in two different conformational states of EMC7 with potentially distinct functions.

The OMIM database (Amberger, 2019) lists a mutation of unknown significance linked to retinitis pigmentosa (hEMC1^A144T^) residing in the EMC1 distal propeller (**Figure 7E**).

Additionally, we also generated mutations in two surface exposed patches of the membrane-distal EMC1 beta-propeller projecting into the lumen (hEMC1^R69D^, hEMC1^G71S^, hEMC1^H93D + E138D + N282K^, **Figure 7 – figure supplement 2-3**). Overall, these mutations displayed the same client effect: a decrease in the N-cytoplasmic polytopic client reporter (TMEM97), no change in the N-lumenal polytopic client reporter (B1AR), and accumulation of the tail-anchored client reporter (SQS^378-410^) (**Figure 7 – figure supplement 2-3**). Upon identifying antibodies against yEMC we observed that the top two antibodies bind to a similar extended loop in the distal yEMC1 beta-propeller, perhaps suggesting that this site is accessible for co-factor binding in the ER. Intriguingly, this region of the lumenal domain corresponds to the region where hEMC1 has an expanded distal beta-propeller. Taken together, the data provide evidence that the lumenal domain is functionally coupled to the broader EMC role in transmembrane client stabilization. Moreover, these data support that the EMC is acting as a holdase chaperone to shield polytopic clients from degradation while they are folding to their functional form.

**Video 3.**
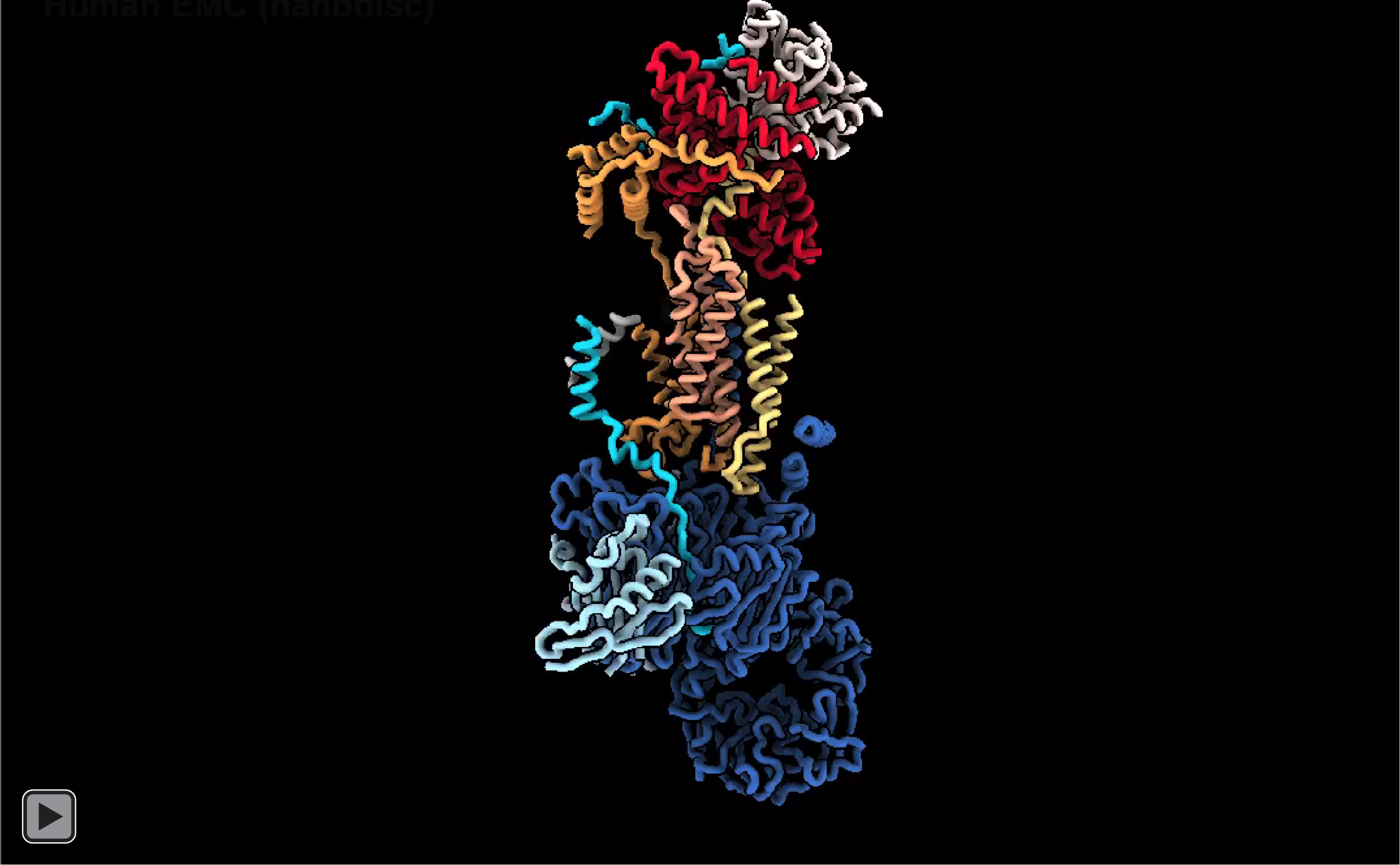
hEMC lumenal domain differences between nanodisc and detergent models1. Overview of hEMC nanodisc model colored and labeled by subunit. Structural landmarks are labeled. hEMC detergent model (colored grey) fades in and both models rotate. As the models rotate several structural features are highlighted.

## Discussion

Our collection of yeast and human EMC structures revealed the intricate and dynamic architecture of this multifunctional transmembrane molecular machine. The structures served as the starting point for our systematic dissection of EMC’s multifaceted functions by exploring the impact of structure-based mutations on the ability of the EMC to support the biogenesis of representative members of three classes of membrane proteins: SQS, a tail-anchored protein, which exploits EMC’s C-terminal insertase activity; B1AR, which relies on EMC’s N-terminal insertase activity; and TMEM97, a polytopic membrane protein, which depends on the EMC for its biogenesis but does not rely on either of EMC’s terminal insertase activities. Our data revealed that a conserved dual membrane cavity architecture supports the biogenesis of this diverse panel of transmembrane clients.

Overall, our studies present a nuanced picture of EMC’s multifunctionality, revealing structural regions that differentially impact production of the three distinct client types. Unexpectedly, we also find that alterations to either the cytoplasmic or lumenal domain of EMC lead to enhanced abundance of the TA substrate. Moreover, our work provides a foundational framework for understanding how discrete yet allosterically coupled regions of the complex enable the multiple functions of the EMC to support membrane protein biogenesis. Taken together these studies suggest a model in which the EMC differentially regulates the biogenesis of distinct membrane proteins, thereby contributing to cellular coordination of membrane protein abundance in accordance with physiological needs.

We propose a model of the EMC functioning both as a terminal insertase as well as a holdase chaperone that is potentially modulated by post-translational modifications, lipid interactions, and protein-protein interactions [**FIGURE 8**].

**Figure 8:**
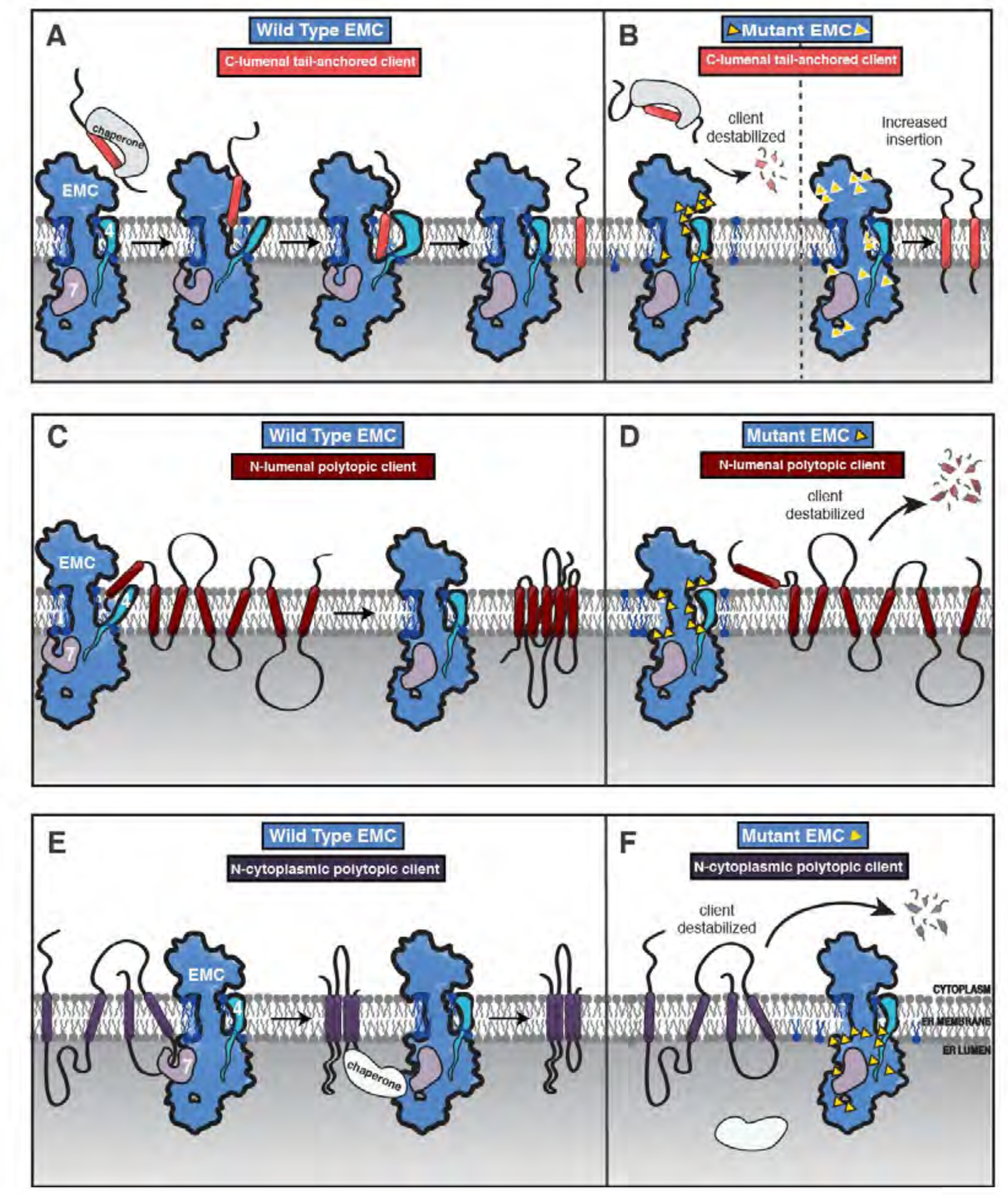
Model of coordinated EMC functions. A) Model of EMC insertase function for a C-lumenal tail anchored client. Cytosolic factors bring post-translationally localized clients to the ER. Then the client engages the EMC cytoplasmic domain. The polar roof modulates entry into the gated cavity. A hydrophobic slide facilitates the client helix fully entering the cavity. A lateral movement of the gate releases the client helix into the membrane and the EMC gate closes. B) Our mutagenesis data provide the following insights into EMC regions of functional importance for each of the three client types we tested. Mutants are depicted by yellow triangles. Tail anchored client (coral) abundance was depleted upon mutagenesis of the cytoplasmic domain entrance to the gated cavity, polar and charged residues at the cytoplasm-membrane boundary, residues along the length of the gated cavity, in the hydrophobic seal to the lumen, and lipid interacting residues in both cavities (left). We also observed a subset of mutants that resulted in higher levels of the C-lumenal tail anchored client (right) that are positioned in the cytoplasmic domain cap, throughout the ER lumenal domain, and one mutation at the center of the gated cavity. C) The EMC facilitates biogenesis of N-lumenal polytopic client protein B1AR (dark red). D) Regions important for B1AR stability primarily map to the transmembrane region of the EMC structure, with depletion observed for lipid interacting residues on both sides of the cavity, the polar entrance roof of the gated cavity, and the EMC1 brace helix. E) The EMC facilitates biogenesis of N-cytoplasmic polytopic client protein TMEM97 (dark purple). F) Regions important for TMEM97 stability were primarily located in the lumenal domain spanning both propellers, in EMC1. In addition to these lumenal regions there was a depletion of TMEM97 at the lipid-interacting positions at the lumenal interface of both membrane cavities of the EMC.

### Terminal insertase clients require an embedded insertase module within the EMC

EMC3’s fold at the interface between the cytoplasm and membrane forms the core of the gated cavity and is reminiscent of proteins from the YidC family of insertases (Borowska, 2015; Dalbey, 2015; Anghel, 2017). Indeed, mutations in either the cytoplasmic or transmembrane domains of EMC3 establish that these features are critical for terminal helix insertase activity. In light of our observation of multiple gate conformations, we speculate that these conformations modulate insertion and release into the ER membrane.

Notably, mutating the surface of the cytoplasmic cap, which extends beyond the EMC3 cytoplasmic helices towards EMC8/9, resulted in an unexpected increase in C-tail anchor client (SQS^378-410^) abundance. Of the three clients analyzed, SQS was the only one to show enhanced levels. It is unclear if this enhancement is SQS-specific or representative more broadly of all post-translationally targeted EMC tail-anchored clients. Future studies will be required to address if this is due to regulated insertion of SQS by the EMC, parallel pathways for inserting SQS into the membrane (i.e. mediated by TRC40/GET), and/or slower cytoplasmic clearance of chaperone-bound SQS.

### Both EMC cavities have resolved lipids and are critical for client biogenesis

Both the N-terminal (B1AR) and C-terminal insertase (SQS) clients depend on the EMC gated cavity. Indeed, both the SQS tail-anchored helix and the first transmembrane helix of B1AR are moderately hydrophobic, with polar residues near the cytoplasmic end of the transmembrane helix, and both showed a strong dependence on the gated cavity.

Nevertheless, our panel of mutants revealed some notable differences in the handling of these two client types. B1AR showed more dependence than SQS on the lipid-filled cavity in contrast to mutants elsewhere in the complex. One possible reason could be due to differences in the mechanism of initial engagement: SQS is targeted to the ER by cytoplasmic chaperones, while B1AR is targeted by SRP. Another key difference is that B1AR is polytopic and needs to overcome the additional challenge of tertiary transmembrane packing to reach its folded state. Future work will address the interplay between B1AR synthesis and its co-translational engagement with the translocon versus the EMC.

### The EMC lumenal domain orchestrates holdase chaperone function important for polytopic clients

Unlike the two terminal insertase clients we investigated, TMEM97 biogenesis was negatively impacted by mutation of the lumenal EMC1. The depletion of TMEM97 observed in these mutant backgrounds is consistent with the lumenal domain contributing to a holdase chaperone function, passively shielding its client while it is being synthesized and/or folded (Zhang, 2017). Interestingly, the diametrically opposed phenotype of mutants in the EMC lumenal domain on SQS raises the possibility that occupancy by one type of client can support an EMC conformation that is unfavorable for receiving the other. Alternative conformations could establish competition between client types for EMC occupancy. One explanation for this observation is that there is a conformational change between the insertase-active versus the holdase-active states.

Interestingly, we identified at least two EMC conformations in our collection of structures, and EMC may adopt different conformations in various client and cofactor-engaged states.

In yeast, the polytopic clients co-purifying with the EMC are also glycosylated. One possible model is that the putative carbohydrate binding domains in EMC7 or EMC10 directly contribute to engagement with client proteins. We speculate post-translational modifications on clients and the EMC could modulate function including client binding, chaperone binding, or regulating signaling in response to cellular cues.

### Potential role of the EMC as a master regulator of membrane protein biogenesis as the basis for its pleiotropic phenotypes

Why does the cell use a multifunctional EMC molecular machinery rather than specialized machinery for each of the functions encompassed by the EMC? Considering that the cell already has general machinery (Sec61 translocon) and tail-anchor insertase machinery (GET/TRC complex), we speculate that the EMC coordinates biogenesis of diverse membrane proteins. Several observations suggest broader roles of the EMC as an integrator of information sensing the protein and lipid environment and coordinating its multiple activities, including the regulating the biogenesis of membrane proteins. For example, the initial identification of the EMC included numerous genetic interactions with both protein and lipid synthesis factors in yeast (Jonikas, 2009) and these disparate interdependencies have been subsequently observed in numerous species including human EMC (Lahiri, 2014; Tang, 2017; Guna, 2018; Volkmar, 2019; Volkmar, 2020).

Also, several client proteins are enzymes or cofactors involved in multiple stages of lipid synthesis or trafficking, and this may provide a unifying explanation for the range of genetic interactions and co-essentiality observations reported to date (Guna, 2018; Shurtleff and Ithzak, 2018; Volkmar, 2018; Tian, 2019; Wainberg, 2019; Corradi, 2019; Volkmar, 2020). In this regard, one structural feature of particular interest is the EMC1 amphipathic brace, which resides adjacent to the lipid-filled cavity. This conserved feature sits within the interfacial membrane boundary, raising the possibility that it can modulate the lipid or protein composition of this cavity. Notably, several other membrane proteins involved in ER homeostasis, including Opi1 and Ire1, also contain amphipathic helices that have been proposed to sense the properties of the lipid bilayer (Volmer, 2013; Jacquemyn, 2017; Halbleib, 2017; Hofbauer, 2018; Cho, 2019). Future work will explore how the EMC overall, and the EMC1 brace helix in particular, govern client release into the membrane, interface with the local structure of the lipid bilayer, and play roles in specific client-lipid interactions.

In addition to the three client classes we investigate here, it is clear that EMC has a broader range of clients including multi-protein assemblies (Richard, 2013; Talbot, 2019), lipid modulating proteins (Volkmar, 2018), lipid binding proteins (Salas-Estrada, 2018; Sejdju, 2020), and those with helices that do not span the bilayer (Lin, 2019; Ngo, 2019). The compartmentalization and interdependence that we observe for effects of mutations on client handling provide a foundation for understanding this multifunctionality. We propose that the complexity of the EMC machine, combining insertase and holdase chaperone functions within one molecular machine, has arisen to mitigate the error prone biogenesis of a diverse range of membrane spanning proteins in the dynamic environment of the ER.

## Materials and Methods

Reagents used for experiments described and reagents made as part of this study are listed in Supplementary Table 2.

### Cell Line Maintenance

K562 dCas9 KRAB cells were grown in RPMI 1640 (GIBCO) with 25 mM HEPES, 2 mM l-glutamine, 2 g/L NaHCO3 and supplemented with 10% (v/v) fetal bovine serum (FBS), 100 units/mL penicillin, 100 µg/mL streptomycin, 2 mM l-glutamine. HEK293T cells were grown in Dulbecco’s modified eagle medium (DMEM, GIBCO) with 25 mM d-glucose, 3.7 g/L NaHCO3, 4 mM l-glutamine and supplemented with 10% (v/v) FBS, 100 units/mL penicillin, 100 µg/mL streptomycin. All cell lines were grown at 37°C. All cell lines were periodically tested for Mycoplasma contamination using the MycoAlert Plus Mycoplasma detection kit (Lonza).

### DNA transfections and virus production

Lentivirus was generated by transfecting HEK39T cells with standard 4^th^ generation packaging vectors using TransIT-LT1 Transfection Reagent (Mirus Bio). Media was changed 10 hours post-transfection. Viral supernatant was harvested 60 hours after transfection, filtered through 0.45 µm PVDF filters and frozen prior to transduction.

### Knockout EMC Cell Lines

A single and dual knockout guide system was developed in the pX458 backbone (Addgene plasmid # 48138) with guides targeting EMC1, EMC2, EMC3, or EMC5 (see reagents table). Targeting guides were selected using the Broad’s guide selection tool (https://portals.broadinstitute.org/gpp/public/analysis-tools/sgrna-design). For the single EMC5 knockout system, an EMC5 targeting guide was cloned into pX458 by digesting with BbsI and ligating to annealed oligos for the EMC5 sgRNA. For the dual knockout system, a four-step cloning process generated the final knockout plasmid: 1) Each of the two guides targeting the same locus were individually cloned into pX458. 2) Then pX458_sgRNA1 was digested with XbaI 3) SgRNA2 cassette from pX458_sgRNA2 was PCR amplified with oligos containing overhangs spanning the XbaI cloning site and purified. 4) Finally, the final dual guide vector was generated by Gibson cloning (NEBuilder).

To generate the EMC knockout cell lines, K562 dCas9 KRAB cells were nucleofected with the respective EMC knockout plasmids using Lonza SF Cell Line 96-well Nucleofector^TM^ Kit (V4SC-2096). Two days post nucleofection, GFP-positive cells were single cell sorted into 96-well plates using BD FACS AriaII. After colonies from single cells grew out, genomic DNA was isolated using QuickExtract (Lucigen), the sgRNA-targeted sites were PCR amplified and then NGS-sequenced via Genewiz’s EZ-Amplicon service. Sequencing data was analyzed and aligned to the respective reference alleles in the human genome. Clones whose alleles harbored only indel mutations for EMC1, EMC2, EMC3, and EMC5 (full knockouts) respectively were further validated on the protein level.

### Dual fluorescent EMC client reporter cell lines

Dual client reporters for TMEM97, ADRB1 (protein name: B1AR), and FDFT1 (protein name: SQS) were introduced lentivirally into each of the EMC1, EMC2, EMC3, and EMC5 knockout cell lines. TMEM97 and ADRB1 full length sequences were used with a C-terminal tag -mCherry-P2A-GFP. The sequence for FDFT1 transmembrane domain (SQS^378-410^) was tagged N-terminally with GFP-P2A-mCherry- and an opsin tag on the C-terminus as used in a prior study (Guna, 2018). Three days post-transduction, GFP/mCherry positive cells were sorted on BDAriaII. Sequences for these constructs are available in Supplementary Table 2.

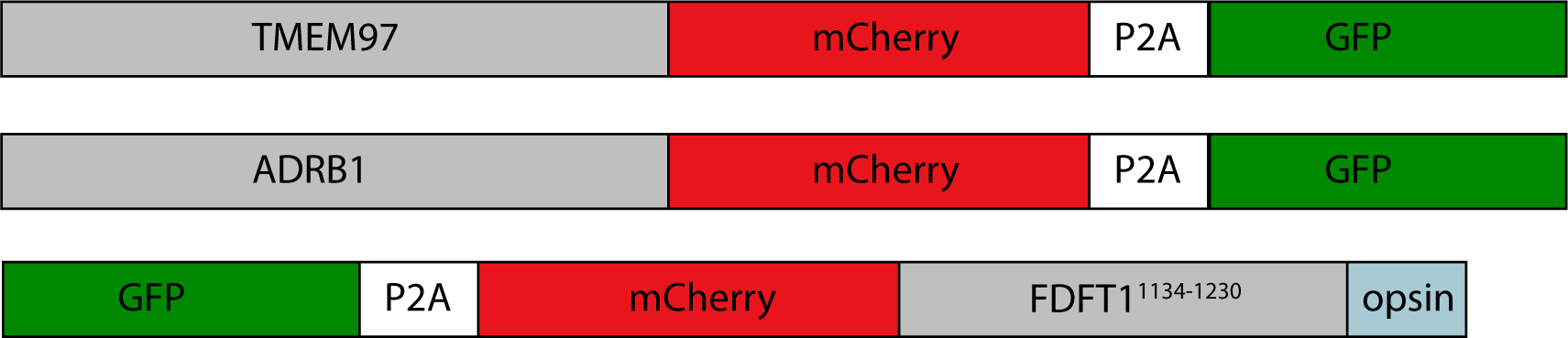

### Mutant EMC cell lines

The EMC mutant genes were synthesized and cloned by Twist into pKDP119-SFFV-[insert site]-IRES-Puro-P2A-BFP. For EMC subunit mutation details refer to the reagents table. Mutant EMC cell lines were generated by lentiviral introduction of the respective EMC mutant subunit into the respective knockout cell lines (EMC1, EMC2, EMC3 or EMC5) containing the dual fluorescent reporters for each EMC client (pKDP110_ADRB1_mCherry_P2A_GFP, pKDP111_TMEM97_mCherry_P2A_GFP, or GFP_P2A_mCherry_FDFT1_TMD_opsintag). The expression of each fluorescent reporter was read out 6 days after puromycin selection in each of the EMC mutant cell lines.

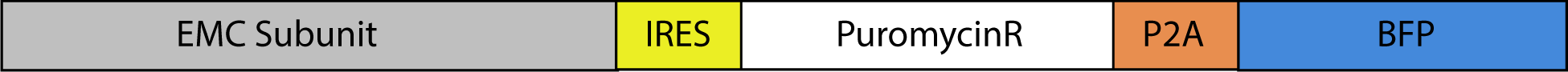

### Flow analysis

For each EMC mutant cell line, 20 000 live cells were recorded on Attune NxT flow cytometer. FlowCal flow analysis package was used for analysis in Python. First, live cells were gated based on FSC/SSC. Then GFP (BL1-A) and mChery (YL2-A) were plotted for each mutant and control cell line. mCherry:GFP intensity ratios were calculated for individual cells in each cell line. Fluorescence ratios for each substrate in an EMC mutant cell line were normalized to the mCherry:GFP ratio of the same substrate in the EMC wild type rescue cell line. Distributions of fluorescence ratios were plotted as histograms in Python using seaborn.

### Western Blotting

Cell pellets were lysed using lysis buffer (20 mM Tris pH 7.5, 150 mM NaCl, 5 MgCl2, 1% Triton x-100, 1 mM DTT, 24 U/ml Turbo DNase (Ambion). Clarified lysate was quantified and samples were boiled with 4x LDS sample (Thermo Fisher, NP0007) buffer for 5 mins at 95C. Samples were separated on 4-12% or 12% Bolt Bis-Tris Plus Gels (Invitrogen, NP0322PK2). Proteins were transferred onto nitrocellulose membranes using Bio-Rad Trans-Blot Turbo transfer system. Membranes were blocked in Odyssey Blocking Buffer (LI-COR, 927-50000) for an hour at room temperature. Blocked membranes were incubated with primary antibody diluted in TBST and incubated overnight at 4C on a shaker. Primary antibodies were detected by incubating membranes with 1:10000 dilution of IRDye-conjugated (LI-COR) secondary anti-mouse and anti-rabbit antibodies for 1 hour at room temperature. Blots were visualized using LI-COR imaging system. The primary antibodies used in this study are in the reagents table.

### Yeast strains

Strain BY4741 and BY4742 were used as the wild-type parental strains for the creation of the yEMC overexpression strain. Yeast homologous recombination (Rothstein, 1991) was used to generate yeast strains. For the overexpression strain, the endogenous promotor for each yEMC subunit (yEMC1, yEMC2, yEMC3, yEMC4, yEMC5, yEMC6, yEMC7, yEMC10) were replaced with a TEF2 promoter. In addition, EMC5 was tagged at the C-terminus with linker-TEV-linker-3xFlag. Auxotrophic markers and drug selection markers in both BY4741 and BY4742 were employed to add this promoter modification to all of these eight subunits and the two strains were crossed to create the resulting BY4743 strain used for immunoprecipitation. Endogenous EMC yeast strain was made using W303a wild type parental background (leu2-3,-112; his3-11,-15; trp1-1; ura3-1; ade2-1; can1-100; MATa). Homologous recombination was used to integrate a linker-TEV-linker-3xFlag at the C-terminus of EMC coding sequence. Genomic PCR was conducted to verify integration.

### Design and purification of fragments antigen binding (Fab) DH4 and DE4

Fabs were identified as described in these studies (Kim, 2011; Wu, 2012). Overexpressed yEMC solubilized in DDM as described above was biotinylated and streptavidin magnetic beads were used to capture yEMC, which was then subjected to a Fab phage library. Unbound Fabs were washed away and then binding Fabs were eluted and analyzed by ELISA. Two Fabs were identified binding EMC, Fab DH4 and DE4.

### Purification of DH4 and DE4 Fabs

Plasmid with either Fab DH4 or DE4 were transformed into BL21 Gold Star cells and plated onto agarose plates with 2x YT + 2% glucose + Ampicillin. Cultures were inoculated from resulting colonies for overnight growth at 30 °C into 2xYT + 2% glucose + Amp. In the morning dilute overnight culture to OD600 of 0.05 in 1L, in a 2.8 L flask of 2xYT + 0.1% glucose + Amp. Grow the culture at 180 rpm at 37 °C shaker until OD600 of 0.6, then, switch to shaking at 19 °C for 1 hour. Next, induce with 0.4 mM IPTG. Shake at 180 rpm at 19 °C for 18-20 hours. Spin 1L cultures down at 3500 rpm in large Beckman Centrifuge at 4 °C for 20 minutes in (8.1 rotor). Discard media and gently resuspend cell pellet in ice-cold 20 ml in Buffer 1 (0.2 M Tris pH 8.0, 0.5 mM EDTA, 0.5 M Sucrose) on ice. Transfer the resuspended cells from step 2 into 2 smaller JLA 25.5 centrifuge tubes. Add 20 ml of ice cold ddH2O with 2x protease inhibitor cocktail (Roche Complete Ultra, Millipore Sigma 5056489001) from step 3 to the resuspended pellets.

Incubate at on ice for one hour occasionally swirling samples gently. Spin periplasmic fractions at 13,000 x g for 15 min, 4 °C, rotor 25.50. Wash 500uL Ni resin (Qiagen, Ni-NTA, 30210) per periplasmic fraction four times in Buffer 2 (50 mM Tris pH 8.0, 250 mM NaCl). Add MgCl2 and imidazole to a final concentration of 10 mM to each periplasmic fraction. Add beads to periplasmic fractions and nutate at 4 °C for 2 hours. Spin down beads at 2000 x g, 10 minutes, 4 °C. Transfer beads either to a 50 mL gravity column. Wash the beads with 20 column volumes of Buffer 3 (50 mM Tris pH 8.0, 500 mM NaCl, 20 mM Imidazole). Elute protein with 3 column volumes of Buffer 4 (50 mM Tris pH 8.0, 500 mM NaCl, 300 mM Imidazole). Analyze eluate by SDS-PAGE 4-12% Invitrogen (Invitrogen, NP0321PK2). Fabs as two bands run around 30 kDa in reducing conditions, or 50 kDa in non-reducing conditions. Dialyze eluate O/N in Dialysis cassette 10 kD molecular weight cutoff at 4 °C against 150 mM KOAc, 20 mM HEPES pH 6.8.

### Purification of overexpressed yeast EMC5-3xflag

The OE-Emc5-3xflag yeast strain were grown in YEPD media in a 40 L fermenter, harvested and flash frozen in liquid nitrogen. Cell pellets were thawed and diluted in lysis buffer (50 mM HEPES pH 6.8, 150 mM KOAc, 2 mM MgOAc, 1 mM CaCl2, 0.2M Sorbital, 2x Protease Inhibitor). Bead beating (10 times à 1 minute on, 2 minutes off) was used to lyse cells. For 25g of cells, 0.1 mm cold beads were added and lysis buffer up to the top of the 50 mL canister. After lysis, beads were filtered and solution centrifuged at 10,000 xg for 10 minutes. Supernatants were ultracentrifuged at 42,000 RPM (Ti 45 rotor) for 2 hours. Supernatant was discarded. Membrane pellet was combined with the lipid layer, and resuspended in lysis buffer and then a precooled dounce homogenizer was used to dounce 20 times. Membranes were aliquoted and flash frozen in liquid nitrogen. On ice, 150 mL of solubilization buffer (50 mM HEPES pH 6.8, 150 mM KOAc, 2 mM MgOAc, 1 mM CaCl2, 15% glycerol, 1% b-DDM, 2x Protease Inhibitor) was added incrementally to 7.5g of thawing membranes, nutated at 4°C for 1 hour in JA 25.5 rotor tubes, and centrifuged at 20,000 rpm for 45 minutes.

Meanwhile 2.5 mL of *α*FLAG agarose beads (Millipore A2220) were rinsed in 50mL of low salt buffer (50 mM HEPES pH 6.8, 150 mM KOAc). Supernatant was added to *α*FLAG beads and nutated at 4°C for 2 hours. Resulting solution was applied over a glass column. After flowing through unbound solution, *α*FLAG beads were washed with 100 mL low salt buffer, 100 mL high salt buffer (50 mM HEPES pH 6.8, 300 mM KOAc, 0.05% b-DDM), and 100 mL low salt buffer. *α*FLAG beads were resuspended in 10 mL of low salt buffer and 300 uL of TEV (1.15mg/mL) was added and nutated overnight at 4°C. Removed supernatant from beads by low speed spin and applied over 500 uL of NiNTA beads equilibrated with low salt buffer to remove excess TEV. Flow through glass column and collect supernatant. Using a 100 kD concentrator (Millipore, UFC910008) solution was concentrated to 2mg/mL. Concentrated EMC protein was applied to the Akta Explorer Superose 6 Increase column (Cytiva, 29091596) for size exclusion chromatography in the size exclusion buffer (20 mM HEPES pH 6.8, 150 mM KOAc, 0.05% b-DDM). Fractions were evaluated by SDS-PAGE Coomassie stain and negative stain electron microscopy then EMC peak fractions were pooled and incubated with 2x molar excess of Fab, either Fab DH4 or Fab DE4, for 30 minutes on ice. Solution was applied to Akta Explorer Superose 6 Increase for size exclusion of Fab bound EMC. Resulting EMC-Fab fractions were evaluated by SDS-PAGE Coomassie stain and EMC-Fab peak fractions were pooled.

### Purification and nanodisc reconstitution of endogenous yeast EMC5-3xflag

Yeast was grown in rich media (YPAD) in a 65L fermenter until OD 2.6. Cell pellets were harvested and flash frozen in liquid nitrogen. Pellets were ground using three cycles in a French press. As above, the resulting solution was ultracentrifuged to separate membranes, dounced to homogenize, and flash frozen in liquid nitrogen. Thawed membranes were solubilized in 1% b-DDM (Anatrace, D310) nutating at 4°C for 1 hour then centrifuged to separate solubilized membranes from the pellet. Supernatant was applied to equilibrated *α*FLAG beads, nutated at 4°C for 1 hour, and applied over a disposable plastic column at 4°C. *α*FLAG beads were washed with low salt buffer and high salt buffer. Then washed with low salt buffer with b-DDM+CHS (Anatrace, CH210) (10:1) in place of b-DDM. *α*FLAG beads were then transferred to a 15 mL Eppendorf tube for TEC cleavage and nanodisc reconstitution.

Bio-Beads SM-2 (Bio-Rad) were prepared ∼400 uL biobeads, rinsing with EtOH, and then water four times. Yeast Extract Total (Avanti Polar Lipids, 190000C-100mg) was prepared by transferring chloroform resuspended solution to a glass vial, drying the lipids into a film with nitrogen gas, drying in a vacuum desiccator overnight, and then solubilizing the lipids first in water and then in size exclusion buffer with DDM+CHS by bath sonication, aliquots stored at -20 °C until use. 200 uL of TEV protease (5 mg/mL) and 150 uL of 1mg/mL Yeast Total Extract solubilized in b-DDM+CHS, at room temperature for 30 minutes. Then added MSP1D1, purified as described previously (Ritchie, 2009), to a ratio of 200:10:1 (Yeast total extract:MSP1D1:EMC), at 4°C for 10 minutes. Then activated Bio-Beads SM-2 (Bio-Rad), ∼300 uL, were added and nutated overnight. On-bead reconstitution employed adapted from (Laverty, 2019). In the morning ∼100 uL more Bio-Beads SM-2 (Bio-Rad) were added and 2x molar excess of FabDH4, nutated for another hour. Beads and solution applied to an EconoPac column (Bio-Rad). Flow through was collected and solution was applied to a 100 kD (Amicon) concentrator. Resulting concentrated EMC was applied to the Akta Explorer Superose 6 Increase column for size exclusion chromatography. Peak fractions were pooled for SDS-PAGE Coomassie stain, negative stain, and cryo-EM evaluation.

### Cryo-EM Sample Preparation and Data Collection for yEMC

#### Overexpressed EMC + Fab DE4 in b-DDM

Following size exclusion sample was prepared for cryo electron microscopy. 3 µL of sample (0.1mg/mL EMC + Fab DE4 in 20mM HEPES pH 6.8, 150 mM KOAc, 0.05% b-DDM) was applied to the grid, incubated for 10 seconds, then blotted with no offset for 6.5 seconds and plunge frozen in liquid ethane using a Vitrobot Mark III at 5° C, Whatman #1 filter paper, and 100% humidity. Protein was frozen on glow discharged Ultrathin Carbon Film on a Lacey Carbon Support Film (Ted Pella, 01824) and stored under liquid nitrogen until imaging. This dataset was collected on the 300 kV Technai Polara at UCSF with a 30 µm C2 aperture, 100 µm Objective aperture, and K2 Summit detector operated in super-resolution mode. 1536 micrographs were collected using SerialEM (Mastronarde, 2005) at a magnification of 31,000X (0.6078 Å/ super resolution pixel) as dose-fractionated stacks of 40 frames x 0.2 second exposures (1.42 e^-^/Å^2^) for a total dose of ∼56.85 e^−^/Å^2^ (see Table S1).

#### Overexpressed EMC + Fab DH4 in b-DDM

Following size exclusion sample was prepared for cryo electron microscopy. 3 µL of sample (0.1mg/mL EMC + Fab DH4 in 20mM HEPES pH 6.8, 150 mM KOAc, 0.05% b-DDM) was applied to the grid, incubated for 10 seconds, then blotted with no offset for 7 seconds and plunge frozen in liquid ethane using a Vitrobot Mark III at 4° C, Whatman #1 filter paper, and 100% humidity. Protein was frozen on glow discharged Ultrathin Carbon Film on a Lacey Carbon Support Film (Ted Pella 01824). This dataset was collected at the HHMI Janelia Research Campus on Titan Krios 2, a 300 kV microscope equipped with a 50 µm C2 aperture, 70 µm objective aperture, and K2 Summit detector operated in super-resolution mode. 3357 micrographs were collected using automated SerialEM (Mastronarde, 2005) collection with defocus range set between -1 and -3 µm at a magnification of 22,500X (0.655 Å/ super resolution pixel) as dose-fractionated stacks of 50 frames x 0.2 second exposures (1.165 e^-^/Å^2^) for a total dose of ∼58.3 e^−^/Å^2^ (see Table S1).

#### Endogenous EMC + Fab DH4 in MSP1D1-Yeast Total Extract nanodisc

Following size exclusion sample was prepared for cryo electron microscopy. 4 µL of sample (∼0.8mg/mL EMC + Fab DH4 in nanodisc in 20mM HEPES pH 6.8, 150 mM KOAc, 0.05% b-DDM) was applied to the grid from the left side, then blotted with no offset for 2.5 seconds, then another 4 uL of sample was applied to the right side of the grid (without glow discharge) and blotted for 3.5 seconds, and plunge frozen in liquid ethane using a Vitrobot Mark IV at 4° C, Whatman #1 filter paper, and 100% humidity. Protein was frozen on R 1.2/1.3 grids with 300 Au mesh (Quantifoil, Germany). This dataset was collected at UCSF on the Titan Krios 2, a 300 kV microscope equipped with a 70 µm C2 aperture, 100 µm objective aperture, and K3 detector operated in CDS mode. 5949 micrographs were collected using automated SerialEM (Mastronarde, 2005) collection with defocus range set between -0.8 and -2 µm at a magnification of 105X (0.4265 Å/ super resolution pixel) as dose-fractionated stacks of 100 frames x 0.06 second exposures (0.67 e^-^/Å^2^) for a total dose of ∼67 e^−^/Å^2^ (see Table S1).

### Image Analysis and 3D Reconstruction for yEMC

#### Overexpressed EMC + Fab in b-DDM

Image processing schematic (**Figure 2 – Figure Supplement 1**) and Supplementary Table 1 have additional details. All dose-fractionated image stacks were corrected for motion artefacts, 2x binned in the Fourier domain, and dose-weighted using MotionCor (Li, 2013) for the DDM datasets, resulting in one dose-weighted and one unweighted integrated image per stack with pixel sizes of 1.22Å (DDM - Polara) or 1.31Å (DDM – Janelia Krios). The parameters of the Contrast Transfer Function (CTF) were estimated using GCTF-v1.06 (Zhang, 2016) and the motion-corrected but unweighted images. For each dataset ∼1000 particles per dataset were manually selected and averaged in 2D using RELION 2.0 (Kimanius, 2016). The resulting class sums were then used as templates for automated particle picking using Gautomatch-v0.55 (Zhang, 2016), followed by extraction in RELION 2.0. Five rounds of 2D classification were performed to eliminate ice contamination, particles near carbon edges, and 2D class without visible secondary structure features. Subsequent particles were subjected to 3D auto-refine in Relion 2.0. The Polara dataset was processed providing a reference model created in Spider (Shaikh, 2008) roughly mimicking the dimensions seen in 2D projections, then a second round was run using the resulting volume before two rounds of 3D classification without alignments. The resulting subset of particles were subjected to 3D auto-refine and then 3D classification with local alignments. The best 83,599 particles were then subjected to 3D refinement resulting in a 3D volume with ∼8 Å reported resolution, which was rescaled and low-pass filtered for use as the reference for the DDM Krios dataset. 3D classification without alignments, 3D refinement, 3D classification with local alignments, and 3D auto refinement were performed resulting in a ∼7 Å structure composed of 170,186 particles. Both resulting reconstructions overlay with one another, despite having Fab DH4 in one sample and DE4 in the other. Furthermore, they both displayed a severe orientation bias, and 3D reconstructions appeared streaky.

Particles from both datasets were re-extracted and scaled to a common pixel size of 1.35 Å and box size of 266. The combined dataset was subjected to two rounds of 3D refinement to form a consensus structure at ∼6.8 Å all conducted in Relion 2.0. These particles were then subjected to 3D refinement in THUNDER (Hu, 2018) using soft-edged mask. THUNDER produced a resulting 3D reconstruction that visually appeared less distorted along the axis of overrepresented views and resulted in a ∼4.8 Å consensus structure. Postprocessing was done in Relion 3.0 resulting in a ∼4.3 Å sharpened map and output was used to generate the FSC plot (**Figure 2 – figure supplement 1**). Molecular graphics and analyses were performed with the UCSF Chimera package (Pettersen, 2004) and Coot 0.8.7 and Coot 0.9 (Emsley, 2004; Emsley, 2010). Local resolution was computed by inputting mask and half maps into Cryosparc 2 local resolution (Stagg, 2014; Punjani, 2017; Punjani, 2019) and visualizing the resulting map and scaling in UCSF Chimera.

#### Endogenous EMC + Fab DH4 in MSP1D1-Yeast Total Extract nanodisc

All dose-fractionated image stacks were corrected for motion artefacts, 2x binned in the Fourier domain, and dose-weighted using MotionCor2 (Zheng, 2017) using Focus (Biyani, 2017) resulting in a 2x binned pixel size of 0.835Å (nanodisc – UCSF Krios). The parameters of the Contrast Transfer Function (CTF) were estimated using GCTF-v1.06 (Zhang, 2016) and the motion-corrected but unweighted images. Data were then split into five groups of 1000 micrographs for processing until they were combined in 3D. Roughly ∼1000 particles per subset were manually selected and averaged in 2D using RELION 3.0 (Zivanov, 2018) for the nanodisc dataset. The resulting class sums were then used as templates for automated particle picking using Autopick in Relion 3.0, followed by extraction and one round of 2D classification per subset to remove ice contamination. The resulting subsets of particles were subject to 3D refinement.

Combining the RELION star files these particles were imported into Cryosparc 2.0 (Punjani, 2017; Punjani, 2019) along with a reference model. These data were subjected to non-uniform homogenous refinement, a round of four class 3D heterogeneous refinement, another round of non-uniform refinement for the best class (roughly 1.2 million particles), non-uniform homogeneous refinement, a round of two class 3D heterogeneous refinement, and another non-uniform homogeneous refinement for the best class (roughly 500,000 particles). These were then exported to RELION 3.0 using PyEM (Asarnow, 2019). 3D Classification was performed with local alignments, then CTF refinement of the best class (230,528 particles) resulting in a ∼3.2 Å final reconstruction. This was post-processed in both RELION 3.0 and using phenix.autosharpen, both resulting maps were used for model building.

### Model building and refinement of yEMC in nanodiscs

Structural biology applications used in this project were compiled and configured by SBGrid (Morin, 2013). The yeast EMC structure was built de novo using Coot (**version 0.8.7 and 0.9**) and UCSF ChimeraX (Goddard, 2018). Visible secondary structure was built by hand for the entire structure using overlays of the yEMC detergent consensus map as well as the yEMC nanodisc unsharpened and sharpened map. Starting with the best resolved transmembrane helices, sequence was placed for each of the predicted transmembrane helices, using TMHMM (Krogh, 2001), in the yEMC proteins. Visual inspection for landmark residues (tryptophan, tyrosine, leucine, and proline) in the sequences that correlated with the position of well densities as well as fit correlation in UCSF Chimera was computed to assign identities for yEMC1, yEMC3, yEMC5, and yEMC6. Connectivity between the EMC1 assigned helix to the lumenal domain was used to start assigning sequence for the lumenal portion of EMC1. Secondary structure prediction was computed for all yEMC proteins using Phyre2 (Kelley, 2015) and Quick2D, a tool within the Max-Plank Institute for Developmental Biology Bioinformatics Toolkit that visualizes several different secondary structure predictors (Jones, 1999; Cuff and Barton, 1999; Ouali and King, 2000; Rost, 2001; Lupas, 1991; Jones, 1994; Ward, 2004; Peng, 2006; Obradovic, 2005). Secondary structure prediction was used to check and guide sequence assignment of beta strands and helices. Next several homology models were computed and overlain for yEMC2, with a predicted TPR structural domain, using Robetta (Raman, 2009; Song, 2013), I-TASSER (Zhang, 2008; Roy, 2010; Yang, 2015), Phyre2 (Kelley, 2015), and RaptorX (Kallberg, 2012). These were used in addition to secondary structure prediction to guide sequence assignment, loop building, and helical packing. Fab DH4 starting structure was computed using Phyre2 1-to-1 threading against a crystal structure of a monoclonal Fab (PDB 1M71, Vyas, 2002). EMC3, EMC5, and EMC6 were built off of the transmembrane helices using sphere refinement, real space refinement, regularization, and visual monitoring of the Ramachandran plot in Coot. EMC7 and EMC10 both form beta sandwich folds on the exterior of the EMC1 lumenal domain, beta strand sequence was placed for both in both densities, position of aromatic residues and loop length differed between the two allowing assignment of each. After building EMC1-3, EMC5-7, and EMC10, there remained several transmembrane helices and a beta strand fitted into the lumen but not connected to EMC1, EMC7, or EMC10. The resolution of the lumenal domain is better than 3 Å in most parts allowing for sequence placement of the EMC4 C-terminus and C-terminal transmembrane helix. The connectivity of the transmembrane helix to the cytoplasmic domain was not resolved. However, there was an additional poorly resolved short helix and loop density in the cytoplasmic domain which was assigned to EMC4.

Two poorly resolved transmembrane helices remained, however due to the fact they did not have clear connectivity to any built strand, poly alanine alpha helices were built in but not assigned to a yEMC protein (**Figure 5 – figure supplement 3**). EMC4 had density in the cytoplasmic domain as well as the lumenal domain, suggesting that it has either one or three transmembrane passes. EMC7 and EMC10 were predicted to have transmembrane helices however the connection between the lumenal densities and those predicted transmembrane helices was not clear.

Additional density that was not built into was visualized in UCSF ChimeraX (Goddard, 2018) and allowed for subsequent assignment of several glycosylated residues and one POPC molecule. Each subunit was built in a separate pdb file and subjected to iterative rounds of phenix.real_space_refine (Adams, 2011; Liebschner, 2019) into segmented maps preceded and followed by adjustment in Coot. Manual assignment of secondary structure restraints was used and improved during Phenix refinement. Once all of the well resolved secondary structure was assigned to yEMC subunits, PDBs were combined and subjected to iterative rounds of phenix.real_space_refine (Adams, 2011; Afonine, 2018; Liebschner, 2019) in the unsharpened and then sharpened maps. Loops were built back where the connectivity was clear and then refined again in Phenix and Coot. PDBs were prepared for refinement steps using phenix.reduce to add hydrogens throughout refinement steps, ReadySet to generate cif restraints, and Phenix PDB preparation tool for creating mmCIF files for deposition. Representative regions of the model as well as the map-to-model FSC can be found in **Figure 2 – Figure Supplement 5**.

### Cryo-EM sample preparation and imaging for hEMC

4 µL of freshly purified hEMC (in detergent or nanodisc) was applied to glow discharged copper Quantifoil holey carbon grids (R1.2/1.3 300 mesh) at 100% humidity and 4 °C in a Vitrobot Mark IV (Thermo) and incubated for 30 seconds. Excess liquid was blotted away with filter paper (blot force 4 – 6, blot time 4 seconds) and the grid plunge-frozen into liquid ethane. Samples were imaged on a FEI Titan Krios microscope operating at 300 kV, equipped with a post-column GIF and a K3 direct detector operating in counting mode. Images were recorded at a nominal magnification of 105,000x (0.8512 Å/pixel at the specimen level) for hEMC in nanodiscs or 81,000x (1.094 Å/pixel at the specimen level) for hEMC in detergent, with target defocus ranging between 0.7 and 2.8 µm and total exposure of ∼70 e/Å^2^ using SerialEM (Mastronarde, 2005). On-the-fly motion correction, CTF estimation and templated particle auto-picking were performed using a pipeline implemented in Focus (Biyani, 2017).

### Cloning and expression constructs for hEMC

A modified version of the biGBac (Weissmann, 2016) multi-gene cloning method was combined with the BacMam (Goehring, 2014) mammalian expression system to allow for recombinant production of human EMC (hEMC). hEMC subunits were individually inserted into pEG, with EMC5 bearing a C-terminal Flag-tag. To amplify gene-expression cassettes (GEC) from pEG, original forward primers from biGBac were used in combination with modified reverse primers bearing complementarity downstream of the SV40 terminator sequence. GECs were inserted into pBIG1a-e vectors as follows: pBIG1a (EMC1 - Uniprot code Q8N766-1), pBIG1b (EMC4 – Q5J8M3-1; EMC5-Flag –Q8N4V1-1, which encodes DYKDDDDK immediately after R131; EMC6 – Q9BV81), pBIG1c (EMC2 – Q15006; EMC3 – Q9P0I2-1; EMC7 – Q9NPA0), pBIG1d (EMC8 – O43402-1; EMC9 – Q9Y3B6), pBIG1e (EMC10 – Q5UCC4-1). These were subsequently combined into pBIG2abcde to yield a single expression vector containing all ten hEMC subunits. Bacmid was generated in DH10 EMBacY *E. coli* and subsequently transfected into Sf9 insect cells using FuGENE (Promega) reagent. Virus was amplified in Sf9 cells up to P3 and virus supernatant sterilized by filtration.

### hEMC expression, purification and nanodisc reconstitution

Recombinant hEMC was expressed by baculovirus transduction of human embryonic kidney (HEK) 293S GnTI-cells grown in suspension. Cells were maintained at 37 °C in Freestyle 293 Expression Medium (Thermo) and expanded with home-made suspension medium (Chaudhary, 2012) in 2 L shaker flasks. For expression of hEMC, 10% (v/v) P3 virus was added to 800 mL of HEK culture at a cell density >3 x 10^6^. 16 hours post-transduction, 10 mM butyrate was added and the temperature reduced to 30 °C. Cells were harvested 48 hours later and stored frozen at -80 °C.

For purification, 15 – 20 g of cell pellet was thawed and resuspended in 60 – 80 mL Lysis Buffer containing 50 mM ammonium citrate pH 6.0, 150 mM sodium chloride, 0.001 mg/mL Benzonase, EDTA-free protease inhibitor cocktail (1 tablet per 50 mL of buffer), and lysed by Dounce homogenization on ice (50 strokes). Glyco-diosgenin (GDN, Anatrace) was added to the lysate at 2% (w/v) and cellular membranes solubilized for 3 hours at 4 °C under constant stirring. Insolubilized material was removed by centrifugation at 100,000 x g, supernatant incubated with 2 mL M2 Flag-affinity resin in-batch for 2 hours at 4 °C. The resin was poured into a column and unbound proteins washed away with 25 column volumes (CV) of Wash Buffer containing 20 mM ammonium citrate pH 6.0, 150 mM sodium chloride, 0.01% (w/v) GDN. Bound hEMC was eluted in 10 CV Wash Buffer containing 0.3 mg/mL Flag peptide and concentrated to < 500 µL using centrifugal concentration filters with 100 kDa cut—off (Amicon).

Sample was polished using size-exclusion chromatography (SEC) on a Superose 6 Increase 10/300 GL column (GE Healthcare) with Running Buffer containing 10 mM ammonium citrate pH 6.0, 100 mM sodium chloride, 0.25 mM TCEP, 0.01% (w/v) GDN. Peak fractions containing hEMC were pooled, concentrated to ∼3 mg/mL and used immediately for cryo-EM grid preparation.

hEMC in nanodiscs composed of MSP1D1 scaffold protein and 1-palmitoyl-2-oleoyl-sn-glycero-3-phosphatidylcholine (POPC) was reconstituted following Flag-affinity chromatography. The MSP1D1 expression vector was a gift from Franz Hagn (TUM, Germany) and the scaffold protein purified from *E. coli* following a published protocol (Hagn, 2018). Prior to reconstitution, hEMC purified by Flag-affinity chromatography was mixed with MSP1D1 and POPC (solubilized as 25 mM stock in 5% n-dodecyl *β*-D-maltoside) in a 1:4:50 ratio and this mixture incubated on ice for 2 hours. Nanodisc reconstitution was achieved by incubation with 0.5 - 1 mL Bio-Beads SM-2 (Bio-Rad) for 16 hours at 4°C under constant rotation. The liquid phase was aspirated, concentrated to < 500 µL and injected onto a Superose 6 SEC column with buffer containing 10 mM ammonium citrate pH 6.0, 100 mM sodium chloride, 0.25 mM TCEP, to separate nanodisc-embedded hEMC from empty nanodiscs. Peak fractions were pooled and concentrated to ∼2 mg/mL for immediate cryo-EM grid preparation.

### Cryo-EM sample preparation and imaging for hEMC

4 µL of freshly purified hEMC (in detergent or nanodisc) was applied to glow discharged copper Quantifoil holey carbon grids (R1.2/1.3 300 mesh) at 100% humidity and 4 °C in a Vitrobot Mark IV (Thermo) and incubated for 30 seconds. Excess liquid was blotted away with filter paper (blot force 4 – 6, blot time 4 seconds) and the grid plunge-frozen into liquid ethane. Samples were imaged on a FEI Titan Krios microscope operating at 300 kV, equipped with a post-column GIF and a K3 direct detector operating in counting mode. Images were recorded at a nominal magnification of 105,000x (0.8512 Å/pixel at the specimen level) for hEMC in nanodiscs or 81,000x (1.094 Å/pixel at the specimen level) for hEMC in detergent, with target defocus ranging between 0.7 and 2.8 µm and total exposure of ∼70 e/Å^2^ using SerialEM (Mastronarde, 2005). On-the-fly motion correction, CTF estimation and templated particle auto-picking were performed using a pipeline implemented in Focus (Biyani, 2017).

### Cryo-EM data processing for hEMC in detergent

Preprocessing in Focus included dose-weighted motion correction using Motioncor2 (Zheng, 2017), CTF estimation using Gctf (Zhang, 2016) and templated autopicking using Gautomatch (Kai Zhang). The autopicking template originated from a reconstruction of hEMC in GDN micelles, with data acquired on a K2 (Gatan) direct electron detector (operated in counting mode) under liquid nitrogen conditions using a Glacios microscope (Thermo) operated at 200 kV. 3713 micrographs with a maximal resolution estimate better than 5 Å were imported into Relion 3.0 (Zivanov, 2018), from which ∼3.35 million particles were extracted applying 4-fold binning. These were subjected to three rounds of 2D classification and two rounds of 3D classification (using the reconstruction obtained from the 200kV dataset as reference), followed by 3D autorefinement. This reconstruction was used as initial model for three rounds of 3D classification of the original ∼3.35 million particles (first round: K = 10, T = 10; second round: K = 10, T = 10; third round: K = 3, T = 16), yielding a set of 144,222 particles.

This set was re-extracted at full pixel size, followed by masked 3D autorefinement, producing a reconstruction at 3.77 Å overall resolution. Application of non-uniform refinement in cryoSPARC (Punjani, 2017; Punjani, 2019) further improved the map quality and overall resolution to 3.60 Å.

### Cryo-EM data processing for hEMC in nanodiscs

Micrographs were preprocessed using Focus in a similar manner as for hEMC in detergent. 9164 micrographs with a maximal resolution estimate better than 5 Å were imported into Relion 3.0, from which ∼5.9 million particles were extracted applying 4-fold binning. These were subjected to three rounds of 3D classification (using hEMC in GDN as reference for the first round), after which 386739 particles were kept and re-extracted to full pixel size. Particles were aligned using global angular search 3D classification (K = 1, T = 4) before one further round of 3D classification with a soft mask and skipping alignment (K = 6, T = 8), to isolate a set of 177560 homogeneous hEMC particles. Masked 3D autorefinement of this particle set yielded a map at 3.6 Å overall resolution. Implementation of cryoSPARC non-uniform refinement led to a consensus map at 3.4 Å global resolution. To aid de novo model building of cytoplasmic and luminal domains, these parts were subjected to masked focused classification (K = 5, T = 8), 3D autorefinement and post-processing in Relion, yielding improved maps at 3.4 Å and 3.2 Å, respectively. To obtain highest quality maps of the transmembrane domains, the 177560 particles from consensus refinement were processed using Sidesplitter (Ramlaul, 2020), producing a 3.3 Å global map after Relion post-processing, where transmembrane helix pitch and side chains were well resolved and allowed for unambiguous sequence assignment. The final particle set was further subjected to 3D variability analysis (Punjani, 2020) in cryoSPARC, revealing the presence or absence of the EMC7 lumenal domain between the EMC1 beta-propellers. Heterogeneous refinement, using a map from 3D variability analysis containing stronger EMC7 density as reference, allowed for further sub-classification of the consensus particle set. Non-uniform refinement of the class containing stronger EMC7 density produced a map at 3.5 Å global resolution, which was subsequently used to build an EMC7 model.

### Model building and refinement of hEMC in nanodiscs and detergent

Given the higher quality hEMC nanodisc map compared to the detergent map, the former was used for de novo model building in Coot (Emsley, 2004; Emsley, 2010). Focused luminal and cytoplasmic, as well as Sideplitter maps, permitted assignment of amino acid sequence throughout all parts of hEMC. Inspection of structural homology and secondary structure predictions for the hEMC subunits produced via HHpred and Quick2D servers (Zimmermann, 2018) predicted the luminal domain of EMC1, the largest hEMC subunit, to consist of two beta-propellers. EMC7 and EMC10 are predicted to feature beta-sandwich structures in the lumen. A final missing beta-strand of the EMC1 membrane proximal propeller could be assigned to the luminal C-terminus of EMC4, which forms a parallel sheet with EMC1 residues 668 – 674. Almost all of EMC2 is predicted to form an alpha-solenoid structure harboring several TPR motifs. Analysis of EMC8 and EMC9 amino acid sequences revealed structural homology to CSN5 (deneddylase subunit of the CSN complex) and Rpn11 (deubiquitinase subunit of the 19S proteasomal regulatory particle) peptide hydrolase folds. The globular density sitting on the distal face of the EMC2 solenoid, facing away from the rest of the complex, was modeled with the EMC8 sequence, which shares ∼45% amino acid sequence identity with EMC9. Additional helical density sitting sideways on top of the EMC2 solenoid could be modeled as two cytoplasmic helices of EMC3 as well as the extended, partially helical meander of the EMC3 C-terminus. Beta-strand-like density on the EMC8 surface, commonly occupied by deubiquitinase substrate peptides, was assigned to the extreme N-terminus of EMC4, with a further downstream part of this cytoplasmic domain snaking along EMC2 and EMC3 towards the transmembrane part of hEMC.

Clear side-chain resolution and excellent connectivity of the Sidesplitter map, within the nanodisc encircled membrane domain, allowed us to model all predicted transmembrane helices of EMC1, EMC3, EMC5 and EMC6. EMC5 extends its C-terminus outside the membrane, which snakes through the central cavity of the EMC2 solenoid on the cytoplasmic side. Inspection of the map at lower thresholds revealed density for at least two additional transmembrane helices facing EMC3 and EMC6 on one side of the complex: continuous density from one of these helices towards the luminal EMC4 C-terminus indicates that at least one of these gate helices represent EMC4’s C-terminal transmembrane helix. However, given poor map resolution and connectivity in this region, we left the other gate helices unassigned.

Model refinement was performed using real-space refinement in Phenix (Adams, 2011), applying secondary structure and Ramachandran restraints. Initially, luminal and cytoplasmic domains were refined individually against their focused maps, after which the improved models were rigid-body placed and refined against the non-uniform refined consensus map. The transmembrane domain was likewise first refined against the Sidesplitter map, after which all parts of hEMC were combined into a consensus model and refined against the consensus map.

The refined hEMC nanodisc model was subsequently docked into the hEMC detergent map, revealing a relative rotation of the entire lumenal domain. The fitted model was manually adjusted in Coot (Emsley, 2004; Emsley, 2010) and refined using Phenix real-space refinement (Adams, 2011). Different masking strategies failed to produce stronger density for the EMC7 lumenal domain in the hEMC detergent maps, despite EMC7 levels being comparable to the other hEMC subunits in subsequent mass spectrometry analysis. EMC7 thus remains absent from our hEMC detergent model, perhaps due to conformational heterogeneity.

### Mass spectrometric analysis of purified hEMC samples in detergent or nanodiscs

GDN solubilized or nanodisc reconstituted hEMC purified by Flag-affinity chromatography and SEC was subjected to mass spectrometric analysis to assess hEMC subunit abundance. For reduction and alkylation of the proteins, proteins were incubated with SDC buffer (1% Sodiumdeoxycholate, 40nmM 2-Cloroacetamide (Sigma-Aldrich), 10 mM tris(2-carboxyethyl) phosphine (TCEP; PierceTM, Thermo Fisher Scientific) in 100 mM Tris, pH 8.0) for 20 min at 37 °C. Before digestion the samples were diluted 1:2 with MS grade water (VWR). Samples were digested overnight at 37 °C with 1 µg trypsin (Promega).

The solution of peptides was then acidified with Trifluoroacetic acid (Merck) to a final concentration of 1% and a pH value of < 2, followed by purification via SCX StageTips (Rappsilber, 2007) washed with 1% TFA in Isopropanol, followed by a second wash with 0.2% TFA, eluted as one fraction with 80% Acetonitrile and 5% Ammonia (Merck).

Samples were vacuum dried and re-suspended in 6 µl of Buffer A (0.1% Formic acid (Roth) in MS grade water (VWR)).

Purified and desalted peptides were loaded onto a 15-cm column (inner diameter: 75 microns; packed in-house with ReproSil-Pur C18-AQ 1.9-micron beads, Dr. Maisch GmbH) via the autosampler of the Thermo Easy-nLC 1000 (Thermo Fisher Scientific) at 50 °C. Using the nanoelectrospray interface, eluting peptides were directly sprayed onto the benchtop Orbitrap mass spectrometer Q Exactive HF (Thermo Fisher Scientific).

Peptides were loaded in buffer A (0.1% (v/v) Formic acid) at 250 nl/min and percentage of buffer B (80% Acetonitril, 0.1% Formic acid) was ramped to 30% over 45 minutes followed by a ramp to 60% over 5 minutes then 95% over the next 5 minutes and maintained at 95% for another 5 minutes. The mass spectrometer was operated in a data-dependent mode with survey scans from 300 to 1650 m/z (resolution of 60000 at m/z =200), and up to 10 of the top precursors were selected and fragmented using higher energy collisional dissociation (HCD with a normalized collision energy of value of 28). The MS2 spectra were recorded at a resolution of 15000 (at m/z = 200). AGC target for MS and MS2 scans were set to 3E6 and 1E5 respectively within a maximum injection time of 100 and 60 ms for MS and MS2 scans respectively. Dynamic exclusion was set to 30ms.

Raw data were processed using the MaxQuant computational platform (Cox, 2008) with standard settings applied. Shortly, the peak list was searched against the reviewed human Uniprot database with an allowed precursor mass deviation of 4.5 ppm and an allowed fragment mass deviation of 20 ppm. MaxQuant by default enables individual peptide mass tolerances, which was used in the search. Cysteine carbamidomethylation was set as static modification, and methionine oxidation and N-terminal acetylation as variable modifications. The iBAQ algorithm was used for calculation of approximate abundances for the identified proteins (Schwanhäusser, 2011) which normalizes the summed peptide intensities by the number of theoretically observable peptides of the protein.

### Sequence alignments

T-coffee PSI-Coffee extension (Notredame, 2000) was used to compute sequence alignments between yEMC, hEMC, and homologous proteins (**Figure 1 – figure supplement 6-7, Figure 3 – figure supplement 3, Figure 5 – figure supplement 4**). Outputs of these alignments were visualized in Jalview (Waterhouse, 2009) for figure creation and colored by ClustalX convention.

### Figure and video creation

All figures were assembled and edited in Adobe Illustrator. **Figure 1 and Figure 1 – figure supplement 3** were created using BioRender. All of the visualization, structure figures, and structure videos were made using UCSF ChimeraX 1.0 (Goddard, 2018) and UCSF Chimera 1.14 (Pettersen, 2004). Flow cytometry plots were generated in Python and labeled in Adobe Illustrator.

## Acknowledgements

We thank J. Weibazahn, P. Walter, R. Irannejad, J. Gestwicki, R. Scheltema, Ö. Karayel, H. Nguyen, I. Johnson, N. Talledge, L. Kenner, E. Thompson, K. Hickey, J. Kellermann, S. von Gronau, M. Feige, K. Swain, M. Liao, C-W. Lee, F. Wilfling, and members of the Weissman, Frost, and Schulman laboratories for assistance and helpful discussion; L. Metzger, Z Roe-Zurz, M. Tessema, D.W. Chester, and S. Aller for assisting with fermentation; M. Sun, H. Autzen, and E. Green for advice on nanodisc reconstitution; F. Hagn and I. Goba for the gift of the MSP1D1 vector and advice on nanodisc reconstitution; P. Thomas and D. Asarnow for computational support; M. Braunfeld, G. Gilbert, E. Tse, D. Bulkley, M. Harrington, A. Myasnikov and Z. Yu of the UCSF Center for Advanced CryoEM for microscopy support and funded by NIH grants S10OD020054 and 1S10OD021741; J. Baker-LePain and the QB3 shared cluster (NIH grant 1S10OD021596-01) for computational support; and the Howard Hughes Medical Institute (HHMI); Z. Yu and H. Chou of the CryoEM Facility at the HHMI Janelia Research Campus (NIH grant 1S10OD021596-01). D. Bollschweiler, T. Schäfer and the cryo-EM facility at the Max Planck Institute of Biochemistry; B. Steigenberger, the mass spectrometry core facility at the Max Planck Institute of Biochemistry; E. Gouaux for the gift of the pEG vector; A Titan X Pascal used for this research was donated by the NVIDIA Corporation. This study was supported in part by the HDFCCC Laboratory for Cell Analysis Shared Resource Facility through a grant from NIH (P30CA082103). Molecular graphics and analyses performed with UCSF ChimeraX, developed by the Resource for Biocomputing, Visualization, and Informatics at the University of California, San Francisco, with support from National Institutes of Health R01-GM129325 and the Office of Cyber Infrastructure and Computational Biology, National Institute of Allergy and Infectious Diseases. UCSF Chimera is developed by the Resource for Biocomputing, Visualization, and Informatics at the University of California, San Francisco (supported by NIGMS P41-GM103311).

## Data and materials availability

All data needed to evaluate the conclusions in the paper are present in the paper and/or the supplementary materials, and the structural data will be made available in EMPIAR, PDB, and EMDB public databases upon publication. All of the raw data files will be in the supplemental data files and code for evaluating will available at GITHUB (MMMMMMM). Accession numbers are as follows: hEMC nanodisc map (EMD- and model (PDB-); hEMC detergent map (EMD-) and model (PDB-); yEMC nanodisc map (EMD-) and model(PDB-); yEMC detergent map (EMD-).

## Author contributions

Conception and design, analysis and interpretation of data: L.E.M.V., B.B., K.D.P, N.T.S.O, J.L.B., J.R.P., E.A.B., N.S., M.J.S., R.M.S., C.S.C, B.A.S, A.F., and J.S.W.; acquisition of data: L.E.M.V., B.B., K.D.P., N.T.S.O., J.L.B., J.R.P ., E.B., and N.S.; writing (original draft): L.E.M.V., B.B., and K.D.P.; writing (review and editing): L.E.M.V., B.B., K.D.P, N.T.S.O, J.L.B., J.R.P., E.B., N.S., M.J.S., R.M.S., C.S.C, B.A.S, A.F., and J.S.W

