## Supplemental Figures for "Structural and mechanistic basis of the EMC-dependent biogenesis of distinct transmembrane clients"

#### Table of Contents

| Item | Page |
|---|---|
| *Supplementary Figures* | |
| Figure 1 – figure supplement 1: **Purification of yEMC.** | 4 |
| Figure 1 – figure supplement 2: **Purification of recombinant hEMC.** | 5 |
| Figure 1 – figure supplement 3: **Overview of strain creation and functional assays.** | 6 |
| Figure 1 – figure supplement 4: **Western blots for EMC1 and EMC2.** | 7 |
| Figure 1 – figure supplement 5: **Western blots for EMC3 and EMC5.** | 8 |
| Figure 1 – figure supplement 6: **Amino acid conservation of EMC1.** | 9 |
| Figure 1 – figure supplement 7: **Amino acid conservation of EMC2, EMC3, EMC5.** | 10 |
| Figure 1 – figure supplement 8: **Genotyping of ten mutants** | 11 |
| Figure 2 – figure supplement 1: **Cryo-EM Reconstruction of yEMC.** | 12-13 |
| Figure 2 – figure supplement 2: **Cryo-EM Reconstruction of hEMC.** | 14 |
| Figure 2 – figure supplement 3: **Cryo-EM data processing workflow for yEMC.** | 15 |
| Figure 2 – figure supplement 4: **Cryo-EM data processing workflow for hEMC.** | 16 |
| Figure 2 – figure supplement 5: **yEMC cryo-EM map validation.** | 17 |

Figure 2 – figure supplement 6: **hEMC cryo-EM map validation.** 18

Figure 2 – figure supplement 7: **Comparison of yeast and human EMC subunits.** 19

Figure 3 – figure supplement 1: **Flow cytometry for mutations in the EMC cytoplasmic domain.** 20-21

Figure 3 – figure supplement 2: **Additional flow cytometry for mutations in the EMC cytoplasmic domain.** 22-23

Figure 3 – figure supplement 3: **Both EMC8 and EMC9 can be fitted into the hEMC cryo-EM maps.** 24

Figure 5 – figure supplement 1: **Flow cytometry of gated cavity mutants.** 25-26

Figure 5 – figure supplement 2: **Additional flow cytometry of gated cavity mutants.** 27-28

Figure 5 – figure supplement 3: **Ambiguity in assignment of flexible gate helices.** 29

Figure 5 – figure supplement 4: **Comparison of EMC3 to YidC-family members.** 30

Figure 6 – figure supplement 1: **Resolved lipid densities in hEMC and yEMC nanodisc maps.** 31

Figure 6 – figure supplement 2: **Flow cytometry of lipid-filled cavity mutants.** 32-33

Figure 6 – figure supplement 3: **Additional flow cytometry of lipid-filled cavity mutants.** 34-35

Figure 7 – figure supplement 1: **Conformational heterogeneity of the hEMC lumenal domain between detergent and nanodisc maps.** 36

Figure 7 – figure supplement 2: **Flow cytometry of lumenal domain mutants.** 37-38

Figure 7 – figure supplement 3: **Additional flow cytometry of lumenal domain mutants.** 39-40

*Supplementary Tables and other files*

Supplemental Table 1: **Cryo-EM data acquisition, reconstruction and model refinement statistics.** 41

Supplemental Table 2: **Reagents (available upon request).**
**Reagents created and purchased for this study as described in the methods section of this study**

Supplemental Table 3: **Source Data 1 – Figure 1 – Figure Supplement 2 (available upon request).**
**EMC subunit abundances assessed by mass spectrometry and iBAQ analysis**
SEC purified hEMC in detergent (sheet 1) or nanodiscs (sheet 2) were subjected to tryptic digestion and mass spectrometry. The tables list identified proteins sorted by iBAQ score (descending order).

Supplemental File 1: **Source Data 2 – Figures 2-7, Video 1-3**
**hEMC nanodisc Validation Report (available upon request).**

Supplemental File 2: **Source Data 3 – Figures 2-7, Video 1-3**
**hEMC detergent Validation Report (available upon request).**

Supplemental File 3: **Source Data 4 – Figures 2-7, Video 1-3**
**yEMC nanodisc Validation Report (available upon request).**

Supplemental File 4: **Source Data 5 – Figures 2-7, Video 1-3 (available upon request).**
**yEMC & hEMC models and maps for evaluation of map to model fit**
ChimeraX 1.0 session with yEMC nanodisc model, yEMC nanodisc unsharpened consensus map, yEMC nanodisc sharpened consensus map, yEMC DDM unsharpened consensus map, yEMC DDM sharpened consensus map, hEMC detergent model, hEMC detergent unsharpened consensus map, hEMC detergent sharpened consensus map, hEMC nanodisc (ND) model, hEMC ND unsharpened consensus map, hEMC ND sharpened consensus map. Separate map, model, focused maps, metadata files, a Coot session, and/or a Pymol session are all available upon request. PDB and EMDB uploads in progress. Raw data will be made available in the EMPIAR database after publication (or concurrent if possible).

**FIGURE 1 - figure supplement 1**

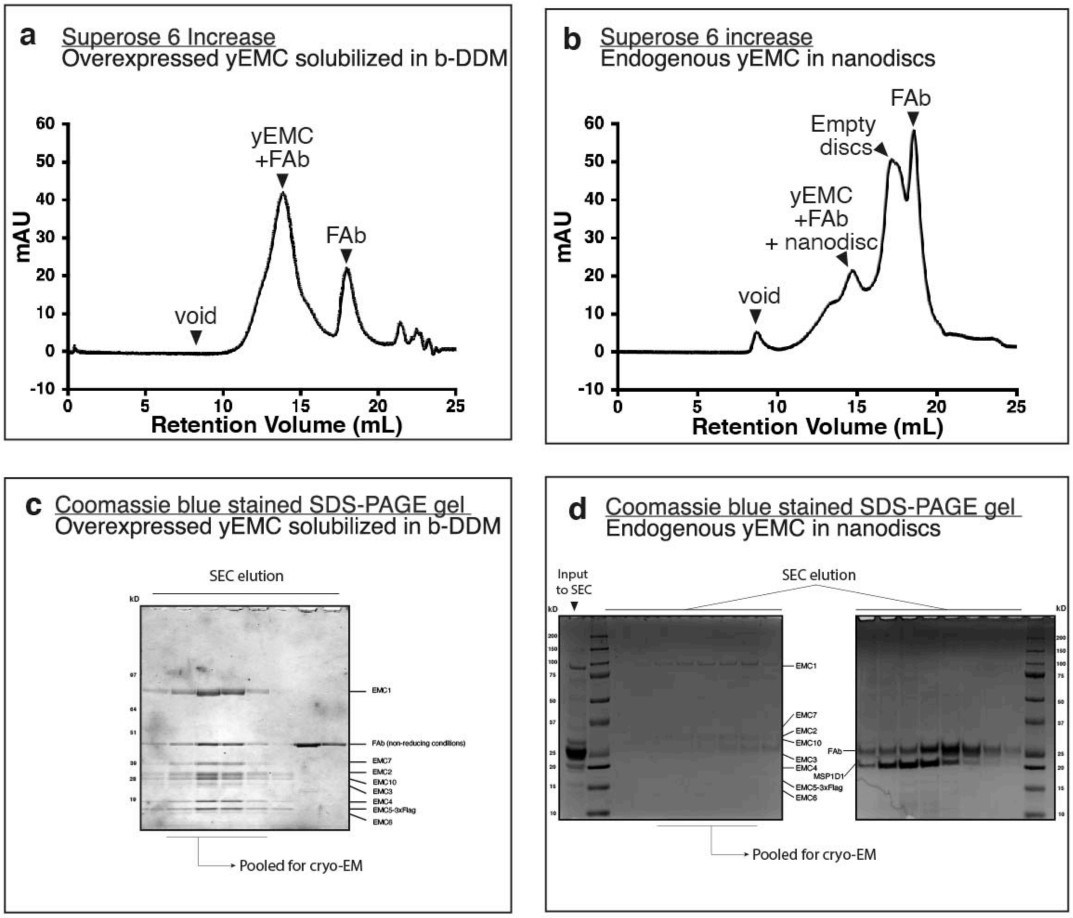

Figure 1 – figure supplement 1: **Purification of yEMC.**

a) Size-exclusion chromatography (SEC) purification of overexpressed yEMC+FAb in DDM detergent.
b) SEC purification of endogenous yEMC + FAb in MSP1D1 nanodisc.
c) Coomassie-stained SDS-PAGE analysis of SEC elution fractions.
d) Coomassie-stained SDS-PAGE analysis of SEC elution fractions.

### FIGURE 1 - figure supplement 2

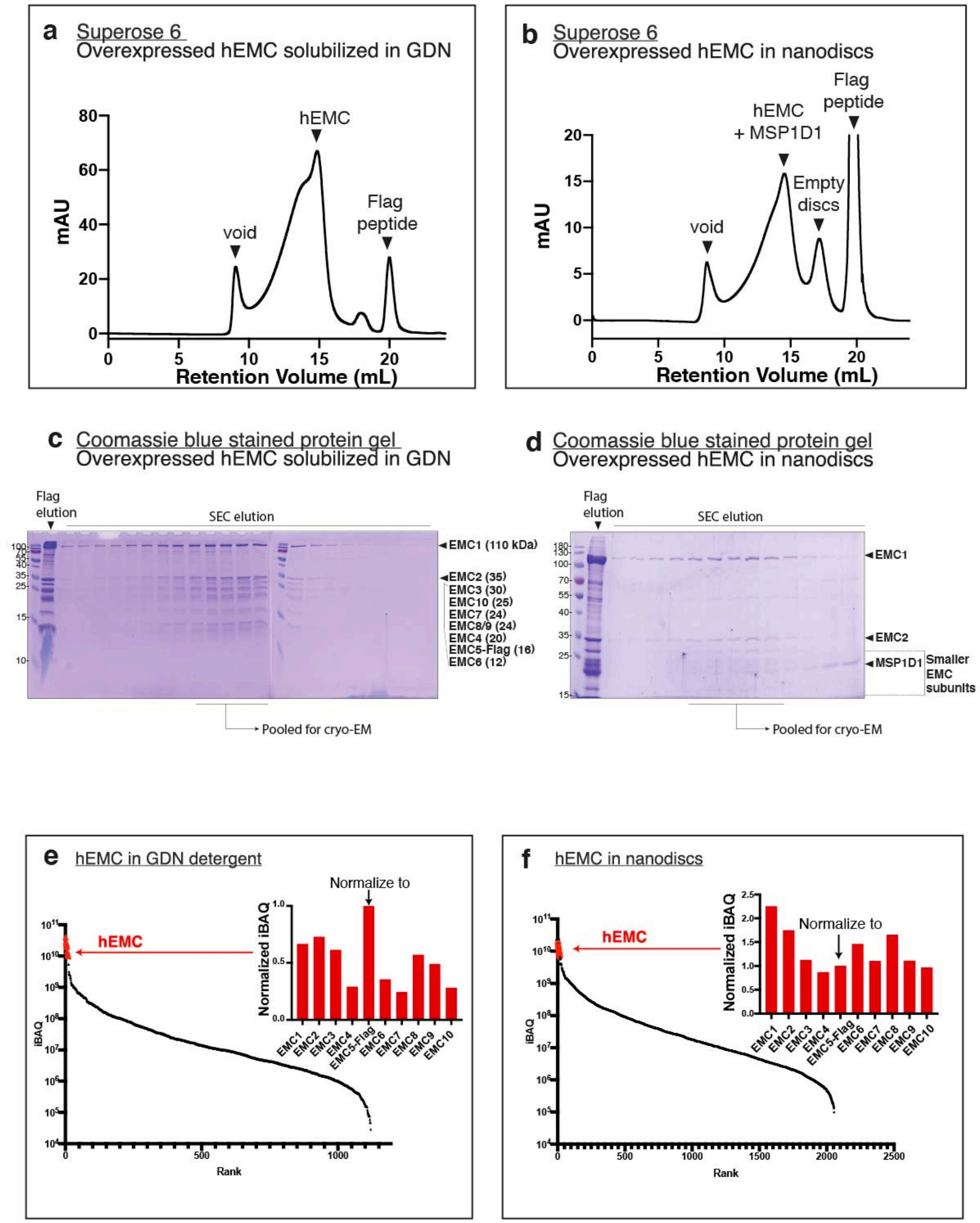

<u>Figure 1 – figure supplement 2</u>: **Purification of recombinant hEMC.**

a) Size-exclusion chromatography (SEC) purification of hEMC in GDN detergent.
b) SEC purification of hEMC reconstituted in MSP1D1 nanodiscs.
c) Coomassie-stained SDS-PAGE analysis of the SEC elution fractions from a).
d) Coomassie-stained SDS-PAGE analysis of the SEC elution fractions from b).
e) Mass spectrometry analysis of purified hEMC in GDN following SEC. iBAQ values for identified proteins in the sample are sorted in descending order along the X-axis. hEMC subunits form a cluster (red) and their normalized iBAQ values (against EMC5-Flag) are shown in the inset.
f) As in c) for hEMC in nanodiscs.

**FIGURE 1 - figure supplement 3**

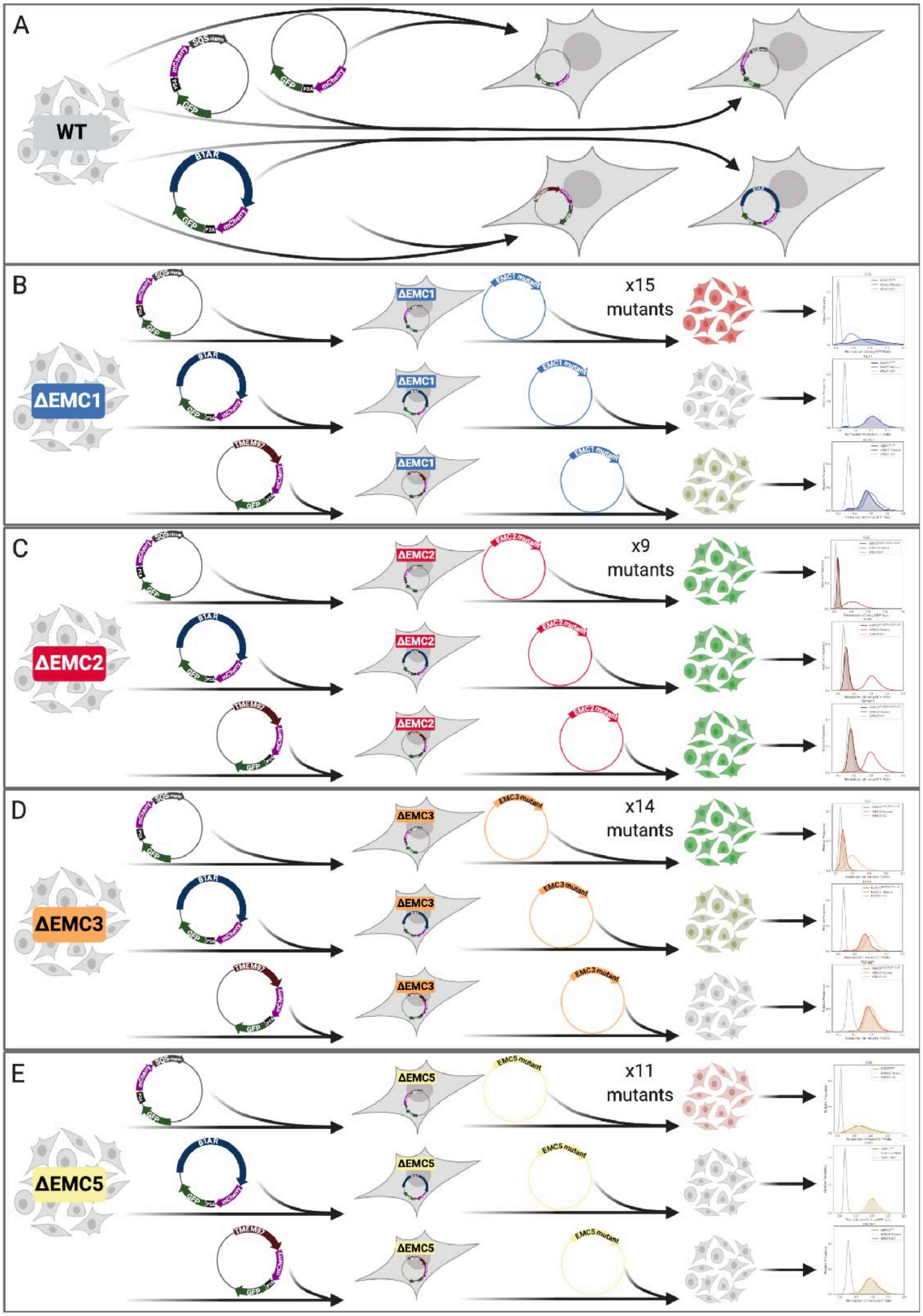

<u>Figure 1 – figure supplement 3</u>: **Overview of cell line generation and functional assays.**

#### FIGURE 1 - figure supplement 4

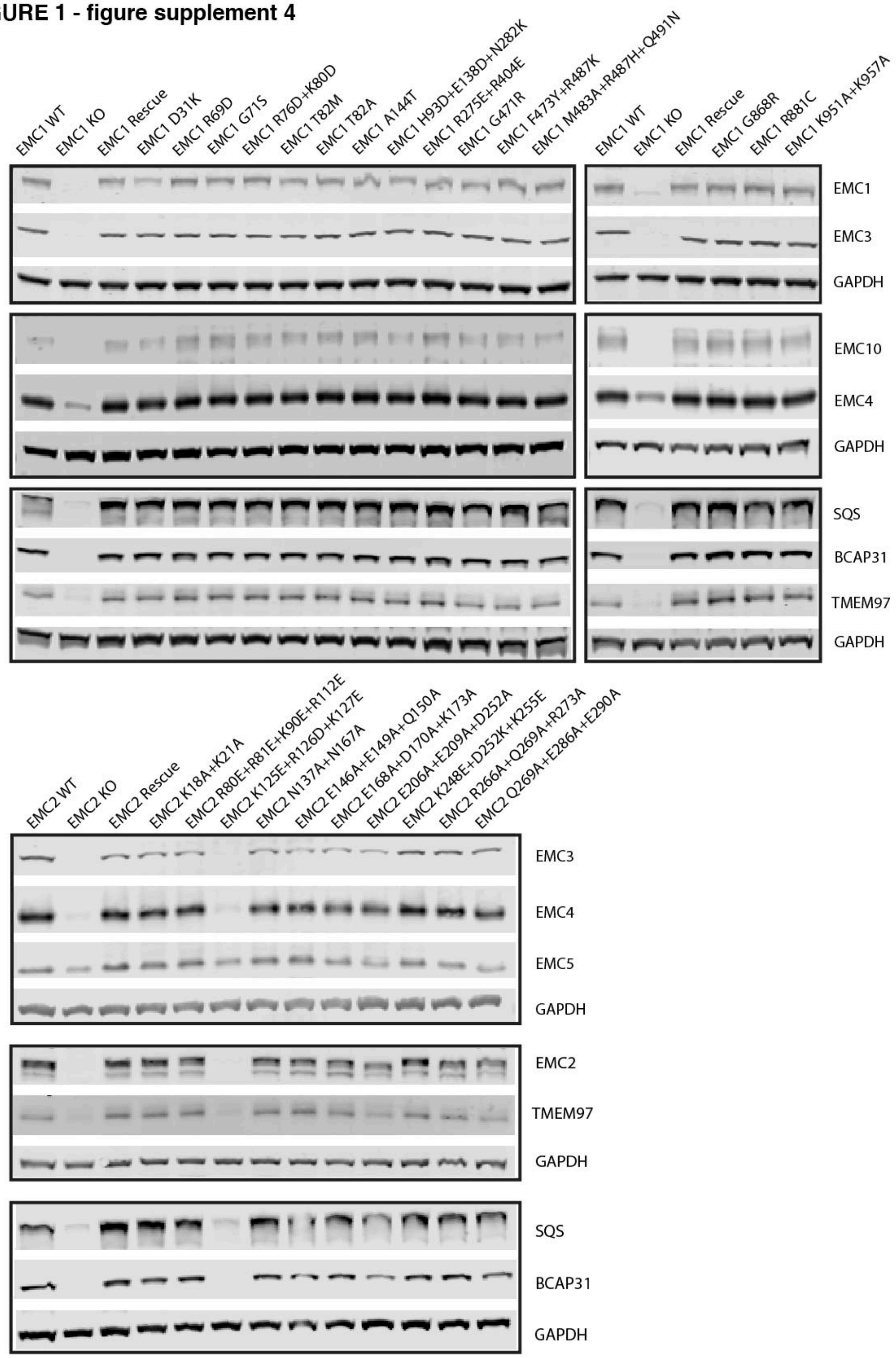

<u>Figure 1 – figure supplement 4</u>: **Western blots for EMC1 and EMC2.**
Human functional assay to check for complex stability. For each mutant, abundance of several EMC subunits as well as several representative client proteins was tested.

#### FIGURE 1 - figure supplement 5

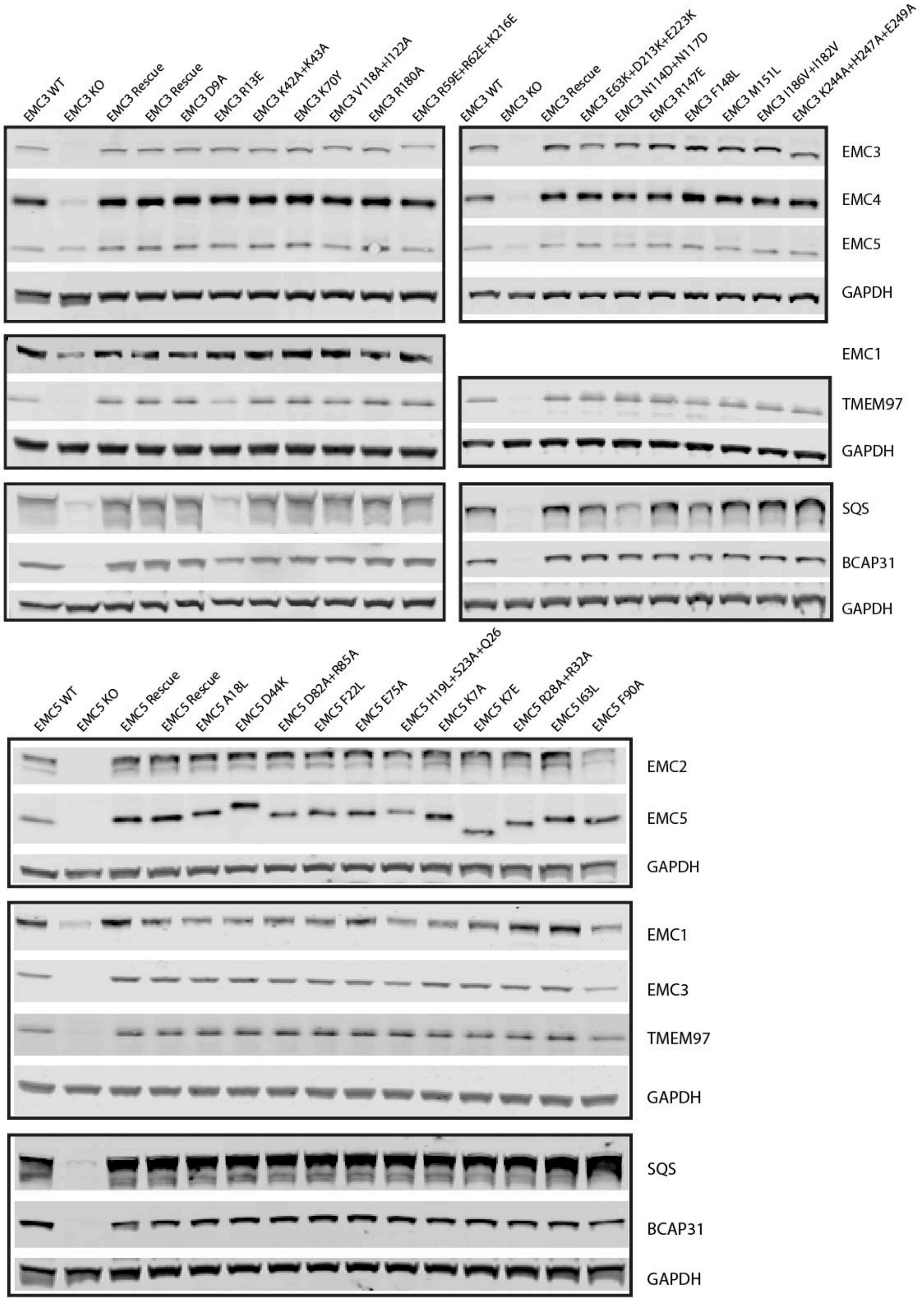

Figure 1 – figure supplement 5: **Western blots for EMC3 and EMC5.**
Human functional assay to check for complex stability. For each mutant, abundance of several EMC subunits as well as several representative client proteins was tested.

#### FIGURE 1 - figure supplement 6

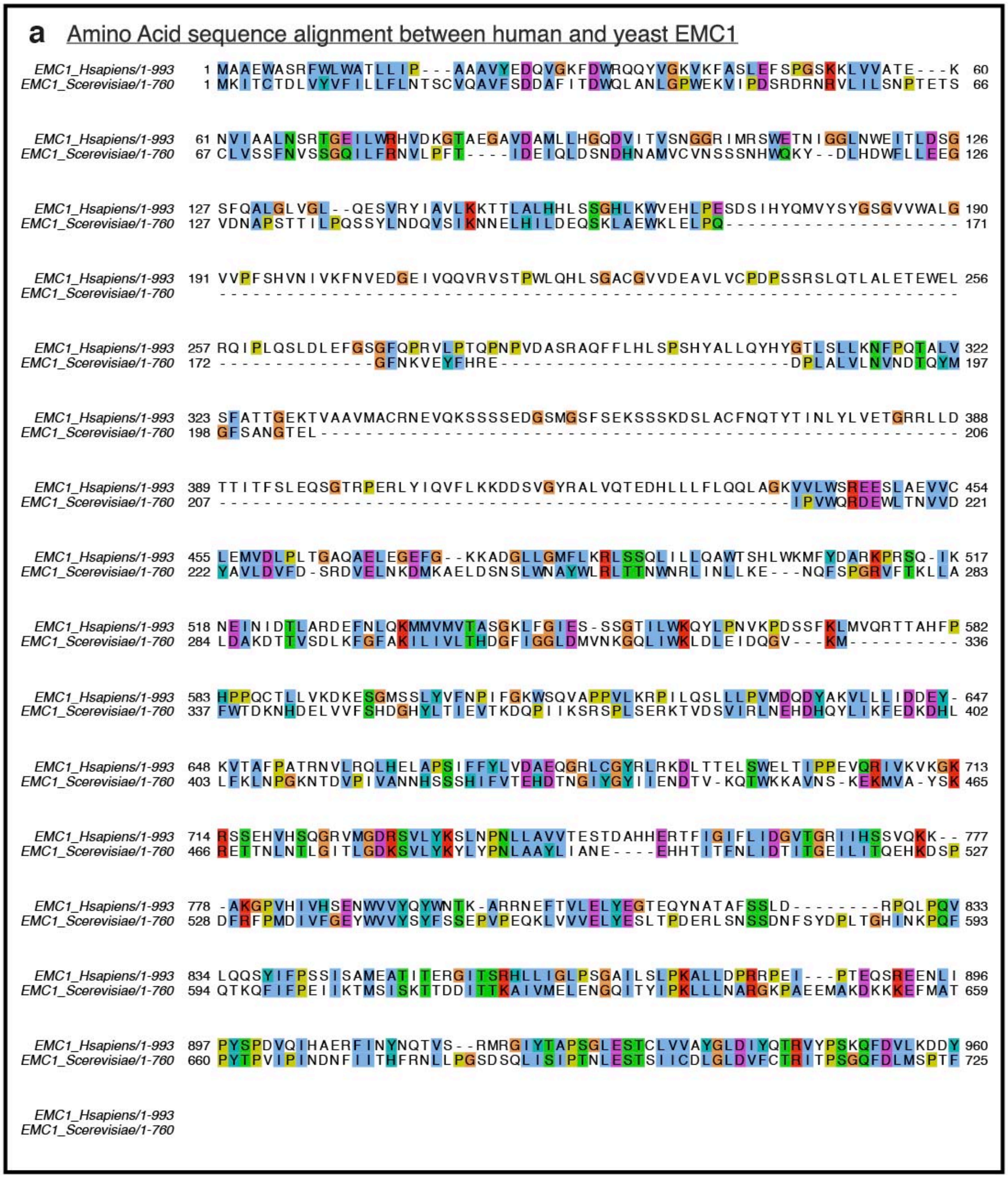

Figure 1 – figure supplement 6: **Amino acid conservation of EMC1.**
Yeast to human sequence alignment of EMC1 by t-Coffee. Colored by ClustalX coloring.

### FIGURE 1 - figure supplement 7

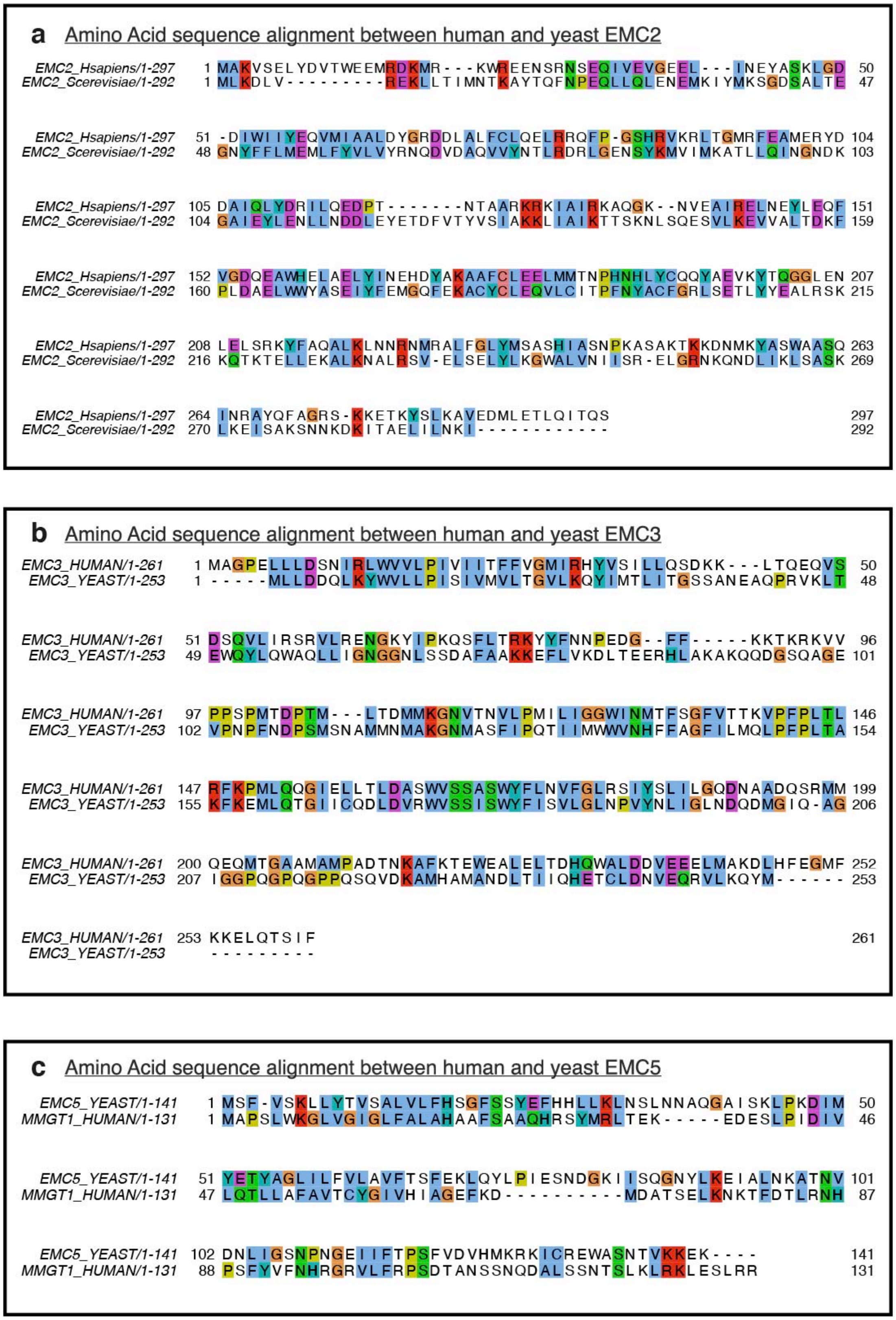

Figure 1 – figure supplement 7: **Amino acid conservation of EMC2, EMC3, EMC5.**
a-c) Yeast to human sequence alignment of EMC2, EMC3, and EMC5. Computed by t-Coffee and colored by ClustalX coloring.

**FIGURE 1 - figure supplement 8**

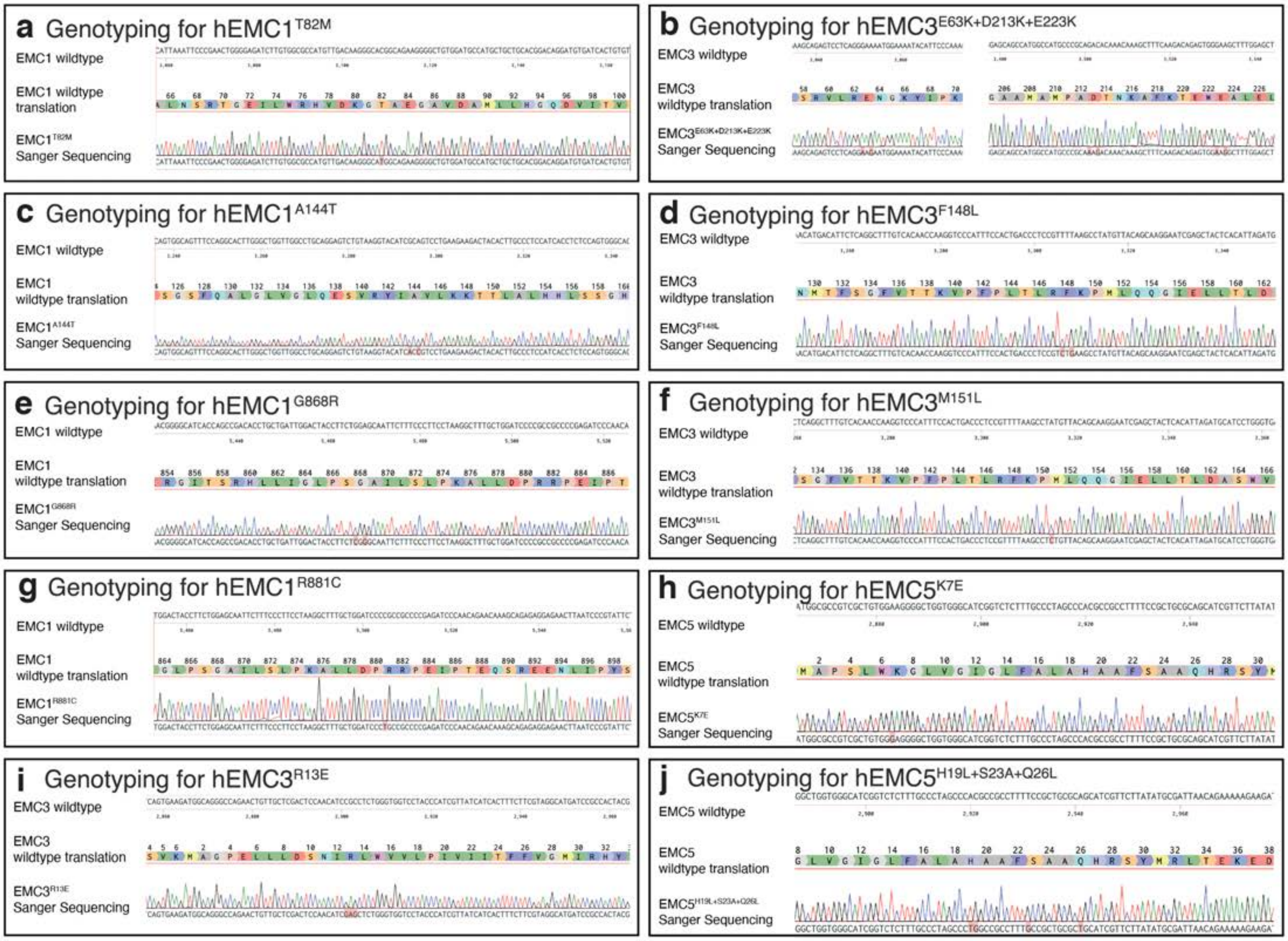

Figure 1 – figure supplement 8: **Genotype confirmation for a subset of mutants.**
a) Sequencing of mutant aligned to wild type sequence for hEMC1 T82M
b) Sequencing of mutant aligned to wild type sequence for hEMC1 E63K+D213K+E223K
c) Sequencing of mutant aligned to wild type sequence for hEMC1 A144T
d) Sequencing of mutant aligned to wild type sequence for hEMC3 F148L
e) Sequencing of mutant aligned to wild type sequence for hEMC1 G868R
f) Sequencing of mutant aligned to wild type sequence for hEMC3 M151L
g) Sequencing of mutant aligned to wild type sequence for hEMC1 R881C
h) Sequencing of mutant aligned to wild type sequence for hEMC5 K7E
i) Sequencing of mutant aligned to wild type sequence for hEMC3 R13E
j) Sequencing of mutant aligned to wild type sequence for hEMC5 H19L+S23A+Q26L

#### FIGURE 2 - figure supplement 1

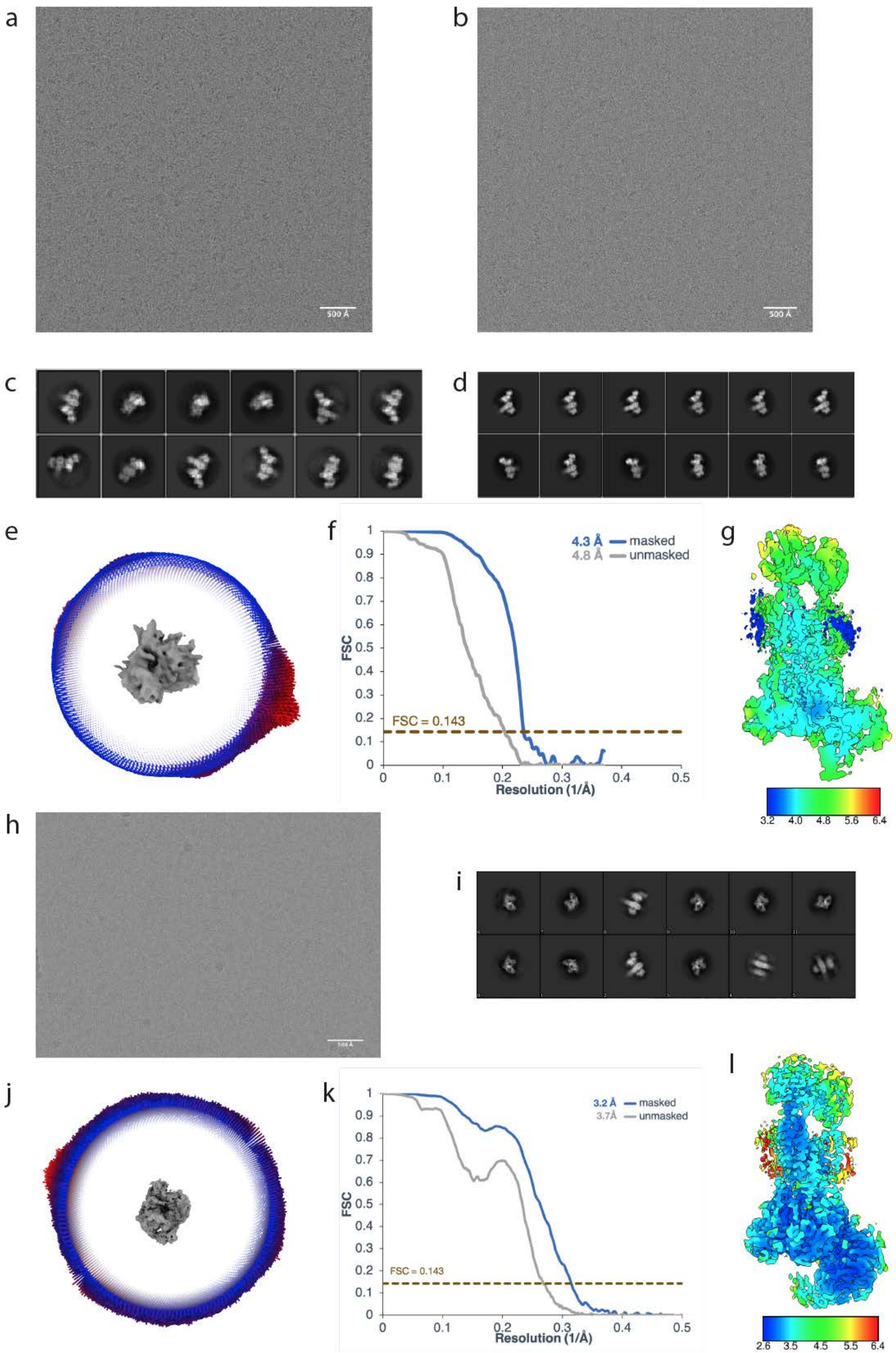

(Legend on next page)

Figure 2 – figure supplement 1: **Cryo-EM Reconstruction of yEMC.**

a) Representative motion-corrected micrograph for yEMC-FabE in DDM. Scale-bar = 500 Å
b) Representative motion-corrected micrograph for yEMC-FabH in DDM. Scale-bar = 500 Å
c-d) Gallery of 2D classes for the final consensus particle set of yEMC in DDM.
e) Angular distribution of the final consensus particle set of yEMC in DDM.
f) Gold-standard Fourier shell correlation (FSC) of the consensus yEMC DDM map. The FSC at 0.143 is indicated by a gold line.
g) A slice through the consensus yEMC DDM map colored by local resolution.
h) Representative motion-corrected micrograph for yEMC in nanodiscs. Scale-bar = 500 Å
i) Gallery of 2D classes of yEMC in nanodiscs.
j) Angular distribution of the final consensus particle set of yEMC in nanodiscs.
k) Gold-standard Fourier shell correlation (FSC) of the consensus yEMC nanodisc map. The FSC at 0.143 is indicated by a gold line.
l) A slice through the consensus yEMC nanodisc map colored by local resolution.

**FIGURE 2 - figure supplement 2**

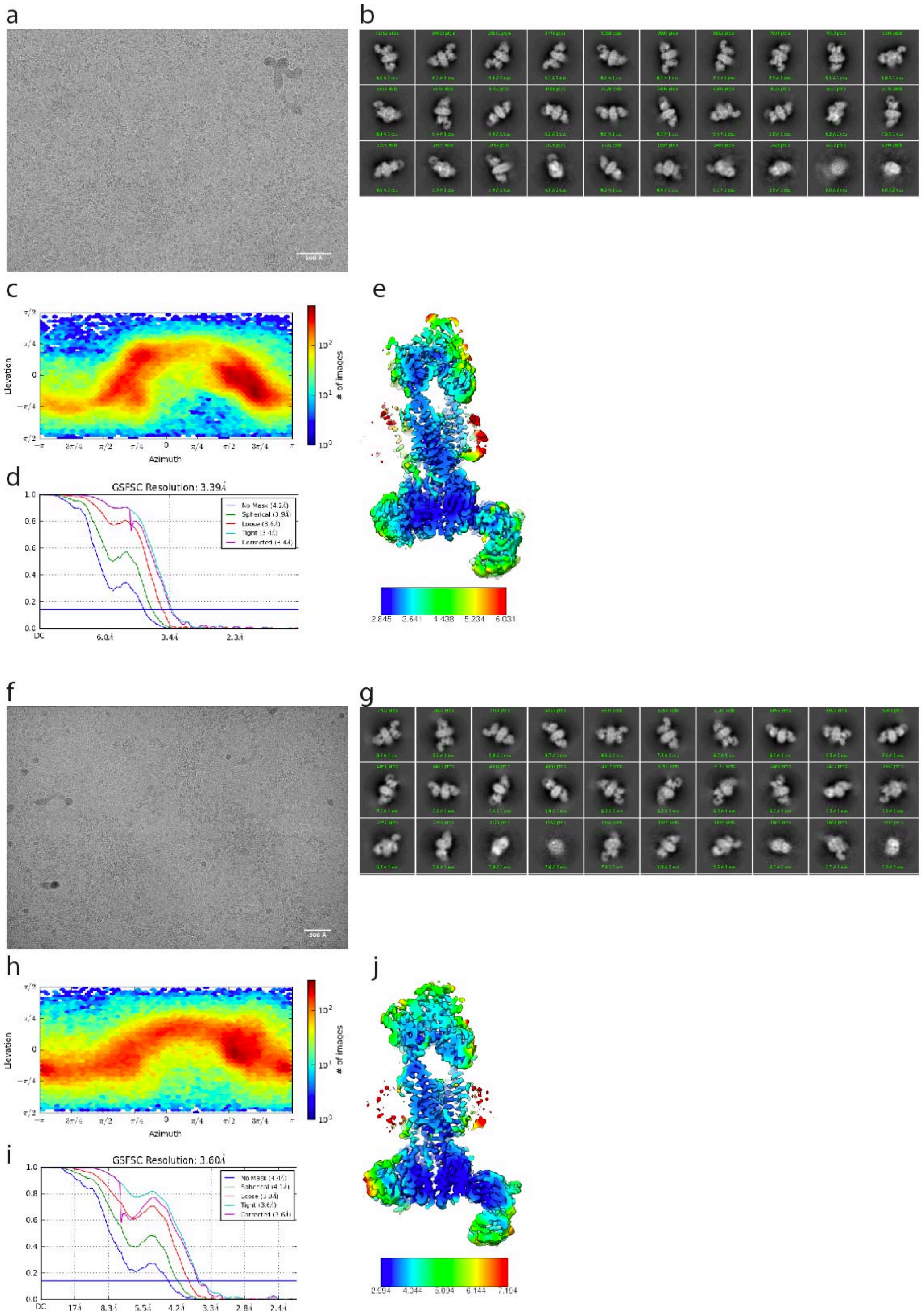

<u>Figure 2 – figure supplement 2:</u> **Cryo-EM Reconstruction of hEMC.**

a) Representative motion-corrected micrograph for hEMC in nanodiscs. Scale-bar = 500 Å
b) Gallery of 2D classes for the final consensus particle set of hEMC in nanodiscs.
c) Angular distribution of the final consensus particle set of hEMC in nanodiscs.
d) Gold-standard Fourier shell correlation (FSC) of the consensus hEMC nanodisc map. The FSC at 0.143 is indicated by a blue line.
e) A slice through the consensus hEMC nanodisc map colored by local resolution.
f - j) As for a – e) but for hEMC in detergent.

#### FIGURE 2 - figure supplement 3

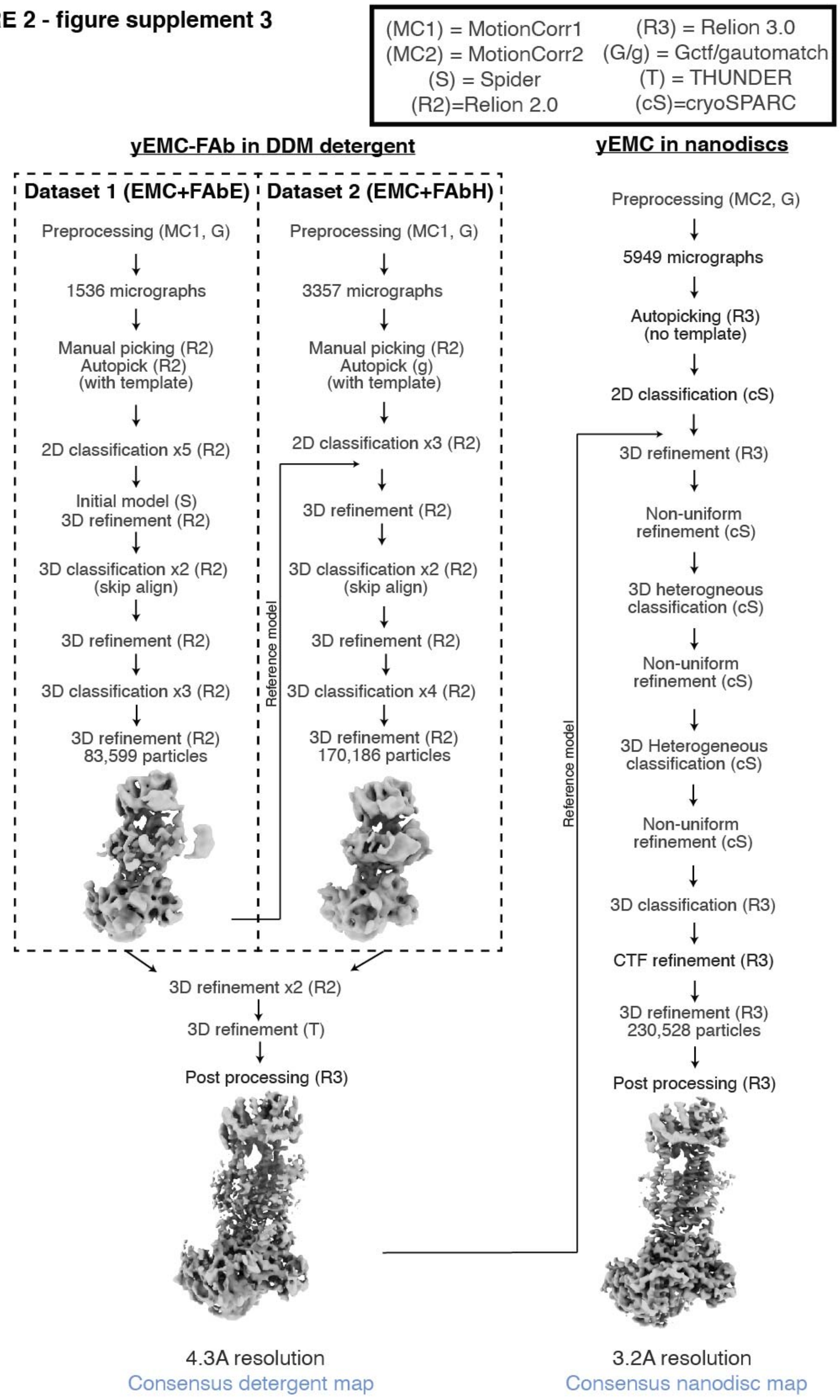

Figure 2 – figure supplement 3: **Cryo-EM data processing workflow for yEMC.**

#### FIGURE 2 - figure supplement 4

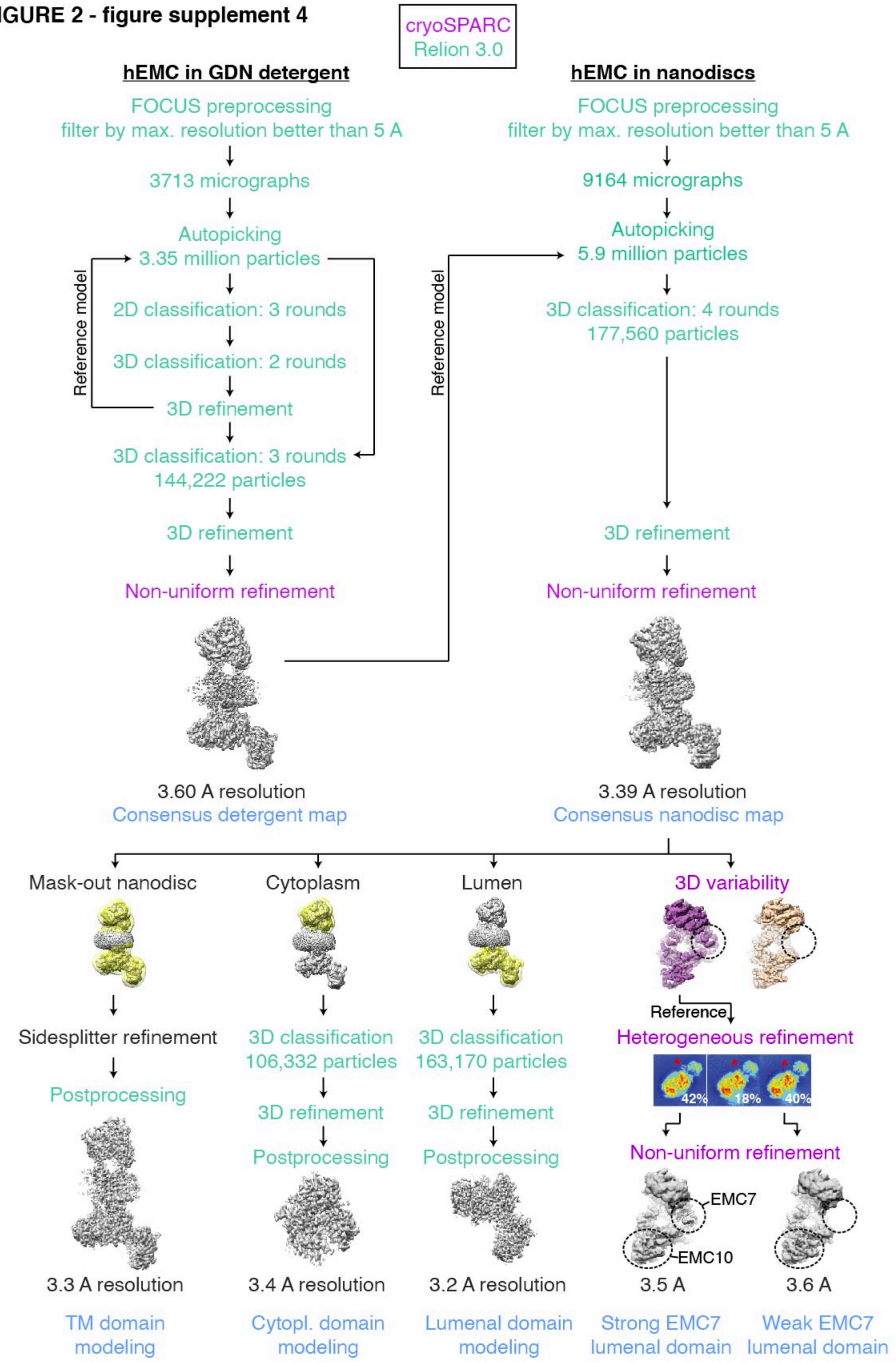

<u>Figure 2 – figure supplement 4:</u> **Cryo-EM data processing workflow for hEMC.**

**FIGURE 2 - figure supplement 5**

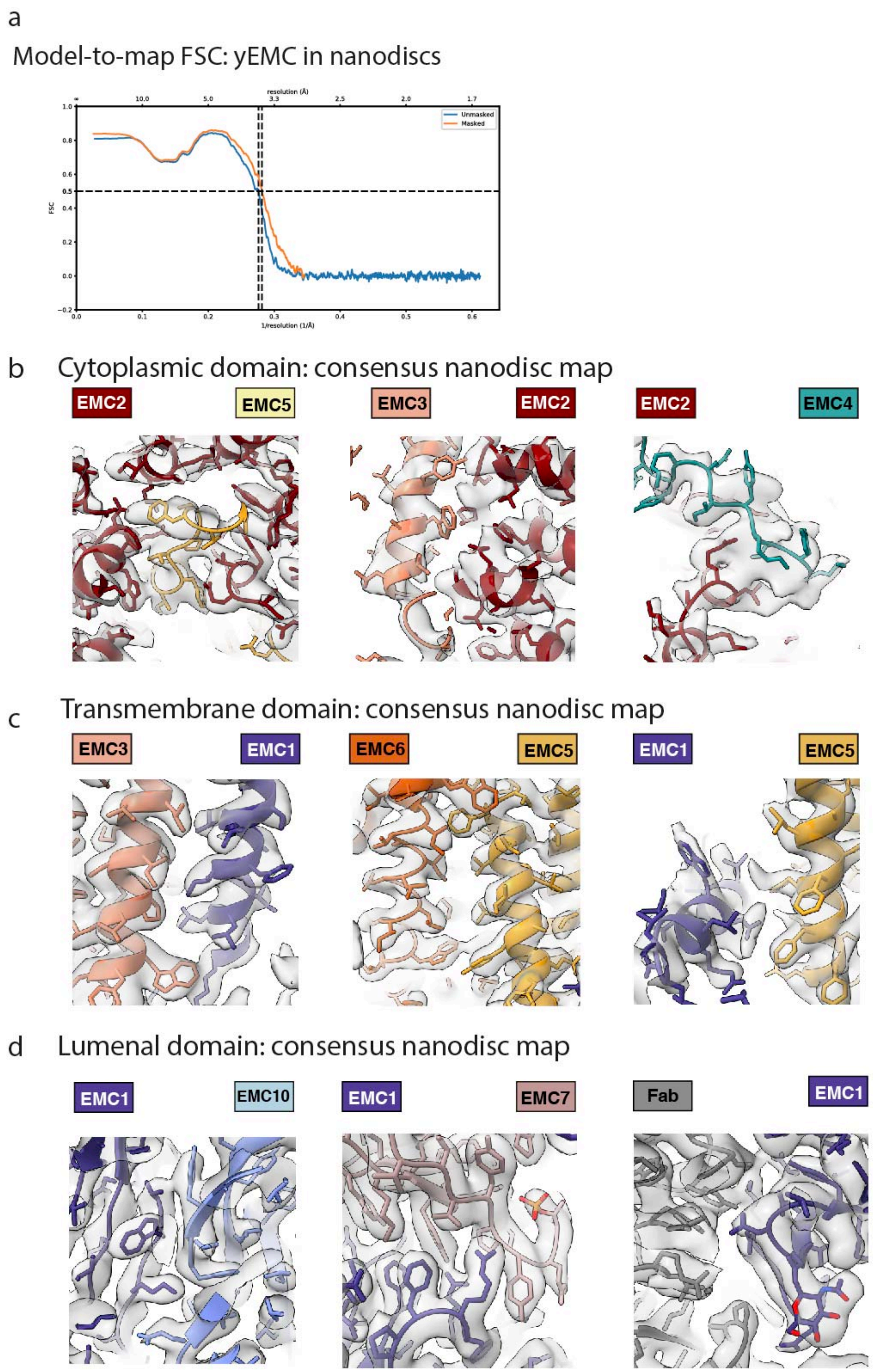

Figure 2 – figure supplement 5: **yEMC cryo-EM map validation.**

a) Final model-to-map FSC curve shown for yEMC in nanodiscs
b) Consensus yEMC nanodisc density shown superposed on the final yEMC nanodisc model. Three cytoplasmic subunit interfaces are depicted.
c) As in b), for transmembrane subunit interfaces.
d) As in b), for lumenal subunit interfaces.

#### FIGURE 2 - figure supplement 6

Figure 2 –

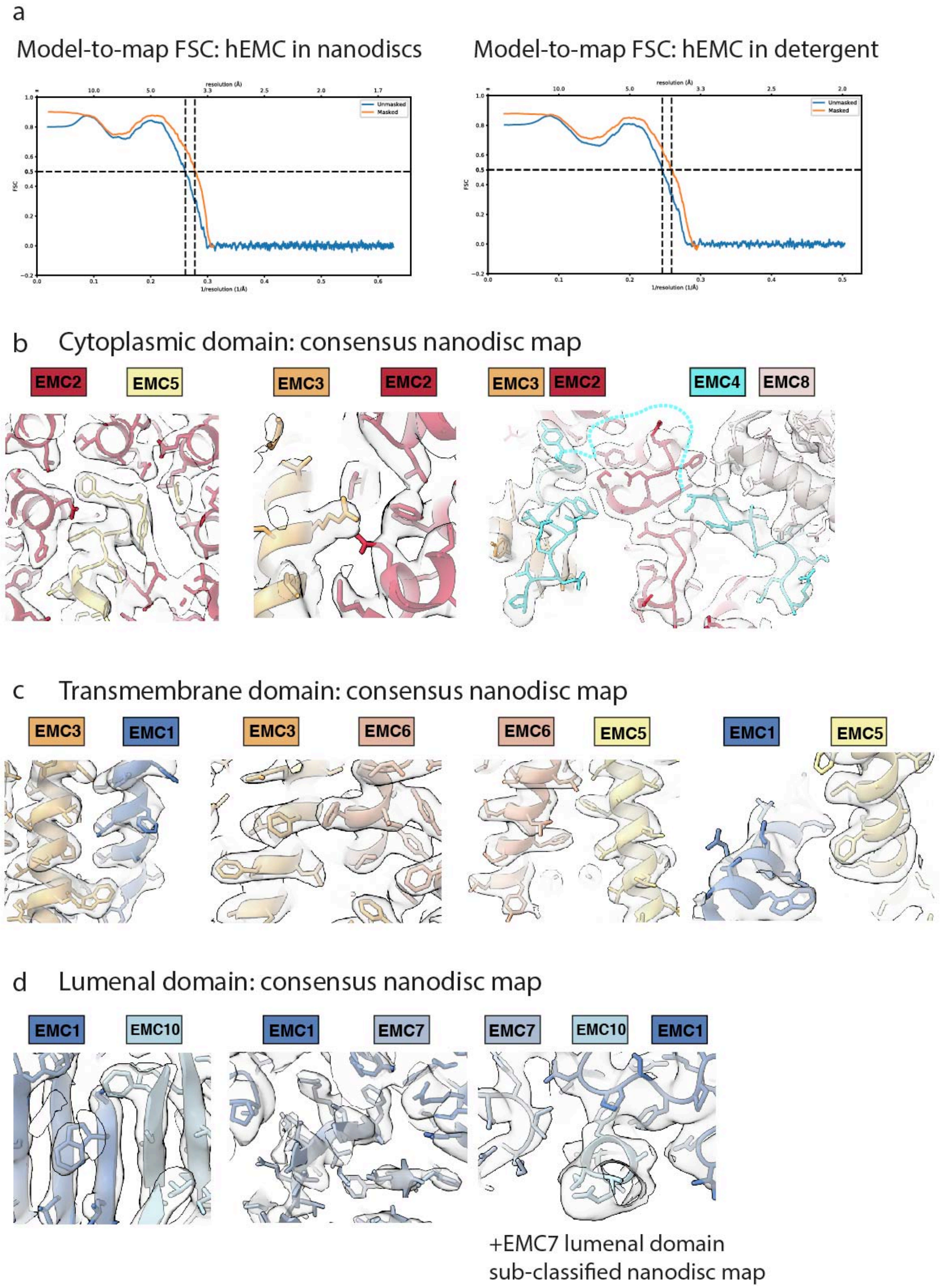

Figure 2 – figure supplement 6: **hEMC cryo-EM map validation.**

e) Final model-to-map FSC curves shown for hEMC in nanodiscs (left) and detergent (right).

f) Consensus hEMC nanodisc density shown superposed on the final hEMC nanodisc model. Three cytoplasmic subunit interfaces are depicted. Cyan dashed lines correspond to an unresolved cytoplasmic loop of EMC4.

g) As in b), for transmembrane subunit interfaces.

h) As in b), for lumenal subunit interfaces. The right most panel depicts the sub-classified hEMC nanodisc map featuring stronger EMC7 lumenal density.

#### FIGURE 2 - figure supplement 7

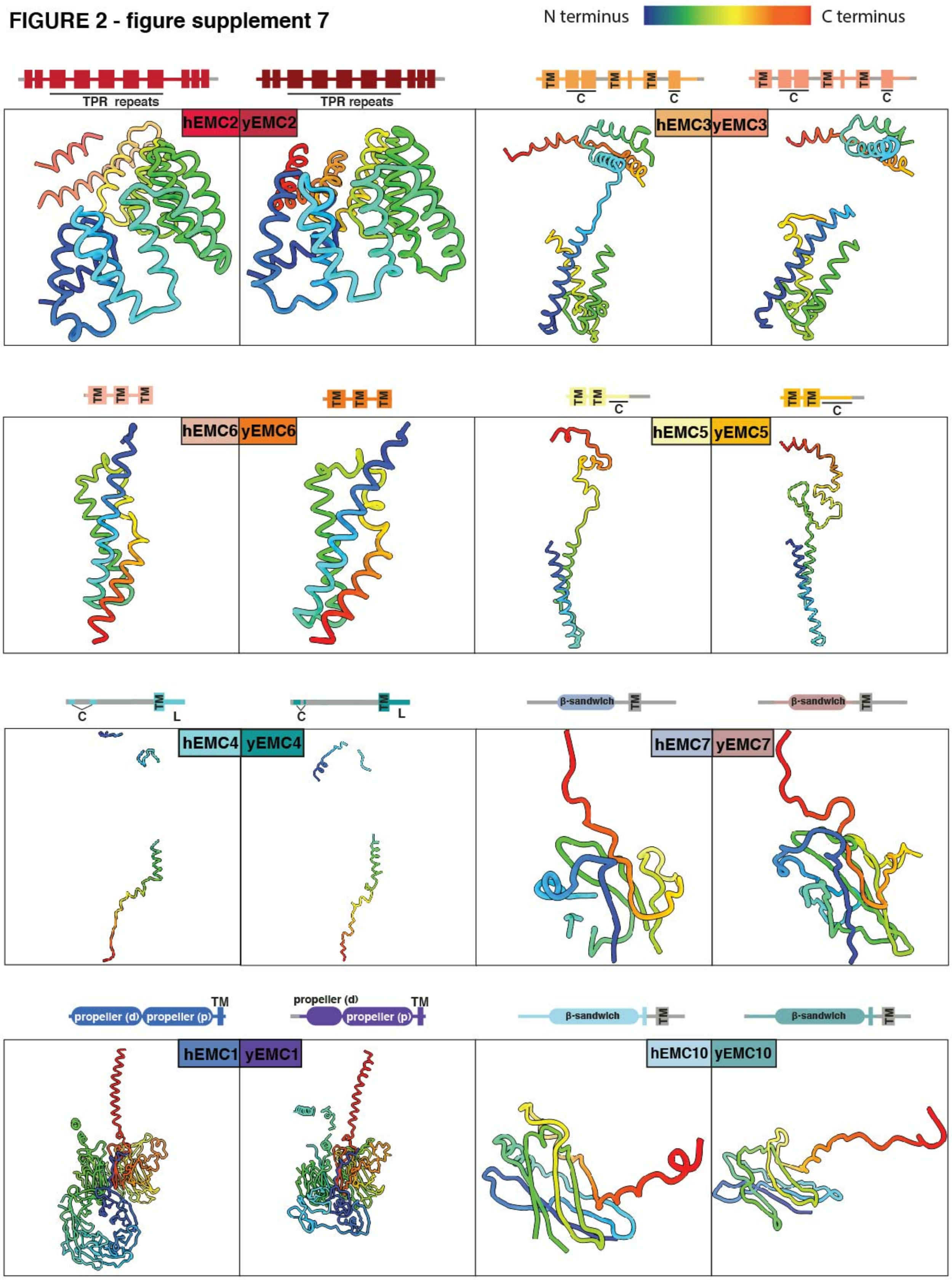

Figure 2 – figure supplement 7: **Comparison of yeast and human EMC subunits.**

#### FIGURE 3 - figure supplement 1

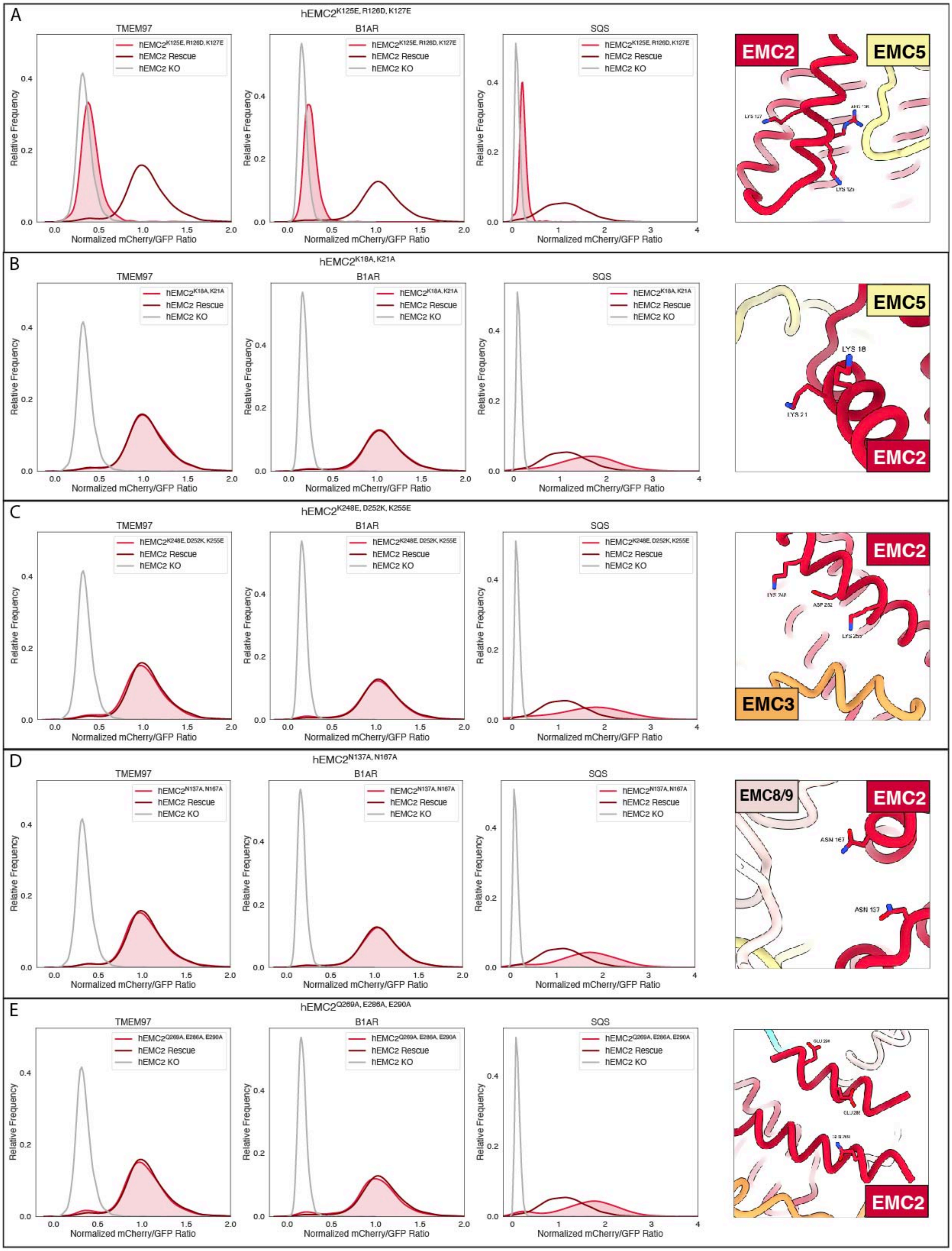

(Legend on next page)

Figure 3 – figure supplement 1: **Flow cytometry for mutations in the EMC cytoplasmic domain.**

a) Mutant hEMC2$^{K125E+R126D+K127E}$ with TMEM97-mCherry, B1AR-mCherry, and mCherry-SQScterm cell lines. Image of hEMC ND model displaying the residues mutated.
b) Mutant hEMC2$^{K18A+K21A}$ with TMEM97-mCherry, B1AR-mCherry, and mCherry-SQScterm cell lines. Snapshot of hEMC ND model displaying the residues mutated.
c) Mutant hEMC2$^{K248E+D252K+K255E}$ with TMEM97-mCherry, B1AR-mCherry, and mCherry-SQScterm cell lines. Snapshot of hEMC ND model displaying the residues mutated.
d) Mutant hEMC2$^{N137A+N167A}$ with TMEM97-mCherry, B1AR-mCherry, and mCherry-SQScterm cell lines. Snapshot of hEMC ND model displaying the residues mutated.
e) Mutant hEMC2$^{Q269A+E286A+E290A}$ with TMEM97-mCherry, B1AR-mCherry, and mCherry-SQScterm cell lines. Snapshot of hEMC ND model displaying the residues mutated.

**FIGURE 3 - figure supplement 2**

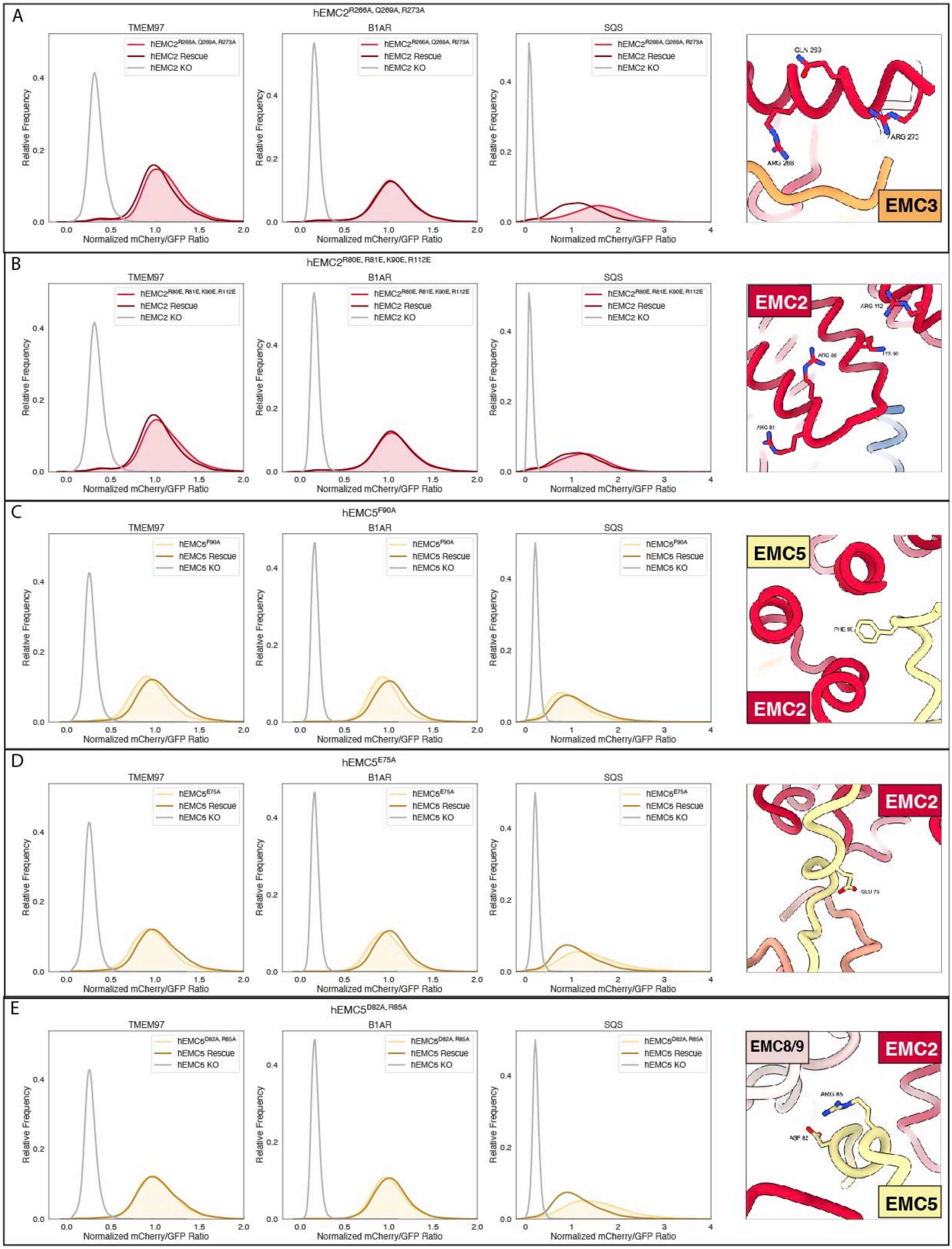

(Legend on next page)

Figure 3 – figure supplement 2: **Additional Flow cytometry for mutations in the EMC cytoplasmic domain.**

a) Mutant hEMC2$^{R266A+Q269A+R273A}$ with TMEM97-mCherry, B1AR-mCherry, and mCherry-SQScterm cell lines. Snapshot of hEMC ND model displaying the residues mutated.
b) Mutant hEMC2$^{R80E+R81E+K90E+R112E}$ with TMEM97-mCherry, B1AR-mCherry, and mCherry-SQScterm cell lines. Snapshot of hEMC ND model displaying the residues mutated.
c) Mutant hEMC5$^{F90A}$ with TMEM97-mCherry, B1AR-mCherry, and mCherry-SQScterm cell lines. Snapshot of hEMC ND model displaying the residues mutated.
d) Mutant hEMC5$^{E75A}$ with TMEM97-mCherry, B1AR-mCherry, and mCherry-SQScterm cell lines. Snapshot of hEMC ND model displaying the residues mutated.
e) Mutant hEMC5$^{D82A+R85A}$ with TMEM97-mCherry, B1AR-mCherry, and mCherry-SQScterm cell lines. Snapshot of hEMC ND model displaying the residues mutated.

#### FIGURE 3 - figure supplement 3

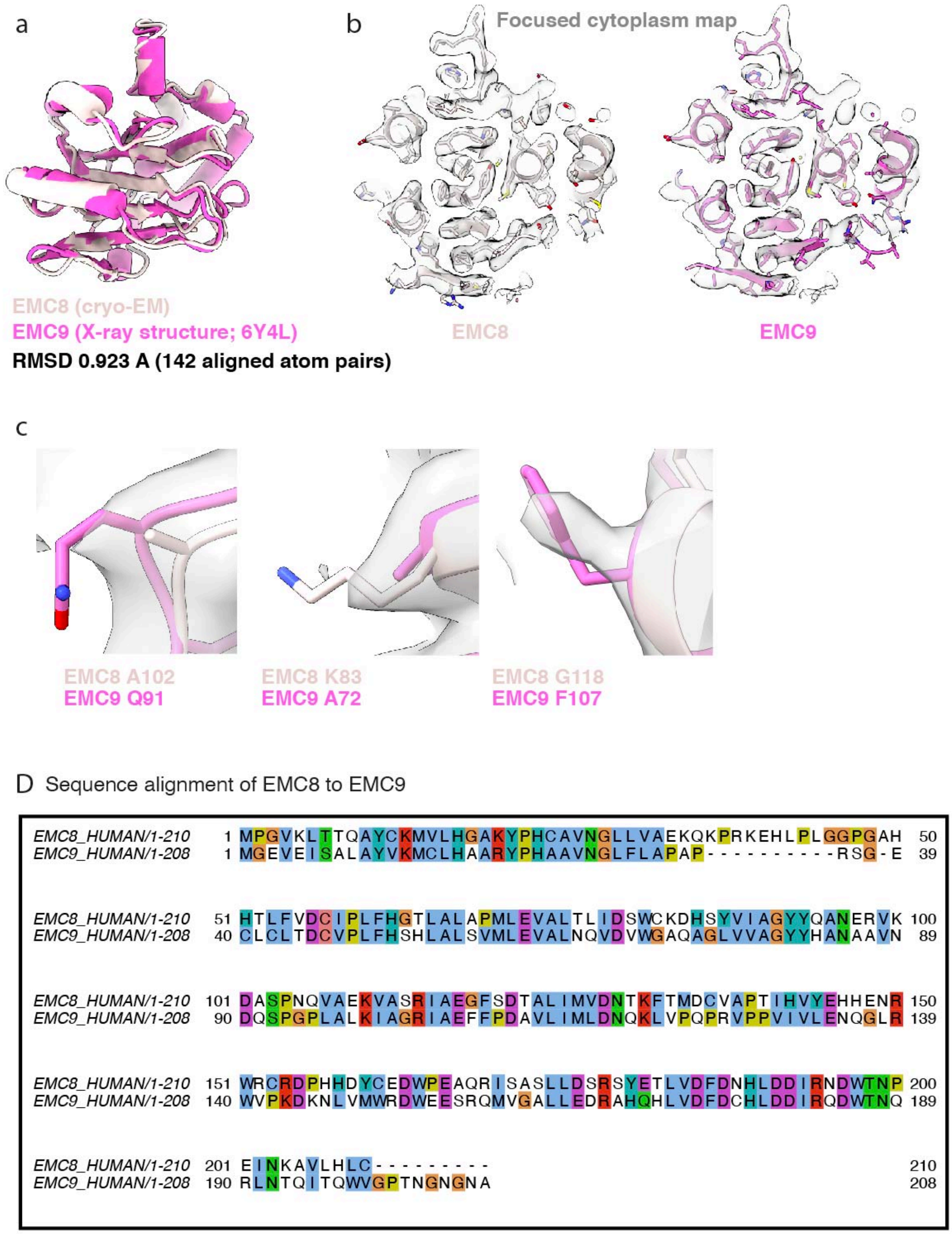

Figure 3 – figure supplement 3: **Both EMC8 and EMC9 can be fitted into the hEMC cryo-EM maps.**

a) Superposition of EMC8 (cryo-EM model, this work) and EMC9 (X-ray model, PDB code 6Y4L) reveals strong structural homology between the two paralogous hEMC subunits.
b) Central slide through the models of EMC8 (cryo-EM) and EMC9 (X-ray) fitted into the hEMC nanodisc cytoplasmic focused map.
c) Representative non-conserved residues in EMC8 and EMC9 showing evidence of side-chain density superposition. The hEMC nanodisc cytoplasmic focused map is depicted.
d) Tcoffee sequence alignment between hEMC8 and hEMC9, displayed in Jalview and ClustalX coloring.

#### FIGURE 5 - figure supplement 1

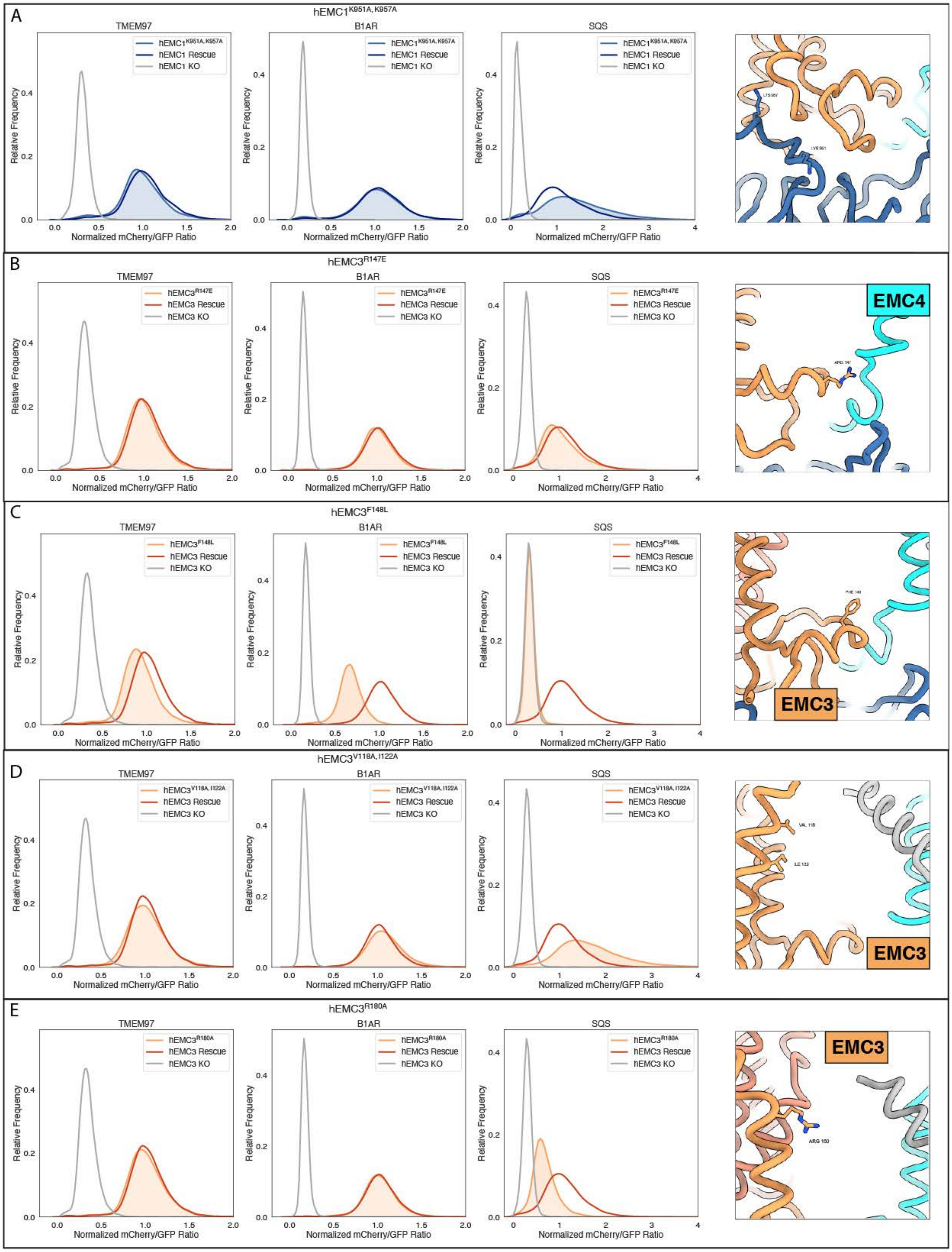

(Legend on next page)

Figure 5 – figure supplement 1: **Flow cytometry and western blot of gated cavity mutants**

a) Mutant hEMC1$^{K951A+K957A}$ with TMEM97-mCherry, B1AR-mCherry, and mCherry-SQScterm cell lines. Snapshot of hEMC ND model displaying the residues mutated.
b) Mutant hEMC3$^{R147E}$ with TMEM97-mCherry, B1AR-mCherry, and mCherry-SQScterm cell lines. Snapshot of hEMC ND model displaying the residues mutated.
c) Mutant hEMC3$^{F148L}$ with TMEM97-mCherry, B1AR-mCherry, and mCherry-SQScterm cell lines. Snapshot of hEMC ND model displaying the residues mutated.
d) Mutant hEMC3$^{V118A+I122A}$ with TMEM97-mCherry, B1AR-mCherry, and mCherry-SQScterm cell lines. Snapshot of hEMC ND model displaying the residues mutated.
e) Mutant hEMC3$^{R180A}$ with TMEM97-mCherry, B1AR-mCherry, and mCherry-SQScterm cell lines. Snapshot of hEMC ND model displaying the residues mutated.

### FIGURE 5 - figure supplement 2

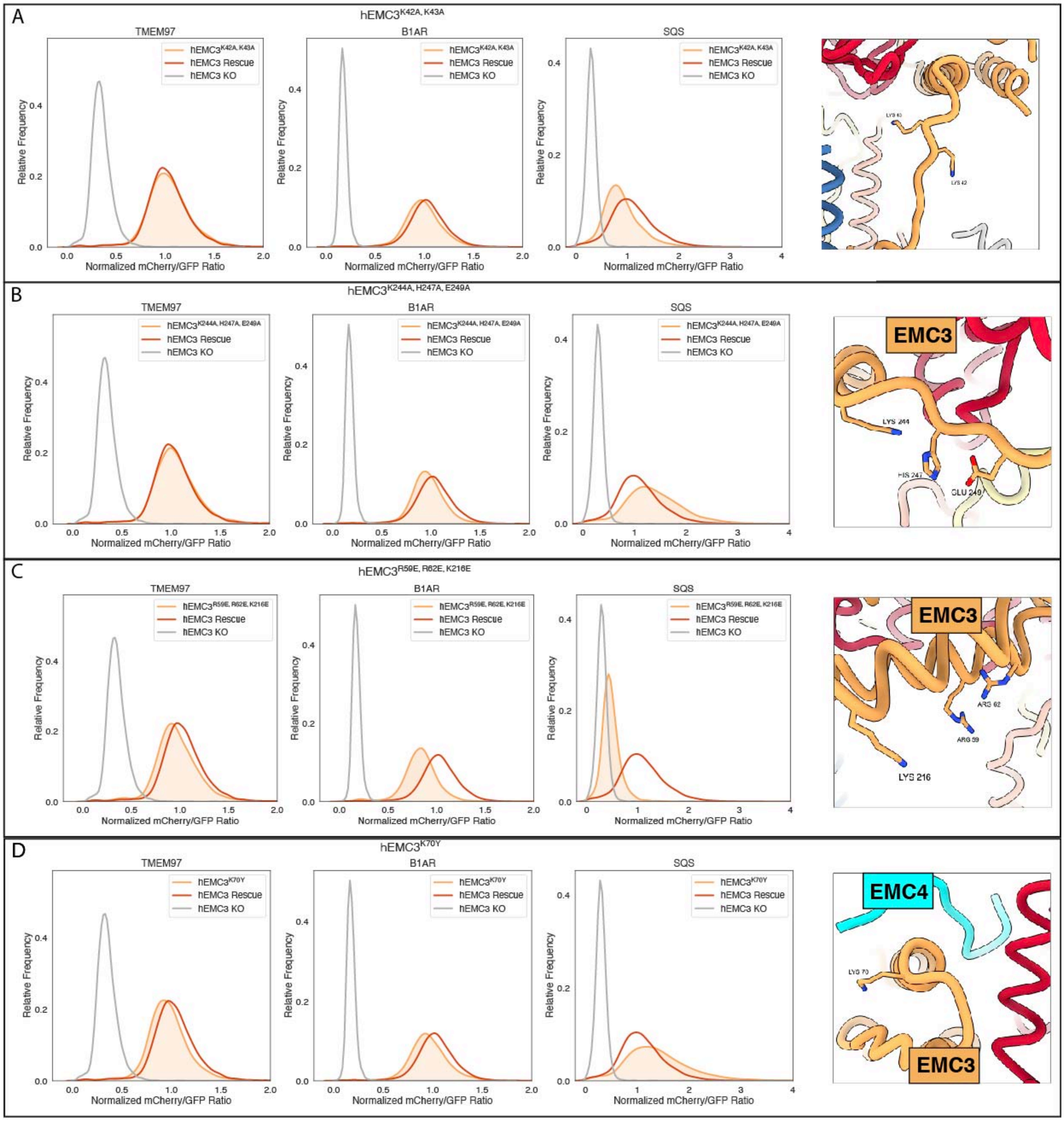

(Legend on next page)

Figure 5 – figure supplement 2: **Additional flow cytometry of gated cavity mutants.**

a) Mutant hEMC3$^{K42A+K43A}$ with TMEM97-mCherry, B1AR-mCherry, and mCherry-SQScterm cell lines. Snapshot of hEMC ND model displaying the residues mutated.
b) Mutant hEMC3$^{K244A+H247A+E249A}$ with TMEM97-mCherry, B1AR-mCherry, and mCherry-SQScterm cell lines. Snapshot of hEMC ND model displaying the residues mutated.
c) Mutant hEMC3$^{R59E+R62E+K216E}$ with TMEM97-mCherry, B1AR-mCherry, and mCherry-SQScterm cell lines. Snapshot of hEMC ND model displaying the residues mutated.
d) Mutant hEMC3$^{K70Y}$ with TMEM97-mCherry, B1AR-mCherry, and mCherry-SQScterm cell lines. Snapshot of hEMC ND model displaying the residues mutated.

#### FIGURE 5 - figure supplement 3

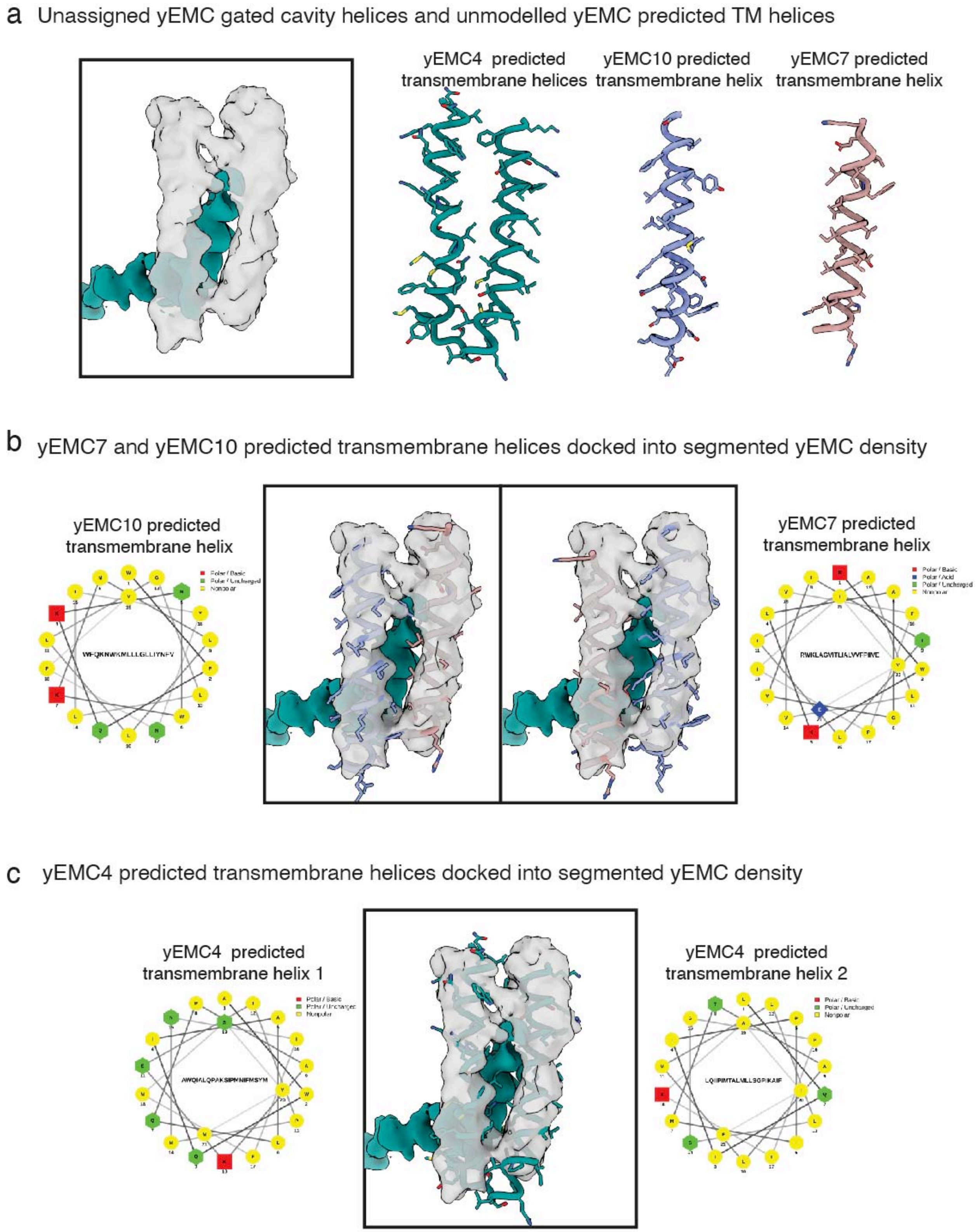

Figure 5 – figure supplement 3: **Ambiguity in assignment of flexible gate helices.**

### FIGURE 5 - figure supplement 4

a Structure comparison of H. sapiens EMC3, S. cerevisiae EMC3, E. coli YidC, and M. jannaschii YidC-like proteins.

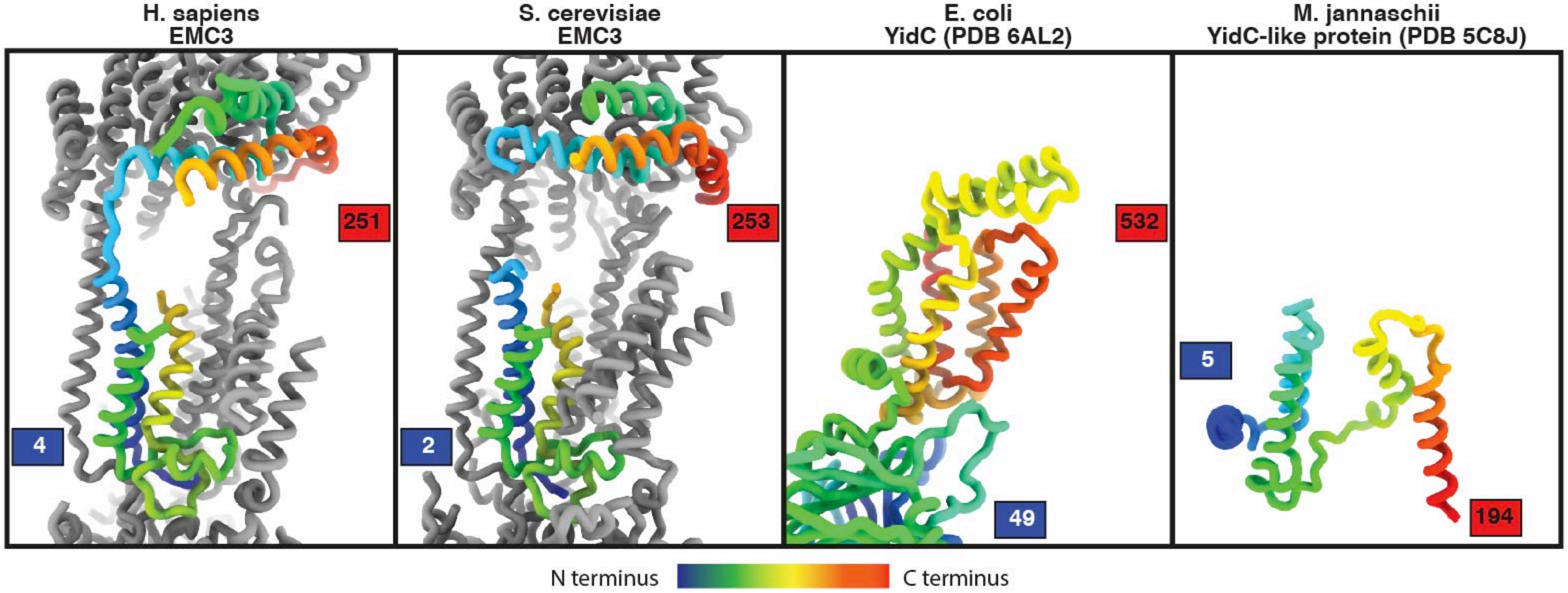

b Sequence alignment of H. sapiens EMC3, S. cerevisiae EMC3, E. coli YidC, and M. jannaschii YidC-like protein colored by percent identity

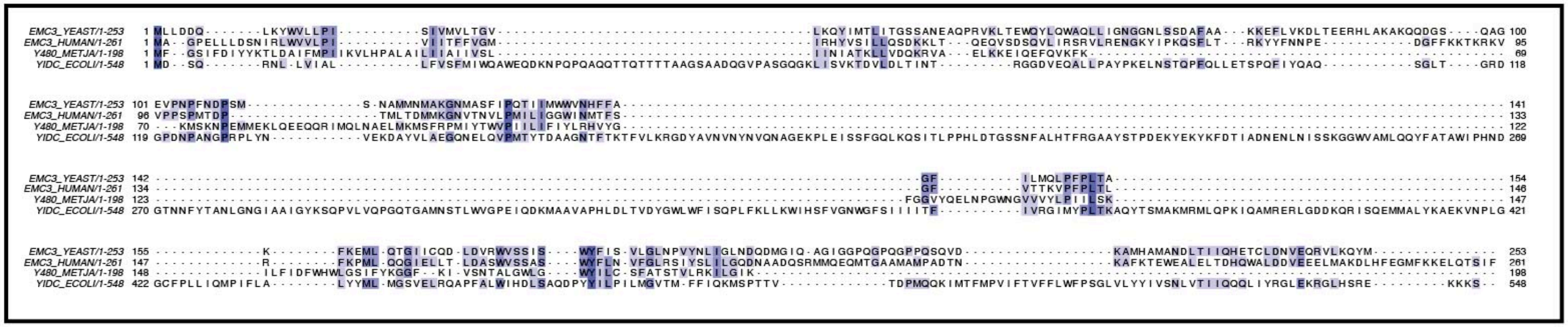

Figure 5 – figure supplement 4: **Comparison of EMC3 to YidC-family members.**

#### FIGURE 6 - figure supplement 1

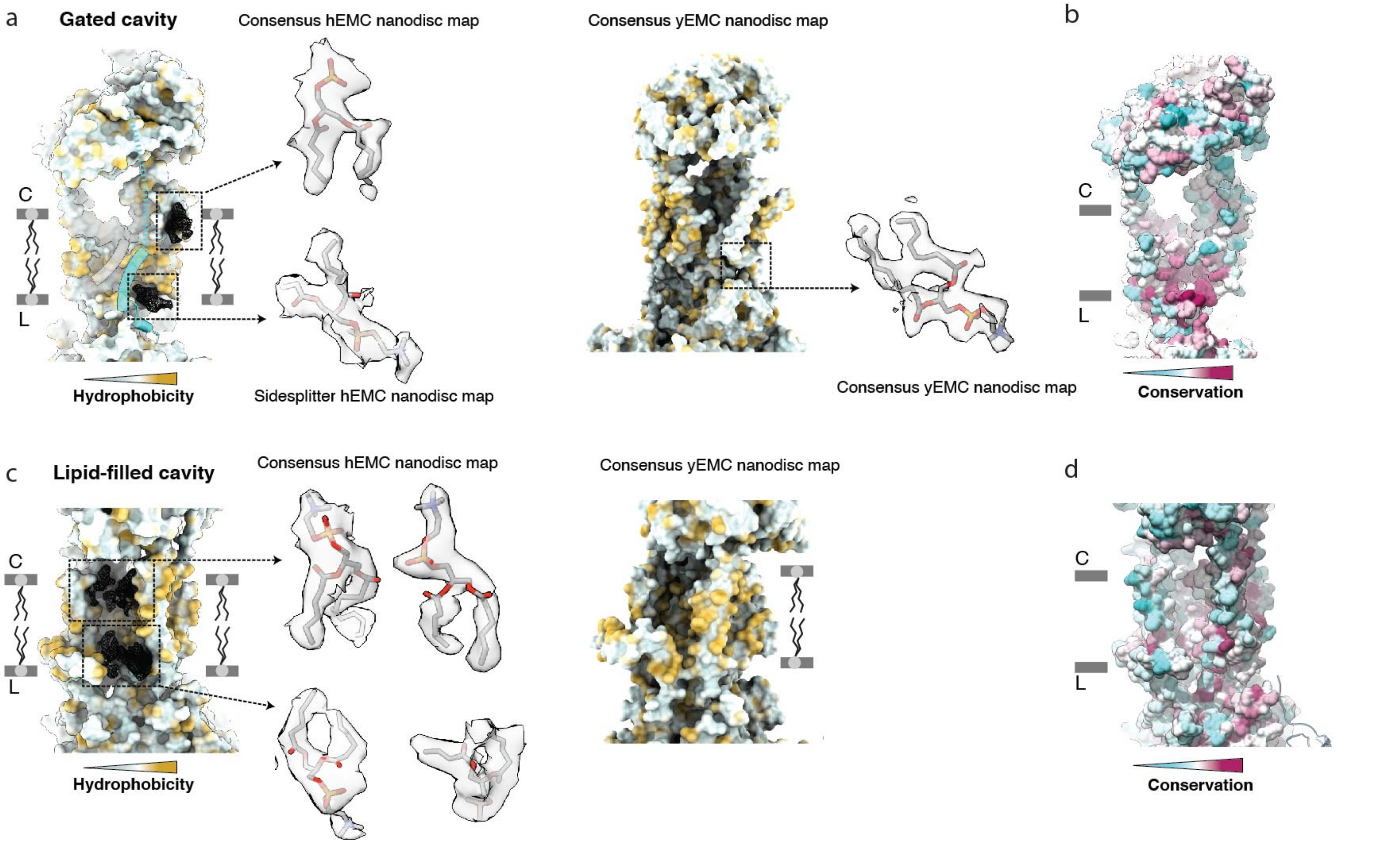

<u>Figure 6 – figure supplement 1:</u> **Resolved lipid densities in hEMC and yEMC nanodisc maps.**

a) Resolved lipids in the hEMC gated cavity. Left: The hEMC model is shown as surface rendering, colored by molecular lipophilicity potential (mint = hydrophilic, gold = hydrophobic). Black mesh corresponds to the hEMC consensus nanodisc map zoned within 3 Å of the modeled POPC molecules. Middle: Enlarged views of lipid densities. Note that the second lipid on the lumenal side of the cavity is shown with the corresponding zoned Sidesplitter hEMC nanodisc map. Since this particular lipid molecule shows only weak features in the consensus map, it is left unmodeled in the hEMC nanodisc model. Right: The yEMC model is shown as surface rendering, colored by molecular lipophilicity potential. One POPC lipid is modeled at the lumenal side of the yEMC gated cavity and shown to the right is the yEMC consensus nanodisc map zoned within 3 Å of a modeled POPC molecule.

b) Same view as in a), with the hEMC model molecular surface colored according to amino acid conservation.

c) As for a) but for the hEMC lipid-filled cavity.

d) Same view as in c), with the hEMC model molecular surface colored according to amino acid conservation.

### FIGURE 6 - figure supplement 2

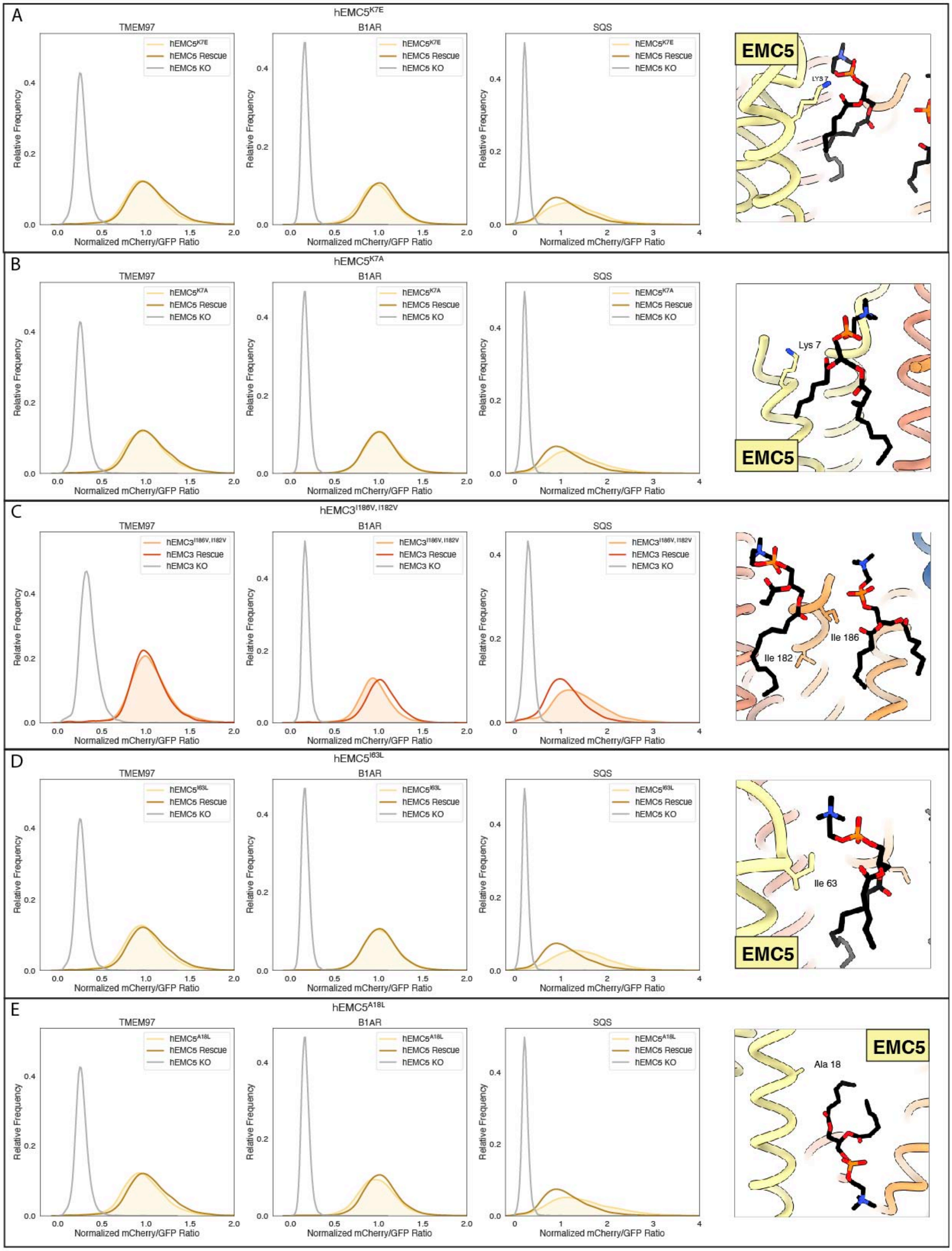

(Legend on next page)

Figure 6 – figure supplement 2: **Flow cytometry of lipid-filled cavity mutants.**

a) Mutant hEMC5$^{K7E}$ with TMEM97-mCherry, B1AR-mCherry, and mCherry-SQScterm cell lines. Snapshot of hEMC ND model displaying the residues mutated.

b) Mutant hEMC5$^{K7A}$ with TMEM97-mCherry, B1AR-mCherry, and mCherry-SQScterm cell lines. Snapshot of hEMC ND model displaying the residues mutated.

c) Mutant hEMC3$^{I182V+I186V}$ with TMEM97-mCherry, B1AR-mCherry, and mCherry-SQScterm cell lines. Snapshot of hEMC ND model displaying the residues mutated.

d) Mutant hEMC5$^{I63L}$ with TMEM97-mCherry, B1AR-mCherry, and mCherry-SQScterm cell lines. Snapshot of hEMC ND model displaying the residues mutated.

e) Mutant hEMC5$^{A18L}$ with TMEM97-mCherry, B1AR-mCherry, and mCherry-SQScterm cell lines. Snapshot of hEMC ND model displaying the residues mutated.

### FIGURE 6 - figure supplement 3

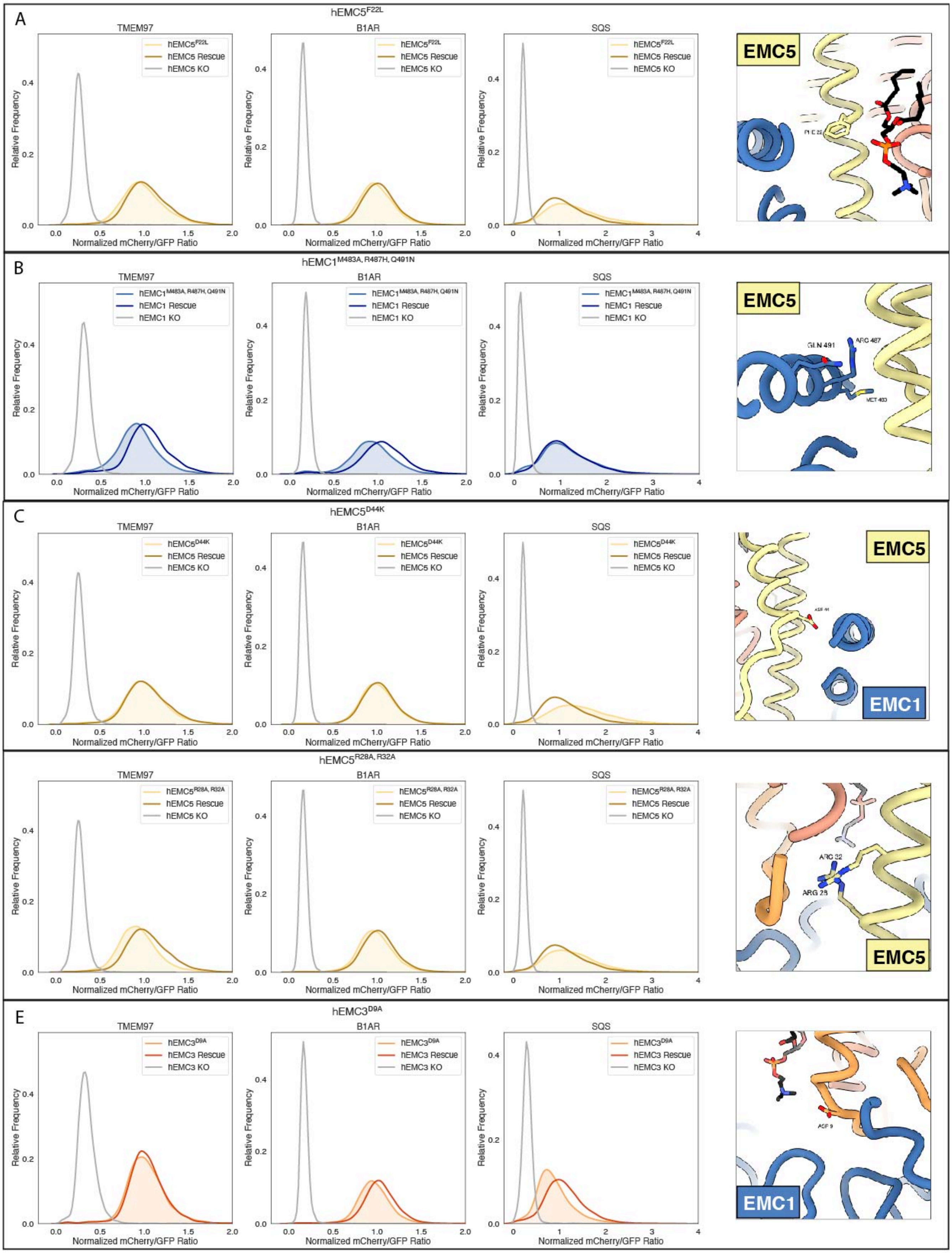

(Legend on next page)

Figure 6 – figure supplement 3: **Additional flow cytometry of lipid-filled cavity mutants.**

a) Mutant hEMC5$^{F22L}$ with TMEM97-mCherry, B1AR-mCherry, and mCherry-SQScterm cell lines. Snapshot of hEMC ND model displaying the residues mutated.
b) Mutant hEMC1$^{M483A+R487H+Q491N}$ with TMEM97-mCherry, B1AR-mCherry, and mCherry-SQScterm cell lines. Snapshot of hEMC ND model displaying the residues mutated.
c) Mutant hEMC5$^{D44K}$ with TMEM97-mCherry, B1AR-mCherry, and mCherry-SQScterm cell lines. Snapshot of hEMC ND model displaying the residues mutated.
d) Mutant hEMC5$^{R28A+R32A}$ with TMEM97-mCherry, B1AR-mCherry, and mCherry-SQScterm cell lines. Snapshot of hEMC ND model displaying the residues mutated.
e) Mutant hEMC3$^{D9A}$ with TMEM97-mCherry, B1AR-mCherry, and mCherry-SQScterm cell lines. Snapshot of hEMC ND model displaying the residues mutated.

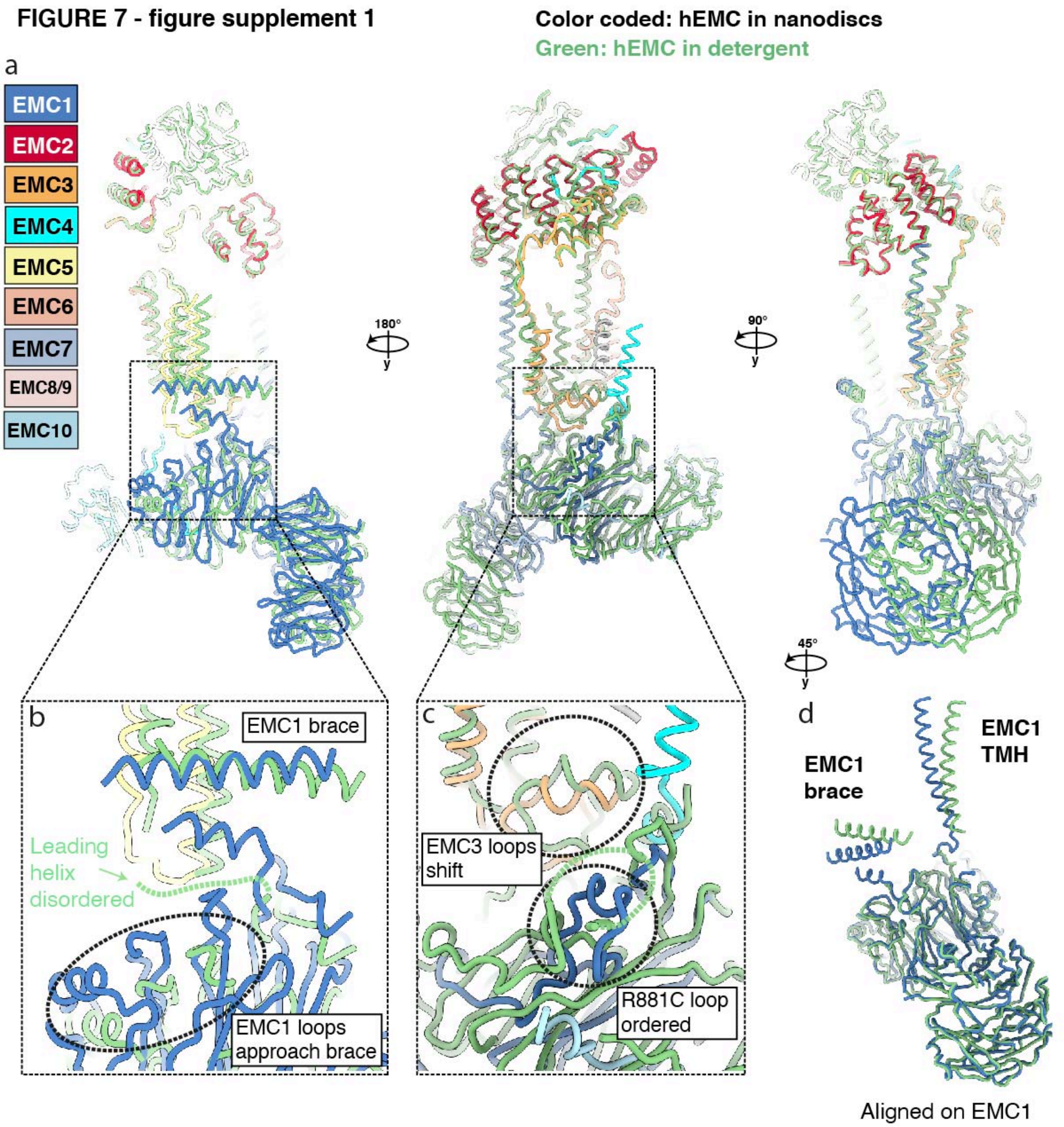

Figure 7 – figure supplement 1: **Conformational heterogeneity of the hEMC lumenal domain between detergent and nanodisc maps.**

a) Superposition of hEMC in detergent (green) and nanodiscs (color-coded). Models were aligned on EMC2 (cytoplasm) and EMC5 (transmembrane), revealing a rotation of the lumenal domain.
b) Enlarged view on the superposition around the area of the EMC1 brace and the membrane proximal beta-propeller. A helix leading towards the amphipathic brace is resolved in the nanodisc map and disordered in the detergent map. In the nanodisc structure, several proximal propeller loops approach and form stabilizing interactions with the leading helix (dashed circle).
c) Enlarged view on the superposition around the area of the lumenal hydrophobic seal of the gated cavity. The loop stabilized by EMC1 R881 (corresponding to the disease mutant R881C) is well resolved in the hEMC nanodisc map and more disordered in the detergent map. The loops of the EMC3 hydrophobic seal shift concomitantly with the lumenal rotation between detergent and nanodisc structures.
d) Structural alignment of EMC1 from the detergent (green) and nanodisc (blue) models. While the beta-propellers align extremely well, a clear rotation of the EMC1 brace and the EMC1 transmembrane helix can be observed.

#### FIGURE 7 - figure supplement 2

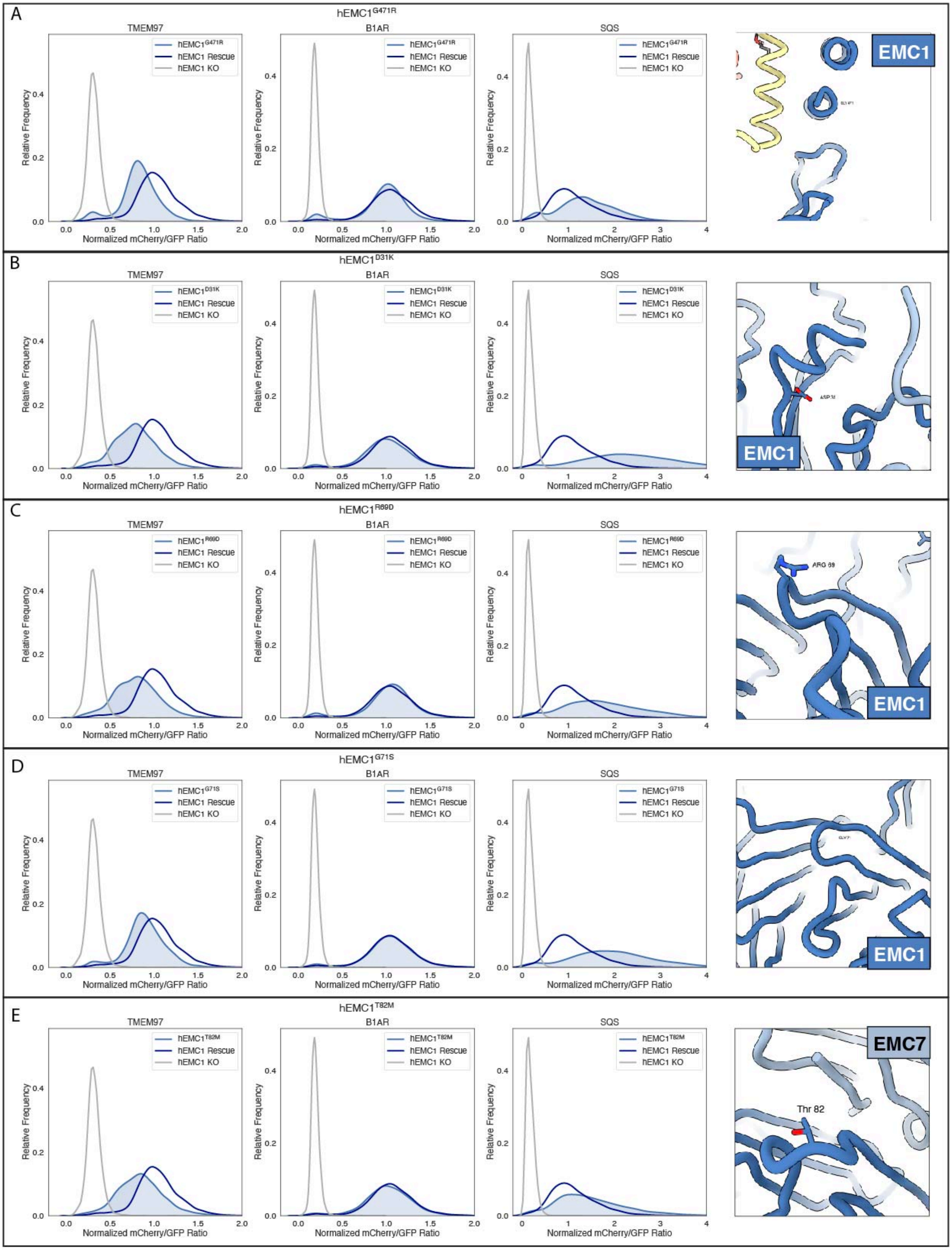

(Legend on next page)

Figure 7 – figure supplement 2: **Flow cytometry of lumenal domain mutants.**

a) Mutant hEMC1$^{G471R}$ with TMEM97-mCherry, B1AR-mCherry, and mCherry-SQScterm cell lines. Snapshot of hEMC ND model displaying the residues mutated.
b) Mutant hEMC1$^{D31K}$ with TMEM97-mCherry, B1AR-mCherry, and mCherry-SQScterm cell lines. Snapshot of hEMC ND model displaying the residues mutated.
c) Mutant hEMC1$^{R69D}$ with TMEM97-mCherry, B1AR-mCherry, and mCherry-SQScterm cell lines. Snapshot of hEMC ND model displaying the residues mutated.
d) Mutant hEMC1$^{G71S}$ with TMEM97-mCherry, B1AR-mCherry, and mCherry-SQScterm cell lines. Snapshot of hEMC ND model displaying the residues mutated.
e) Mutant hEMC1$^{T82M}$ with TMEM97-mCherry, B1AR-mCherry, and mCherry-SQScterm cell lines. Snapshot of hEMC ND model displaying the residues mutated.

#### FIGURE 7 - figure supplement 3

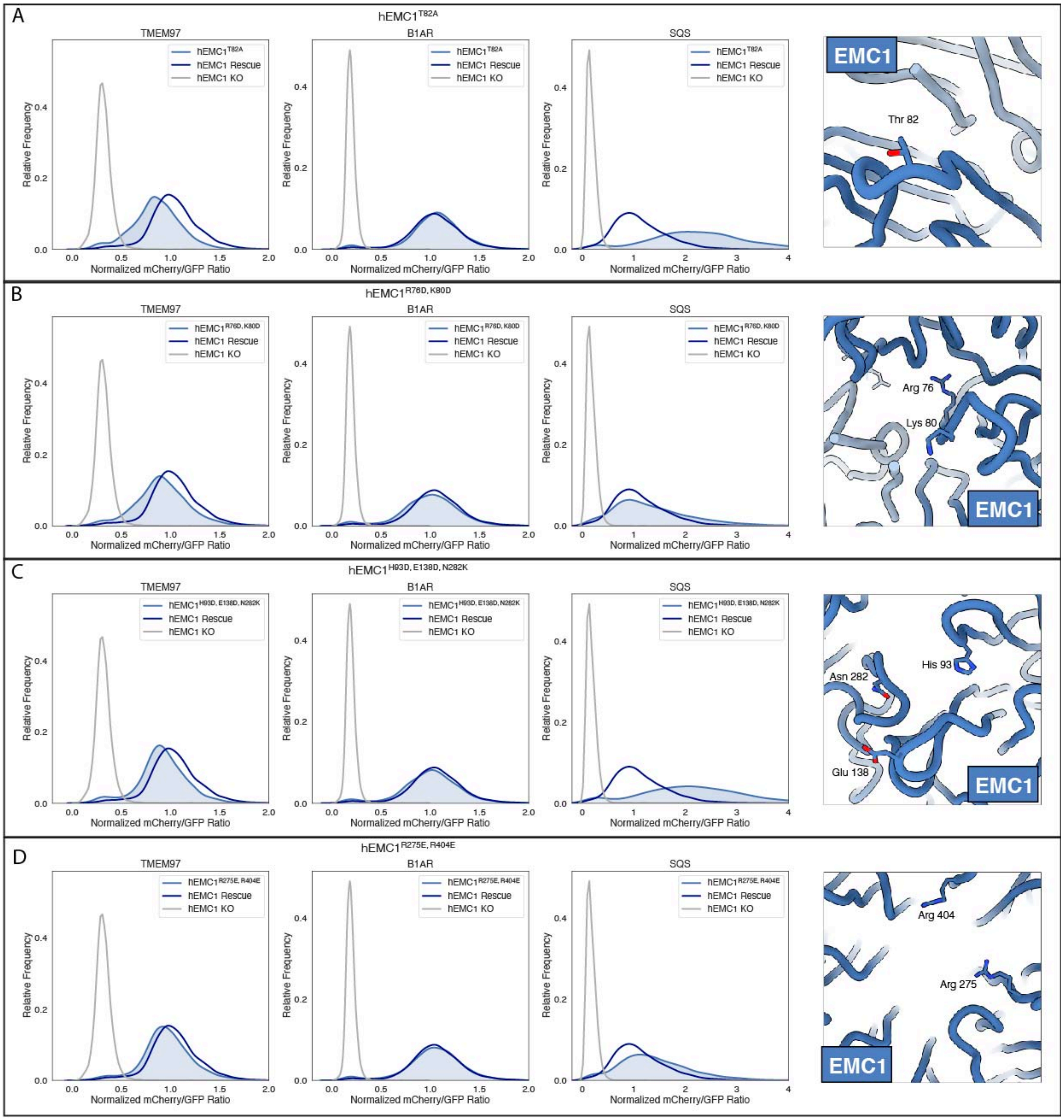

(Legend on next page)

Figure 7 – figure supplement 3: **Additional flow cytometry of lumenal domain mutants.**

a) Mutant hEMC1$^{T82A}$ with TMEM97-mCherry, B1AR-mCherry, and mCherry-SQScterm cell lines. Snapshot of hEMC ND model displaying the residues mutated.
b) Mutant hEMC1$^{R76D+K80D}$ with TMEM97-mCherry, B1AR-mCherry, and mCherry-SQScterm cell lines. Snapshot of hEMC ND model displaying the residues mutated.
c) Mutant hEMC1$^{H93D+E138D+N282K}$ with TMEM97-mCherry, B1AR-mCherry, and mCherry-SQScterm cell lines. Snapshot of hEMC ND model displaying the residues mutated.
d) Mutant hEMC1$^{R275E+R404E}$ with TMEM97-mCherry, B1AR-mCherry, and mCherry-SQScterm cell lines. Snapshot of hEMC ND model displaying the residues mutated.

| | yEMC in detergent dataset 1 | yEMC in detergent dataset 2 | yEMC in nanodiscs | hEMC in detergent | hEMC in nanodiscs |
|---|---|---|---|---|---|
| **Data collection and processing** | | | | | |
| Microscope | Technai Polara | FEI Titan Krios | FEI Titan Krios | FEI Titan Krios | FEI Titan Krios |
| Camera | Gatan K2 Summit | Gatan K2 Summit | Gatan K3 | Gatan K3 | Gatan K3 |
| Magnification | 31,000x | 22,500x | 105,000x | 81,000x | 105,000x |
| Voltage (kV) | 300 | 300 | 300 | 300 | 300 |
| Electron exposure ($e^{-}/Å^{2}$) | 56.8 | 58.3 | 67 | 62 | 72 |
| Defocus range (μm) | -1.0 to -3.0 | -1.0 to -3.0 | -0.8 to -2.5 | -0.7 to - 2.8 | -0.7 to - 2.8 |
| Pixel size (Å) | 1.22 | 1.31 | 0.853 | 1.094 | 0.8512 |
| Software | Relion 2.0, Relion 3.0, THUNDER | Relion 2.0, Relion 3.0, THUNDER | Relion 3.0, cryoSPARC v2 | Relion 3.0, cryoSPARC v2 | Relion 3.0, cryoSPARC v2 |
| Symmetry imposed | C1 | C1 | C1 | C1 | C1 |
| Initial particle images (no.) | 419,907 | 670,078 | 6,100,000 | 3,350,000 | 5,900,000 |
| Final particle images (no.) | 83,599 | 170,186 | 230,528 | 144,222 | 177,560 |
| Overall map resolution (Å) | 8 | 7 | | | |
| FSC threshold 0.143 | 4.3 (combined) | | 3.2 | 3.60 | 3.39 |
| Local map resolution range (Å) | 3.6-6.4 | | 2.6-6.4 | 2.8 – 6.0 | 3.0 – 7.2 |
| **Refinement** | | | | | |
| Software | | | Phenix 1.18 real-space-refine | Phenix 1.18 real-space-refine | Phenix 1.18 real-space-refine |
| Model resolution (Å) | | | | | |
| FSC threshold 0.5 | | | 3.5 | 3.9 | 3.6 |
| Map sharpening B factor ($Å^{2}$) | | | -75 | -115 | -126 |
| Model composition | | | | | |
| Non-hydrogen atoms | | | 17,316 | 15,012 | 16,834 |
| Protein residues | | | 2168 | 1880 | 2085 |
| Ligands | | | NAG: 5; PCW: 1 | NAG: 2 | NAG: 4; PCW: 5 |
| B factors ($Å^{2}$) | | | | | |
| Protein (mean) | | | 103 | 111 | 126 |
| Ligand (mean) | | | 88 | 107 | 127 |
| R.m.s. deviations | | | | | |
| Bond lengths (Å) | | | 0.005 | 0.005 | 0.006 |
| Bond angles (°) | | | 0.923 | 0.608 | 0.666 |
| Validation | | | | | |
| MolProbity score | | | 1.42 | 1.56 | 1.40 |
| Clashscore | | | 3.9 | 9 | 6 |
| Ramachandran plot | | | | | |
| Favored (%) | | | 96.3 | 96.1 | 97.6 |
| Allowed (%) | | | 3.7 | 3.9 | 2.4 |
| Disallowed (%) | | | 0 | 0 | 0 |

Table 1: **Cryo-EM data acquisition, reconstruction and model refinement statistics.**
